# Developmental origins and evolution of pallial cell types and structures in birds

**DOI:** 10.1101/2024.04.30.591857

**Authors:** Bastienne Zaremba, Amir Fallahshahroudi, Céline Schneider, Julia Schmidt, Ioannis Sarropoulos, Evgeny Leushkin, Bianka Berki, Enya Van Poucke, Per Jensen, Rodrigo Senovilla-Ganzo, Francisca Hervas-Sotomayor, Nils Trost, Francesco Lamanna, Mari Sepp, Fernando García-Moreno, Henrik Kaessmann

**Affiliations:** Center for Molecular Biology (ZMBH), DKFZ-ZMBH Alliance, Heidelberg University, Heidelberg, Germany; Department of Medical Biochemistry and Microbiology, Biomedical Center (BMC), Uppsala University, Uppsala, Sweden; Wellcome Sanger Institute, Wellcome Genome Campus, _Hinxton_, Cambridge, UK; LOEWE Centre for Translational Biodiversity Genomics, _60325_ Frankfurt, Germany; Deep Sequencing Core Facility, CellNetworks Excellence Cluster, Heidelberg University, _Im Neuenheimer Feld 267_, _D-69120_, Heidelberg, Germany; IFM Biology, AVIAN Behavioural Genomics and Physiology group, Linköping University, Sweden; Achucarro Basque Center for Neuroscience, Scientific Park of the University of the Basque Country (UPV/EHU), _Leioa_, Spain; INRAE, LPGP, _Rennes_, France; Department of Neuroscience, Faculty of Medicine and Odontology, UPV/EHU, _Leioa_, Spain; IKERBASQUE Foundation, Bilbao, Spain

## Abstract

The advanced cognitive abilities of birds rival those of mammals and have been attributed to evolutionary innovations in the pallium. However, a comprehensive cellular characterization of this brain region in birds has been lacking. We scrutinized the structures, cell types and evolutionary origins of the avian pallium based on single-cell and spatial transcriptomics atlases for the adult and developing chicken, and comparisons to corresponding data from mammals and non-avian reptiles. We found that the avian pallium shares most inhibitory neuron types with other amniotes. While excitatory neuron repertoires in the (medial) hippocampal formation show high conservation, they substantially diverged in other pallial regions during avian evolution, defining novel structures like the avian-specific (dorsal) hyperpallium, whose neuronal gene expression identities partly converge during late development with those of the (ventral) nidopallium. Our work also unveils the evolutionary relationships of pallial structures across amniotes, like the previously unknown homology between avian (lateral) mesopallial and mammalian deep layer cortical neurons.

**One-Sentence Summary:** An avian neural cell type atlas illuminates the developmental origins and evolution of the amniote pallium.

---

Some bird species rival non-human primates in their intelligence (*1*). This observation has been attributed to an increase in relative brain size and cell density, and the presence of circuitries in the avian pallium akin to those of the mammalian isocortex (traditionally called neocortex) (*2*, *3*). The pallium, the equivalent of the dorsal telencephalon, is thought to be the brain region that has undergone the most dramatic morphological changes during the evolution of amniotes (i.e., mammals, birds, and non-avian reptiles) since their last common ancestor ∼320 million years ago (*4*). While in mammals the pallium mainly consists of layered structures, among them the isocortex, which is mostly derived from the dorsal pallium during development, the pallium of birds/reptiles mainly comprises the dorsal ventricular ridge (DVR), which is nuclear in organization and derives from the ventral and lateral pallia (Fig. 1A).

**Fig. 1.**
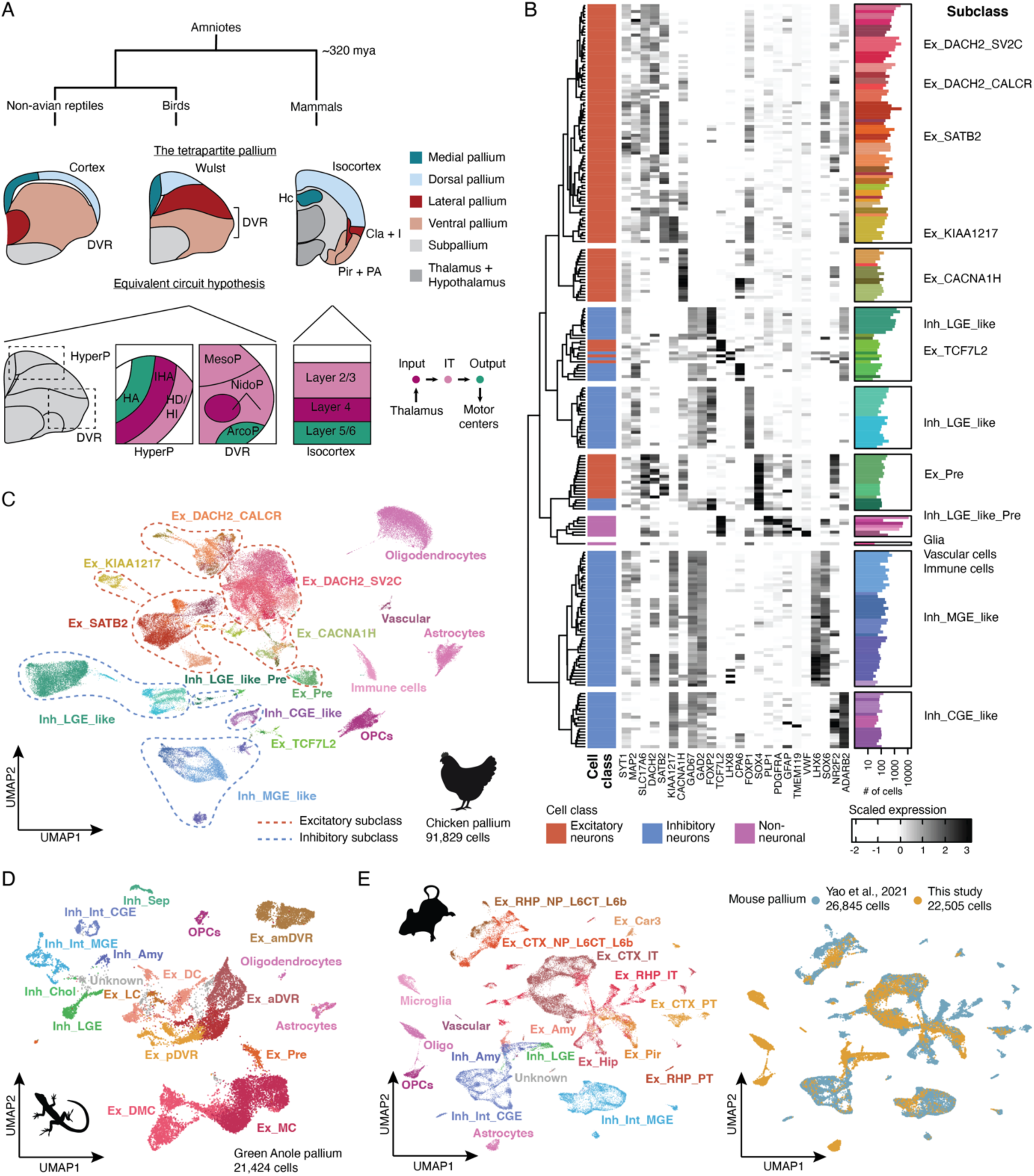
Cell type atlases of the pallium in amniotes. (**A**) Current models of amniote pallium evolution. (Top) Schematic representation of coronal sections of the telencephalon in lizard (left), chicken (middle), and mouse (right). Brightly colored areas represent the pallium divided into developmental, homologous sectors according to the tetrapartite pallium model (*6*). Mya, million years ago; DVR, dorsal ventricular ridge; Hc, hippocampus; Cla, claustrum; I, insular cortex; Pir, piriform cortex; PA, pallial amygdala. (Bottom) Schematic representation of regions constituting sensory circuits in the pallium of birds (left) and mammals (right), colored according to their role in the canonical circuit (illustrated on the far right). Colored areas indicate regions that are homologous according to the equivalent circuit hypothesis, given their comparable circuit functions across birds and mammals. HyperP, hyperpallium; HA, apical hyperpallium; IHA, intercalated hyperpallium; HD/HI, densocellular hyperpallium/intermediate hyperpallium; MesoP; mesopallium; NidoP, nidopallium; ArcoP, arcopallium; IT, intra-telencephalic. (**B**) Heatmap of selected marker gene expression across 237 clusters in snRNA-seq-based chicken pallium atlas, ordered by independently constructed cluster dendrogram. Barplots of cell numbers per cluster on the right are coloured by supertype annotation. (**C**) Uniform manifold approximation and projection (UMAP) of chicken pallium snRNA-seq atlas colored by supertype annotation. Dotted lines and text labels represent subclass annotations. (**D**) UMAP of snRNA-seq based green anole pallium atlas coloured by and labelled with subclass annotation. (**E**) UMAP of snRNA-seq and single-cell RNA sequencing-based mouse pallium atlas, colored by and labelled with subclass annotation (left), or colored according to underlying dataset (right).

The DVR is especially prominent in archosaurs (crocodilians and birds), and in contrast to non-avian reptiles, birds completely lack a layered cortex; instead, birds possess another nuclear structure, called the hyperpallium (encompassing the Wulst and a small caudal region), in its place (Fig. 1A). This anatomical dissimilarity has elicited the emergence of two major opposing views on the evolution of the amniote pallium. One focuses on circuitry and assumes homology for cell types carrying out comparable roles in the conserved circuit (*5*). The other postulates that homologous progenitor domains during development give rise to homologous adult structures, implying that shared circuitry across different pallial domains arose convergently during evolution (*6*). Recent pioneering single-cell molecular studies of the reptilian pallium (*7*) and of small regions in the pallium of songbirds (*8*) lend support to the latter hypothesis. However, cell type level characterizations of the complete avian pallium, and cross-amniote comparisons of this brain region using comparable data, have been lacking.

In this study, we generated a spatially resolved cell type atlases of the entire chicken pallium in adults and across *in ovo* development based on extensive single-nucleus RNA sequencing (snRNA-seq) and spatial transcriptomics datasets, as well as targeted epigenomic data (snATAc-seq). By comparing this atlas to corresponding data from mammals and non-avian reptiles, we trace the structures, cell types and development of the avian pallium as well as its evolutionary relationships with the pallia of the other two amniote lineages.

## A comparative cell type atlas of the adult chicken pallium

To establish a cell type atlas of the adult chicken pallium (excluding olfactory bulb, OB), we profiled broad rostral to caudal sections of this brain region from four individuals, and the complete pallium of one individual, which we specifically dissected into four anatomical regions (DVR excluding arcopallium, arcopallium, Wulst, avian hippocampal region), using snRNA-seq (Methods) (fig. S1 and 2). We obtained high-quality nuclear transcriptomes for 91,829 cells, with a median of 4,808 RNA molecules (unique molecular identifiers, UMIs) and a median of 2,057 transcribed genes detected per cell (fig. S1). To enable the examination of the spatial distribution of cell types in the chicken pallium, we generated in situ sequencing (ISS (*9*)) data for 50 selected marker genes from the complete dataset (Methods). Given that not all supertypes are represented with specific marker genes in these data, we utilized Tangram (*10*) to infer positions based on the collective profile of all 50 mapped genes (Table S5). To complement these targeted spatial data, we additionally generated spatial transcriptomics data using the Visium (10x Genomics) technology (Methods). We then used these data to classify cells into groups at four hierarchical levels: classes, subclasses, supertypes, and clusters. Cells were first assigned to one of three major cell type classes according to the expression of well-established marker genes (Methods): inhibitory (GABAergic, γ-aminobutyric acid-producing) neurons, excitatory (glutamatergic) neurons, and non-neuronal cells (Fig. 1, B and C).

The non-neuronal class included cells belonging to the oligodendrocyte lineage, like oligodendrocyte progenitor cells (*SOX6, PDGFRA*), and mature oligodendrocytes (*PLP1*); astrocytes (*SLC1A3*,*GFAP*); immune cells (*TMEM119)*); and a heterogeneous population of cells associated with vasculature (*VWF*)(Fig. 1B).

Neurons were identified based on the expression of pan-neuronal markers, as well as class-specific genes (Fig. 1B, Table S2). We iteratively clustered inhibitory and excitatory neurons (23,848 cells and 45,425 cells, respectively), thus identifying 109 inhibitory and 120 excitatory cell type ‘clusters’. We then constructed dendrograms for each neuronal class and grouped clusters into ‘subclasses’ and ‘supertypes’ based on their position in the dendrogram and their identities in low resolution Louvain communities (Methods). Overall, we annotated 11 subclasses of differentiated and immature neurons, which were split into 46 supertypes (Fig. 1, B and C) (Table S2). In a common dendrogram, non-neuronal cells, inhibitory, and excitatory neurons mainly cluster separately, except for immature neurons (*SOX4*) and a small subclass (*TCF7L2*), which likely represents cells inadvertently co-dissected from the thalamus (*11*) (Fig. 1B).

To discern potentially homologous cell types across amniotes, it is crucial to survey all pallial regions in each species, ideally employing similar methodology. We thus generated snRNA-seq data for the complete pallium of the green anole lizard (excluding OB), in which the DVR and cortex were profiled separately. The resulting dataset (21,424 cells, median UMIs/genes: 2,269/1,447) was iteratively clustered, as detailed for the chicken data above, and clusters were annotated mainly based on single-cell transcriptomics datasets for another lizard (*12*, *13*) and a turtle (*7*) (Fig. 1D; fig. S3 and 4). To obtain a dataset covering the whole murine pallium for our comparative work, we generated snRNA-seq data (22,505 cells, median UMIs/genes: 6,035,/2,883) for parts of the mouse isocortex and structures deriving from the lateral and ventral pallial sectors (Fig. 1E; fig. S5), which complemented available single-cell RNA-seq data of the murine isocortex and hippocampal formation (*14*). Cell populations unique to our mouse dataset were annotated based on the expression of marker genes identified in single-cell datasets covering smaller regions (*15*, *16*) and the Allen Mouse Brain Atlas data (*17*) (fig. S6). Altogether, we compiled a comprehensive comparative dataset of the amniote pallium, spanning representatives from all three major amniote branches.

## GABAergic inhibitory neurons are mostly conserved across amniotes

Most GABAergic neurons in the amniote pallium originate in the ganglionic eminences in the subpallium (*18*, *19*), a developmental process best described in mammals. To compare inhibitory neurons between birds and mammals, we first sought to characterize and spatially map these neurons in the chicken pallium. Hierarchical clustering of all GABAergic clusters in the chicken pallium revealed three distinct groups, characterized by the expression of known marker genes (Fig. 2A) (Table S2) and likely reflecting their developmental origin the lateral, medial, and caudal ganglionic eminences (LGE, MGE, and CGE) (*8*, *14*). Most GABAergic neurons had either an LGE- or MGE-like identity, and only ∼6% showed a CGE-like profile (Table S2), which is in stark contrast to mammals, where CGE-derived neurons are numerous in the pallium (*14*). Overall, we annotated six LGE-like, eight MGE-like, and three CGE-like supertypes (Fig. 2, A and B; Table S2).

**Fig. 2.**
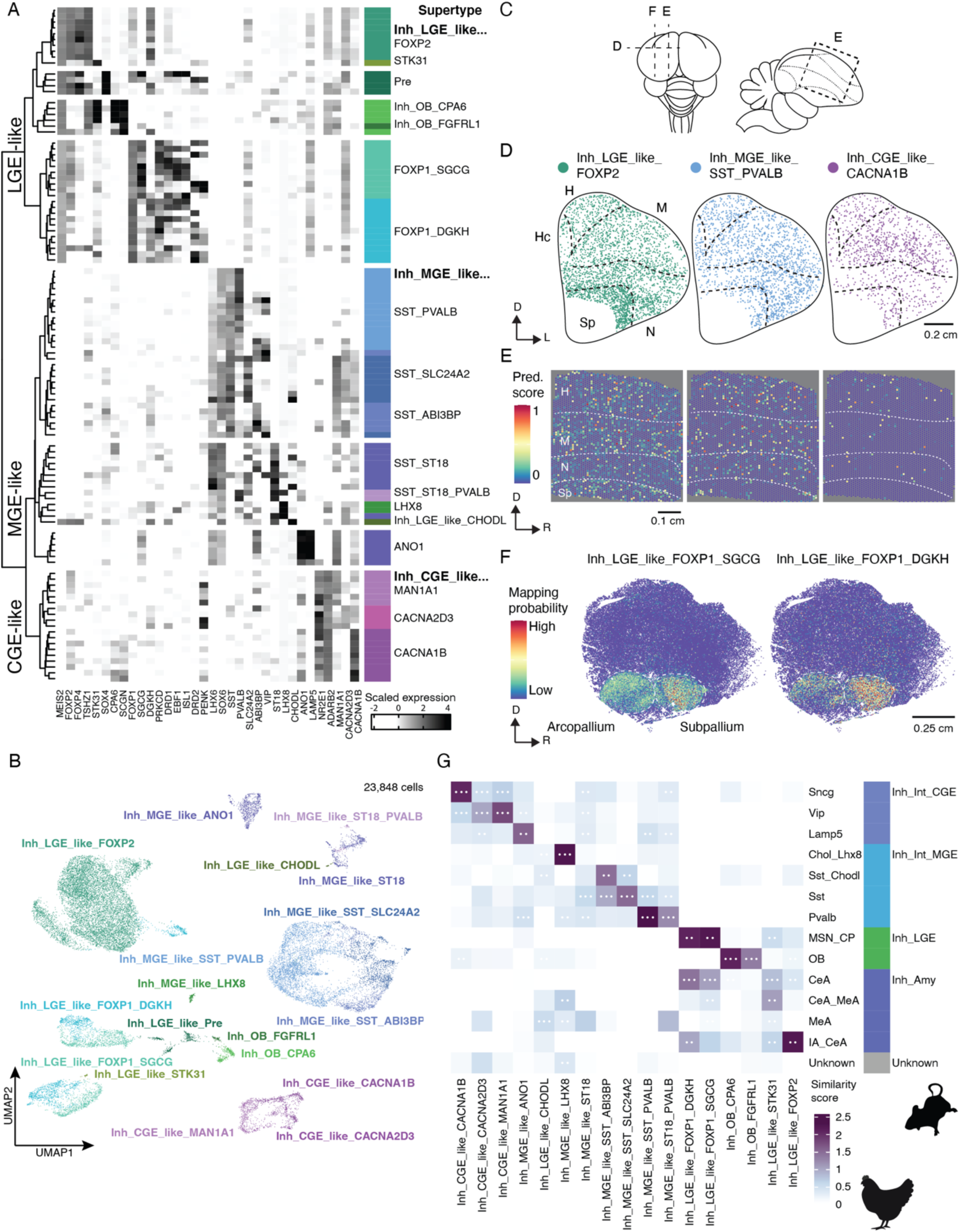
GABAergic inhibitory neurons in the chicken pallium. (**A**) Heatmap of selected marker gene expression across 109 clusters of inhibitory neurons in the chicken pallium ordered according to cluster dendrogram. Color bar and text labels on the right indicate supertype annotation. LGE, lateral ganglionic eminence; MGE, medial ganglionic eminence; CGE, caudal ganglionic eminence. (**B**) UMAP of inhibitory cells colored by and labelled with supertype annotation. (**C**) Schematics of the top view (left) and a midsagittal section (right) of the chicken brain, illustrating positions of tissue sections shown in (D-F). Dotted lines represent borders between pallial brain regions. (**D**) Spatial location of most abundant supertypes from each major inhibitory subclass according to *in situ* sequencing (ISS). Only segmented cells with confidently assigned identity are shown. Hc, hippocampal area; M, mesopallium; N, nidopallium; Sp, subpallium; D, dorsal; l, lateral (**E**) Spatial location of same supertypes as shown in (D) according to Visium data. High prediction (Pred.) scores indicate high probability that cells with the respective identity were present within the spot’s area. R, rostral. (**F**) Spatial location probabilities of both *FOXP1*+ LGE-derived supertypes according to ISS. (**G**) Comparison between chicken supertypes and mouse inhibitory subclasses based on three different methods. Scores were scaled between 0 and 1 per method and summed across all methods to represent the similarity score. White dots in tiles are shown when populations are among the top reciprocal matches according to two or all three methods. Amy, amygdala; MSN; medium spiny neuron; CP, caudate putamen; OB, olfactory bulb; CeA, central amygdala; MeA, medial amygdala; IA, intercalated amygdala.

Most inhibitory supertypes with MGE- and CGE-like identities were confidently mapped to sections of the telencephalon based on the ISS and Visium data, revealing an interspersed distribution across all regions of the pallium (Fig. 2, D and E; fig. S7). Conversely, most LGE-like supertypes display more restricted localizations. Two small cell populations (Inh_OB_CPA6, Inh_OB_FGFRL1) are predicted to be located in or migrating to the olfactory bulb (fig. S7D). The remaining LGE-like supertypes could be divided into two distinct groups based on the expression of *FOXP1* and *FOXP2*. The *FOXP2*+ group featured one very abundant, seemingly homogeneous supertype (Inh_LGE_like_FOXP2), alongside a smaller supertype of neuroblasts (Inh_LGE_like_Pre, *SOX4*+). Inh_LGE_like_FOXP2 represents the only LGE-derived supertype with a widespread distribution across the pallium and subpallium, corresponding to a population previously observed in finches (*8*) (Fig 2, D and E).

The *FOXP1*+ group segregated into two supertypes, distinguished by the expression of *SGCG* and *DGKH*, both mapping to the subpallium (Fig. 2F), parts of which were inadvertently co-dissected with the pallium. However, only the *DGKH*+ supertype also showed robust signals in parts of the caudal pallium, which comprises regions considered to correspond to different nuclei of the mammalian amygdala (Fig. 2, A, B and F; fig. S7C). Both *FOXP1*+ supertypes contain clusters expressing markers of mammalian medium spiny neurons (MSNs) of the striatum and transcriptomically similar neurons of the central amygdala (e.g., *MEIS2*, *PRKCD*, *PENK*) (*15*, *16*) (Fig. 2A). This observation is in agreement with a recent study, which indicates the existence of a central amygdala (CA)-like nucleus in chicken (*20*) and suggests that MSNs of the striatum are shared across amniotes.

In order to more comprehensively compare inhibitory neurons across amniotes, we employed three different methods (gene specificity index correlation (*7*), label transfer based on canonical correlation analysis (*21*) and SAMap (*22*)), which afford assessments of cell population similarities across species based on different gene sets (one-to-one orthologs or all orthologs) and algorithms, to overcome potential limitations of each individual approach (Methods). Moreover, to mitigate potential biases in these comparisons (i.e., differences in matching power), we downsampled cells to similar numbers across cell type populations within each species (Methods).

Our analyses revealed that nearly all chicken inhibitory supertypes exhibited clear correspondences to murine GABAergic populations, with high similarity scores and agreements between all three methods (Fig. 2G). In line with our observations based on marker expression patterns and spatial mapping, the two supertypes Inh_LGE_like_FOXP1_SGCG and Inh_LGE_like_FOXP1_DGKH harbor clusters that match either the murine MSN D1 or D2 type, but Inh_LGE_like_FOXP1_DGKH also contains two clusters that best match cells of the murine central amygdala-like nucleus (fig. S8). Notably, the widespread Inh_LGE_like_FOXP2 supertype clearly resembles cells in the murine intercalated amygdala nucleus, in accordance with their shared transcription factor markers (Fig. 2G; fig. S6). Consistently, both populations also best match a population of GABAergic cells in the lizard (fig. S9 and 10A) that is likely located in a specific region within the amygdala-like area of the lizard (*7*). We could, however, find no indication of Inh_LGE_like_FOXP2 being enriched in any specific region of the chicken pallium (Fig. 2, D and E; fig. S7F). These examples highlight that, while inhibitory neurons have retained conserved transcriptomic identities across amniote species, the spatial organization of certain groups has diversified during evolution.

## Conserved glutamatergic cell types in hippocampal and retro-hippocampal areas

Next, we sought to characterize the diversity of glutamatergic neurons in the avian pallium, a cell class that in mammals includes both evolutionarily novel and amniote-shared types (*7*). We identified 7 glutamatergic subclasses in the chicken pallium, which further split into 28 supertypes (Fig. 3, A and B). The most distinct subclass (Ex_Pre, *SOX4*+) represents glutamatergic neuroblasts and harbors at least two distinct supertypes (Ex_Pre_SATB2, Ex_Pre_KCNH7). This finding corroborates that adult neurogenesis is prevalent in birds (*23*) and indicates that different glutamatergic lineages can be generated in the adult avian brain.

**Fig. 3.**
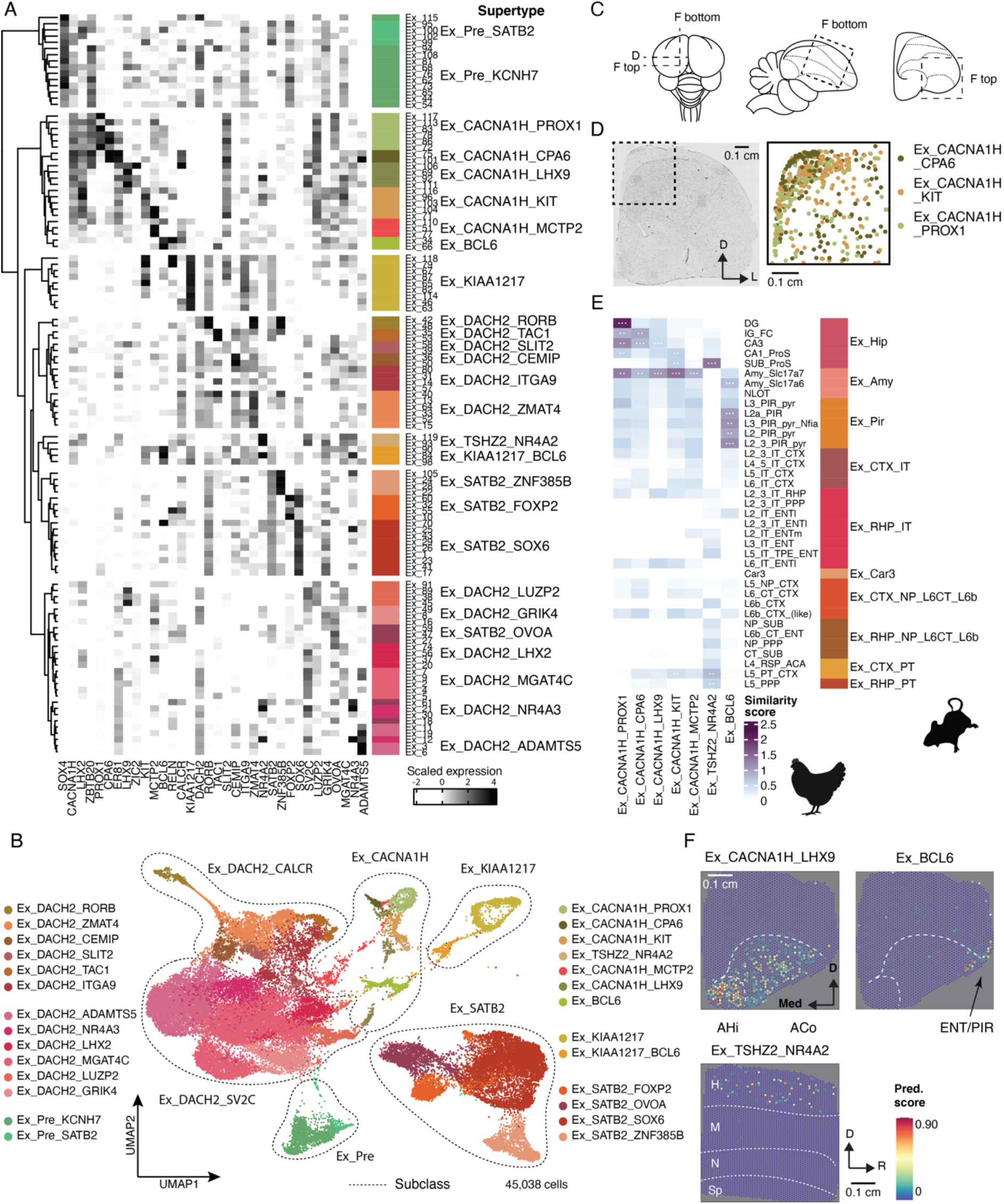
Glutamatergic excitatory neurons in the chicken (medial) pallium. (**A**) Heatmap of selected marker gene expression across 120 clusters of excitatory neurons in the chicken pallium ordered according to cluster dendrogram. Color bar and text labels on the right indicate supertype annotation. (**B**) UMAP of pallial excitatory cells colored by supertype annotation. Text labels represent subclass annotation. (**C**) Schematics of the top view (left) and a midsagittal section (right) of the chicken brain, illustrating positions of tissue sections shown in (D) and (F). Dotted lines represent borders between pallial brain regions. (**D**) Spatial location of hippocampal supertypes according to *in situ* sequencing (ISS). Only segmented cells with confidently assigned identity are shown. D, dorsal; L, lateral. (**E**) Comparison between chicken supertypes of the Ex_CACNA1H subclass and mouse excitatory subclasses based on three different methods. Color bar on the right represents broader neighbourhood labels for mouse populations taken from (*14*). Scores were scaled between 0 and 1 per method and summed across all methods to represent the similarity score. White dots in tiles are shown when populations are among the top reciprocal matches according to two or all three methods. For abbreviations in mouse annotations see Table S3. (**F**) Spatial location of other selected supertypes of the Ex_CACNA1H subclass according to Visium data. High prediction (Pred.) scores indicate high probability that cells with the respective identity were present within the spot’s area. AHi, amygdalo-hippocampal region; ACo arcopallial core nuclei; ENT, entorhinal area; PIR, piriform area; H, hyperpallium; M, mesopallium; N, nidopallium; Sp, subpallium; D, dorsal; Med, medial; R, rostral.

Among mature glutamatergic subclasses, the most distinct subclass (Ex_CACNA1H) specifically expresses *LHX2* (Fig. 3A), a transcription factor that in mammals is crucial for the formation of the isocortex and hippocampus (*24*). Consistently, two supertypes (Ex_CACNA1H_PROX1, Ex_CACNA1H_CPA6) additionally express the mammalian pan-hippocampal marker *ZBTB20*. While different hippocampal subfields (dentate gyrus, DG; Cornu ammonis regions: CA3 and CA1) have been identified in non-avian reptiles (*7*), it has previously remained unclear whether these subfields are present in birds. We find that the supertype Ex_CACNA1H_PROX1 expresses the known DG marker *PROX*1 and localizes to the most medial region of the putative chicken hippocampus equivalent, thus matching the DG’s topological position in mammals (Fig. 3D). Moreover, in our transcriptomic comparisons to glutamatergic cell type populations of the murine and lizard pallium using three different methods (see above; Methods), Ex_CACNA1H_PROX1 matches the murine DG and the lizard medial cortex neuronal types (Fig. 3E; fig. fig. S11). The second supertype, Ex_CACNA1H_CPA6 maps dorsally adjacent to Ex_CACNA1H_PROX1 and corresponds best to murine CA3 and the lizard dorso-medial cortex clusters (Fig. 3, D and E; fig. S11). Together, our observations strongly suggest that the DG and at least CA3 subfields and their cell types are also present in birds and have hence been conserved across amniotes during their evolution.

Whether further CA subdivisions (CA1, CA2) are present in the chicken medial pallium is less clear. We identified one other supertype, Ex_CACNA1H_KIT, partially expressing *ZBTB20*, which also exhibits transcriptomic resemblance to the murine CA1 (Fig. 3, A and E). However, this supertype is transcriptomically most similar to cells found in the murine amygdala, and cells from this supertype localize not only to the medial pallium (Fig. 3D), but also within the arcopallium (fig. S12B). Given these discrepant observations and the relatively low cell numbers for the Ex_CACNA1H subclass, more extensive sampling of the hippocampal region might unveil additional heterogeneity and, potentially, other hippocampal subfields in the chicken.

Despite the notable similarity of the Ex_CACAN1H subclass to the murine *Slc17a7*+ amygdala, only one supertype in this subclass, Ex_CACNA1H_LHX9, is exclusively located in the arcopallium (Fig. 3E and F). The arcopallium comprises pallial, mostly glutamatergic areas in the avian caudal pallium, which are suggested to be homologous to certain nuclei of the mammalian amygdala (*25*). Consistently, Ex_CACNA1H_LHX9 matches both murine *Slc17a7*+ and lizard amygdala clusters (Fig. 3E; fig. S11), which suggests that *SLC17A7*+ cell populations of the amygdala are conserved across amniotes. We did not find extensive similarity between the murine *Slc17a6*+ amygdala or nucleus of the lateral olfactory tract (NLOT) and any chicken supertype. However, one cluster belonging to Ex_CACNA1H_LHX9 supertype expresses *ZIC2* (Fig. 3A), a known marker of cells in the NLOT in mammals (*26*), arguing that deeper sampling of this subclass might reveal further heterogeneity. Interestingly, Ex_CACNA1H_LHX9 also exhibits transcriptomic similarity to murine cortical layer 5 pyramidal tract (L5 PT) neurons (Fig. 3E), suggesting that this supertype represents the primary output populations of the DVR circuit known to reside in the arcopallium (*27*).

Closely related to the Ex_CACNA1H subclass, is a small supertype, Ex_BCL6, selectively expressing *BCL6* and *RELN*. It is located in the putative entorhinal and/or piriform area of the chicken pallium, and corresponds to the murine piriform cortex and lizard lateral cortex neurons (Fig. 3, A and F; fig. S11). Despite an incongruity in the comparison across all three species - in our comparisons the lizard lateral cortex matches best a murine population in the entorhinal cortex, not in piriform cortex (fig. S10B) - this supertype likely corresponds to a conserved olfactory-related population located in either or both piriform or entorhinal regions (*28*).

The small Ex_TSHZ2_NR4A2 supertype does not cluster with the Ex_CACNA1H subclass in the glutamatergic dendrogram, but in addition to the very specific expression of *NR4A2* and *TSHZ2*, it exhibits moderate expression of *LHX2* and *ER81*, markers of the Ex_CACNA1H subclass (Fig. 3A). Spatial analysis reveals its sparse distribution across the apical hyperpallium (HA), the region of the hyperpallium directly bordering the hippocampal areas (Fig. 3F; fig. S12C). Intriguingly, this supertype demonstrates high similarity to the murine subiculum, which represents the major output structure of the hippocampus (*29*), as well as low similarity to exratelencephalic-projecting neurons of the isocortex and retrohippocampal area (L5_PT_CTX, L5_PPP) (Fig. 3E). This supertype might therefore represent the outward projecting neurons of the hyperpallium, known to reside in the HA (*30*), and raises the possibility that at least this subregion of the hyperpallium is more closely related to the mammalian retrohippocampal region than to the isocortex, despite previous models suggesting homology of the complete hyperpallium to the isocortex (*6*). This notion is in agreement with observations in our comparison to the lizard, in which Ex_TSHZ2_NR4A2 is most similar to clusters of the (posterior) dorsal cortex, which in turn match best the mammalian subiculum (fig. S10B and S11).

In sum, many of the glutamatergic supertypes in the hippocampal formation, including the subiculum and retrohippocampal areas, represent amniote-shared cell types.

## Excitatory neurons in the mesopallium (lateral pallium) resemble mammalian deep layer structures

Structures arising from the lateral and ventral areas of the pallium have greatly expanded proportionately in birds compared to mammals (*31*). The lateral pallium has been suggested to give rise to the claustrum, dorsal endopiriform nucleus and insular cortex in mammals, whereas in birds it develops into the anterior dorsal DVR, the so-called mesopallium (*32*). *SATB2* has been reported to represent a mesopallial marker (*33*), and, consistently, we identify two subclasses, Ex_SATB2 and Ex_KIAA1217, which strongly express *SATB2* and are both mostly located in the mesopallium (Fig. 3A and Fig. 4B; fig. S12D-F).

**Fig. 4.**
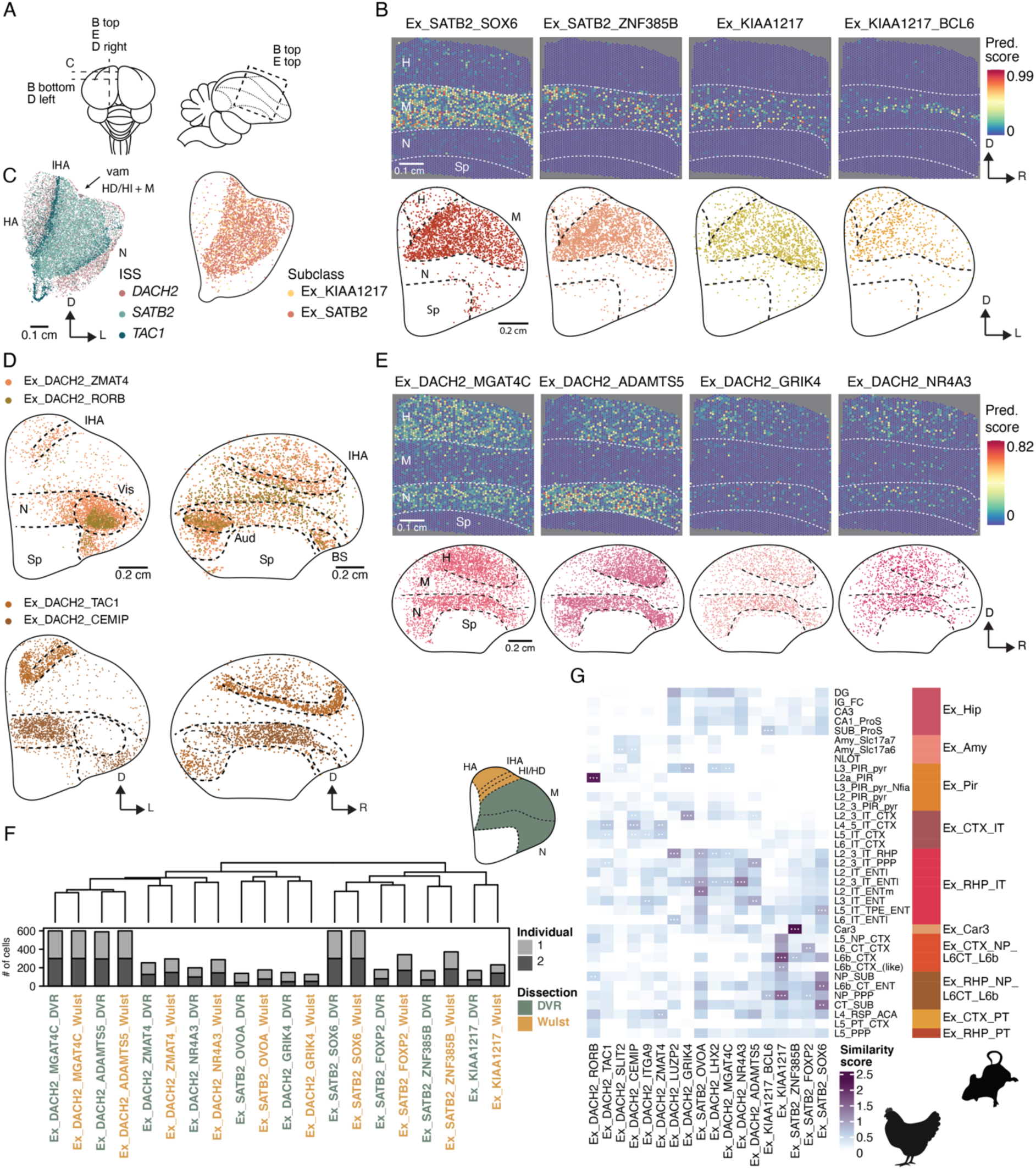
Excitatory neurons in the chicken mesopallium and nidopallium. (**A**) Schematics of chicken brain from the top (left) and in a sagittal section (right) to illustrate positions of tissue sections shown in (B - E). Dotted lines: borders between pallial brain regions. (**B**) Spatial location of mesopallial supertypes. In Visium sections (top) high prediction (Pred.) scores indicate high probability that cells with the respective identity were present within the spot’s area. In ISS sections (bottom) only segmented cells with confidently assigned identity are shown. H, hyperpallium; M, mesopallium; N, nidopallium; Sp, subpallium; D, dorsal; R, rostral; L, lateral. (**C**) Border between mesopallium and hyperpallium. Left: expression of three representative marker genes as profiled by ISS, marking the apical hyperpallium (HA) and nidopallium (N), the intercalated hyperpallium (IHA) and the densocellular hyperpallium, intermediate hyperpallium (HI), and mesopallium (HD+M). Right: area populated by mesopallial cell populations as predicted based on joint ISS profile. Vam, medial vallecula. (**D**) Spatial location of supertypes of the Ex_DACH2_CALCR subclass mostly in and around thalamic input receiving areas in the hyperpallium/nidopallium based on ISS. Vis, visual nidopallial nucleus (also often called entopallium); Aud, auditory L2 field; BS, basal somatosensory nucleus. (**E**) Localization of the most abundant supertypes of the EX_DACH2_SV2C subclass in the hyperpallium/nidopallium according to Visium (top) and ISS (bottom). (**F**) Correlation dendrogram of pseudobulk expression profiles per supertype and dissection (at least 20 cells each). Bootstrap support (n = 1000) was above 80% for all nodes. Schematic top right: illustration of borders between dissections. (**G**) Comparison between hyperpallial, mesopallial, and nidopallial chicken supertypes and mouse excitatory subclasses based on three different methods. Color bar on the right represents broader neighbourhood labels for mouse populations taken from (*14*). Scores were scaled between 0 and 1 per method and summed across all methods to represent the similarity score. White dots in tiles are shown when populations are among the top reciprocal matches according to two or all three methods. For abbreviations in mouse annotations see Table S3.

Ex_SATB2 subclass was split into four supertypes (Fig. 3, A and B) of which only one (Ex_SATB2_OVOA) showed a clear signal outside of the mesopallium (fig. S12D). Despite sharing clear similarities with other supertypes within the Ex_SATB2 subclass (*SATB2*+, *DACH2*-), Ex_SATB2_OVOA clustered with *SATB2*-negative supertypes from other subclasses in the correlation dendrogram (Fig. 3A). Other Ex_SATB2 supertypes are rather homogeneously distributed across the mesopallium (Fig. 4, B and C; fig. S12E). The other *SATB2*+ subclass, Ex_KIAA1217, includes one more abundant supertype, Ex_KIAA1217, and one very small supertype Ex_KIAA1217_BCL6, of which the former also displays a rather homogeneous distribution across the mesopallium and the latter is enriched in an area in the center of the mesopallium (Fig. 4B)

The exact border between the mesopallium and hyperpallium, which represents the putative dorsal pallial derived structure in birds, has been debated (*32*, *34*). We find that cells with Ex_SATB2 and Ex_KIAA1217 subclass identity clearly extend dorsally beyond a groove previously used as an anatomical landmark to discern hyperpallium and mesopallium (*35*), and border a region strongly expressing *TAC1* (Fig. 4C; fig. S12F), which represents the intercalated hyperpallium (IHA; discussed further below). These results indicate that the regions currently annotated as densocellular hyperpallium (HD) and intermediate hyperpallium (HI) contain mesopallial cell types, which raises the question whether this region truly develops from a distinct dorsal pallial sector as has previously been implied (*6*). Additionally, we did not find any obvious evidence for the division of the mesopallium into a dorsal and a ventral region in terms of cell type composition, because all Ex_SATB2 and Ex_KIAA1217 supertypes mapped to both regions. These findings agree with previous observations made at the bulk RNA-seq level in zebra finch (*34*).

In our cross-species analyses, supertype Ex_SATB2_ZNF385B exhibited especially high similarity to cells in the lizard anterior medial DVR (amDVR) and mouse Car3 population (Fig. 4G; fig. S11). The latter is found in deep layers of the murine cortex and especially the claustrum (*36*) and the lizard amDVR represents the claustrum homologue in reptiles (*13*). Our work thus unveils clear correspondences of the claustrum across the three representative amniote species and a high conservation of its major constituent cell type during amniote evolution.

Ex_SATB2_SOX6 and Ex_SATB2_FOXP2 supertypes exhibit slightly weaker transcriptome similarities to any lizard and mouse cell type populations (Fig. 4G; fig. S11). While their closest match in the lizard is also the amDVR, these supertypes mainly match corticothalamic (CT) projecting neurons from various areas in mouse (Fig. 4G), thus resulting in ambiguous correspondences between the three species. Should the similarities to murine CT populations reflect homology, this would be unexpected, given that neurons in the mesopallium predominantly form intra-telencephalic connections (*32*), whereas CT neurons project to the thalamus. Moreover, according to current models, these populations are born from different developmental pallial sectors, as avian mesopallial cell types arise from the lateral pallium, whereas murine CT neurons are mainly derived from the dorsal and medial pallia (*6*). Alternatively, if the similarities to cells in the lizard amDVR reflect homology, our observations suggest a possible origin of the Ex_SATB2_SOX6 and Ex_SATB2_FOXP2 supertypes by diversification of an overall claustrum-like identity in the avian lineage. However, this seems unlikely, given that it would in turn suggest that similarities to murine CT neurons arose convergently during evolution despite the lack of a shared function.

Among the Ex_KIAA1217 sublcass cells, the Ex_KIAA1217 supertype is very similar to murine Layer 6b (L6b) cells and near-projecting neurons of the subiculum (NP PPP) (Fig. 4G), whereas Ex_KIAA1217_BCL6 did not show high similarity to any mammalian cell population. Given that the similarity between Ex_KIAA1217 and murine L6b cells was one of the highest overall in the comparison of chicken to mammalian glutamatergic neurons, but Ex_KIAA1217 only exhibited very low similarity to any lizard cluster (fig. S11), we additionally compared our data to an available dataset for the turtle pallium (*7*). Specifically neurons from the anterior dorsal cortex were sampled more deeply in this dataset, as this region is larger in turtles compared to lizards (*7*). In this comparison, we identified a robust three-way correspondence between Ex_KIAA1217, murine L6b, and a turtle cluster situated in layer 2a of the turtle anterior dorsal cortex (e08) (fig. S13 and S14). L6b represents the deepest layer in the mammalian cortex and is derived from the subplate, a mostly transient structure located beneath the developing cortical plate in mammals (*38*). Although a subplate structure likely does not exist in reptiles (*39*), our observations suggest that subplate-like cells were present in the last common ancestor of all amniotes.

## Shared cell types in the avian hyperpallium and nidopallium

The evolutionary relationships of the avian nidopallium and hyperpallium have been especially intensely debated (*5*, *6*, *34*). The nidopallium is derived from the ventral pallium and represents the majority of the avian DVR (*35*) (Fig. 1A). Parts of the nidopallium receive sensory input relayed by the thalamus, and together with the mesopallium and arcopallium, they constitute the DVR sensory circuit (*31*). The hyperpallium is thought to arise from the dorsal pallium (*6*), and has been divided into several anatomical substructures, which are arranged in a columnar fashion and – as seen from the dorso-medial to ventro-lateral level – are called apical hyperpallium (HA), intercalated hyperpallium (IHA), intermediate hyperpallium (HI) and densocellular hyperpallium (HD) (*35*). These substructures constitute a second pallial sensory circuit, with IHA being the main input structure, HI and HD mostly forming tangential connections within the hyperpallium, and HA projecting to targets outside of the hyperpallium (*3*).

The majority of cells in the nido- and hyperpallium belong to two glutamatergic subclasses, Ex_DACH2_CALCR and Ex_DACH2_SV2C (Fig. 3A). Ex_DACH2_CALCR was split into six supertypes (Fig. 3, A and B; Table S2), of which several seem to be directly associated with receiving and processing sensory input according to their spatial location in specific nidopallial or hyperpallial regions. For instance, Ex_DACH2_RORB supertype maps to the primary sensory input areas of the nidopallium (Fig. 4D), whereas Ex_DACH2_TAC1 supertype is located in the primary input region of the hyperpallium (IHA) (Fig. 4D; fig. S12F and H). Surprisingly, one supertype, Ex_DACH2_ZMAT4, exhibited strong signals in both regions, in the nidopallium (around primary sensory input areas) and in the hyperpallium (within and surrounding the IHA) (Fig. 4D). We observed an even more pronounced intermixing of regional identities in the largest subclass, Ex_DACH2_SV2C, comprising six supertypes. The most abundant supertypes of this subclass showed strong, widespread signals in both regions (Fig. 4E), the nido- and the hyperpallium. And notably, clusters belonging to supertypes Ex_DACH2_NR4A3, Ex_DACH2_GRIK4, and Ex_DACH2_ADAMTS5 were of entirely mixed origin based on the dissection information (fig. S2D). These strikingly mixed regional identities of supertypes within one subclass or even within supertypes and clusters, indicate a high degree of similarity between the nidopallium and hyperpallium. Tangential migration between the nidopallium and hyperpallium was shown to be absent (*19*). Consequently, the observed regional similarity suggests that potentially shared functions within different sensory circuits result in notable similarities between these regions with distinct developmental origins. Supertype Ex_DACH2_NR4A3, belonging to the Ex_DACH2_SV2C subclass, specifically expresses activity-related genes (Table S2). This further raises the possibility that the similarity between cell populations from the two regions may not only arise from shared function, but also from shared cell states.

To investigate this striking similarity further at the transcriptional and regulatory level, we used dissections from a second individual, profiling the Wulst (encompassing the majority of the hyperpallium) and the DVR (encompassing nido- and mesopallium) separately using single-nucleus multiome ATAC and RNA sequencing (Methods). We recovered high-quality nuclei for 8,119 cells, of which 4,801 were glutamatergic and were confidently assigned to the previously defined supertypes (fig. S17). In agreement with our previous observations above, the correlation of transcriptome pseudobulks per supertype and region indicates that cell type populations in the nido- and hyperpallium are more similar between the two distinct regions than to any other cell type population within the same region (Fig. 4F), which suggests that adult cell type signals override potential developmental differences between the dorsal or ventral pallium.

We next sought to identify differentially expressed genes between these two regions within each shared supertype (Methods). As pointed out above, within the putatively sensory-related subclass Ex_DACH2_CALCR, only supertype Ex_DACH2_ZMAT4 showed strong signals of being present in both the nidopallium and hyperpallium (Fig. 4D). We found 24 significantly differentially expressed genes between Wulst (encompassing the majority of the hyperpallium) and DVR dissections for this supertype (Table S4), including the known DVR-specific transcription factor gene *NR2F2* and the GABA receptor gene *GABRG3* as the top Wulst-specific gene, despite comparably low cell numbers per dissection for this supertype in the second replicate (Fig. 4F; Table S2). This indicates that with deeper coverage, one might be able to differentiate the hyper-and nidopallium more clearly within the Ex_DACH2_ZMAT4 supertype. For the supertypes belonging to Ex_DACH2_SV2C that have contributions from both regions (Fig. 4, E and F), only the most abundant supertype Ex_DACH2_MGAT4C showed significant differential expression between Wulst and DVR dissections of 15 DVR specific genes (Table S4). Thus, despite much higher cell numbers compared to Ex_DACH2_ZMAT4 being available for supertypes in the Ex_DACH2_SV2C subclass (Fig. 4F; Table S2), fewer or, in most supertypes, no significantly differentially expressed genes could be identified within this subclass (Table S4).

Subsequently, we sought to investigate whether the distinct developmental origins of supertypes, which are shared between the hyper- and nidopallium in the adult, would still be detectable in their chromatin accessibility profiles. However, we could not identify robust differences between DVR- and Wulst-derived cells of the same supertype identity. Correlation of pseudobulks per supertype and region showed again that supertypes shared between dissections were most often similar to each other, rather than to other supertypes from the same dissection (fig. S17G). Moreover, although we detected 12,595 differentially accessible regions across all cell types, and 3,932 across excitatory neuron supertypes, at FDR <5%, we were unable to detect any significant differentially accessible regions when comparing matching supertypes between Wulst and DVR. Taken together, our results indicate that the gene regulatory programs and transcriptional profiles determining the identity of the supertypes shared between the hyper- and nidopallium, especially the ones belonging to the Ex_DACH2_SV2C subclass, fully override any profiles reflecting their different developmental origins.

In our comparisons to the lizard, despite the mixed regional origin of several supertypes, the Ex_DACH2_CALCR and Ex_DACH2_SV2C subclasses overall showed highest similarity to the DVR and not the dorsal (nor medial) cortex (fig. S11). Most supertypes belonging to the Ex_DVR_CALCR subclass are most similar to the sensory-recipient anterior lizard DVR (aDVR). The most mixed supertype within this subclass (Ex_DACH2_ZMAT4) also shows predominant similarity to the aDVR. However, in the comparison to turtle, in which the anterior dorsal cortex (aDC) is larger than in lizards (see above), we find that the predominantly hyperpallial supertypes (Ex_DACH2_TAC1,Ex_DACH2_ITGA9) are most similar to clusters of the aDC (fig. S13). Consistently, the mixed supertype Ex_DACH2_ZMAT4 shows similarity to both regions in the turtle, the aDVR and aDC.

When we compare Ex_DACH2_CALCR to mouse, we mostly observe low similarities to L4 and L5 of the murine isocortex (Fig. 4G). Cell types in primary sensory input areas of the DVR had been suggested to be homologous to L4 of the mammalian isocortex (*33*, *40*). However, specifically the supertype located within these sensory regions (Ex_DACH2_RORB) is the only supertype within the subclass that does not resemble isocortical cell types. Instead, it is highly similar to cells in L2a of the murine piriform cortex (Fig. 4G). Cells in L2a of the piriform cortex arise from the ventral pallium and are one of the main recipient neurons of olfactory bulb input in mice (*41*). This suggests that sensory processing neuron populations in the DVR employ similar sets of genes as neurons in the mammalian isocortex, but that their ancestry may lie in more ancient olfactory input processing neurons.

Supertypes belonging to the Ex_DACH2_SV2C subclass correspond best to clusters from the (posterior) DVR in lizard, despite their mixed regional origin in the chicken (Fig. 4E; fig. S11). Also, in contrast to the Ex_DACH2_CALCR subclass, even the most mixed supertypes are rather similar to the turtle DVR and not the dorsal cortex, although similarities are overall low (Fig. 4E; fig. S13). In comparison to mouse, Ex_DACH2_SV2C exhibits mostly very low similarity to retrohippocampal areas and the entorhinal cortex (Fig. 4G). Ex_DACH2_GRIK4 is the only chicken supertype, which displays some similarity to mammalian neurons in isocortical layers 2/3. However, all supertypes of the Ex_DACH2_SV2C subclass are most similar to lizard clusters, which, in turn, match the murine piriform cortex or amygdala, suggesting that the low similarity scores in these comparisons do not reflect homology. Thus, the evolutionary origins of Ex_DACH2_CALCR and especially Ex_DACH2_SV2C as well as the mixed regional cellular identities between the hyper- and nidopallium remain elusive.

## Developmental and evolutionary origins of avian excitatory cell types

Notably, the striking transcriptional similarity we observed between the nidopallium and hyperpallium for major cell types had been previously suggested based on bulk-tissue data and was interpreted to reflect a developmental origin, wherein cell populations surrounding the ventricle give rise to analogous populations above and below it (*34*). However, this hypothesis conflicts with other developmental studies that show a confinement of neuronal lineages to each portion of the pallium (*19*, *42*).

We therefore generated snRNA-seq data for the chicken pallium across eight developmental stages, from early-mid neurogenesis (embryonic day 6, E6) to late *in ovo* development (E19) (chicks hatch at around days 20/21 (*43*)) (Methods). From stage E8 onwards, the pallium of at least one individual was dissected into its dorsal and ventral halves, splitting the prospective mesopallium in the middle (Fig. 5A), at the proposed axis of dorso-ventral symmetry (*34*); dissections were then profiled separately. We obtained high-quality snRNA-seq data for a total of 142,429 cells, which we assigned to 14 major cell classes (Fig. 5B and C; fig. S16).

**Fig. 5.**
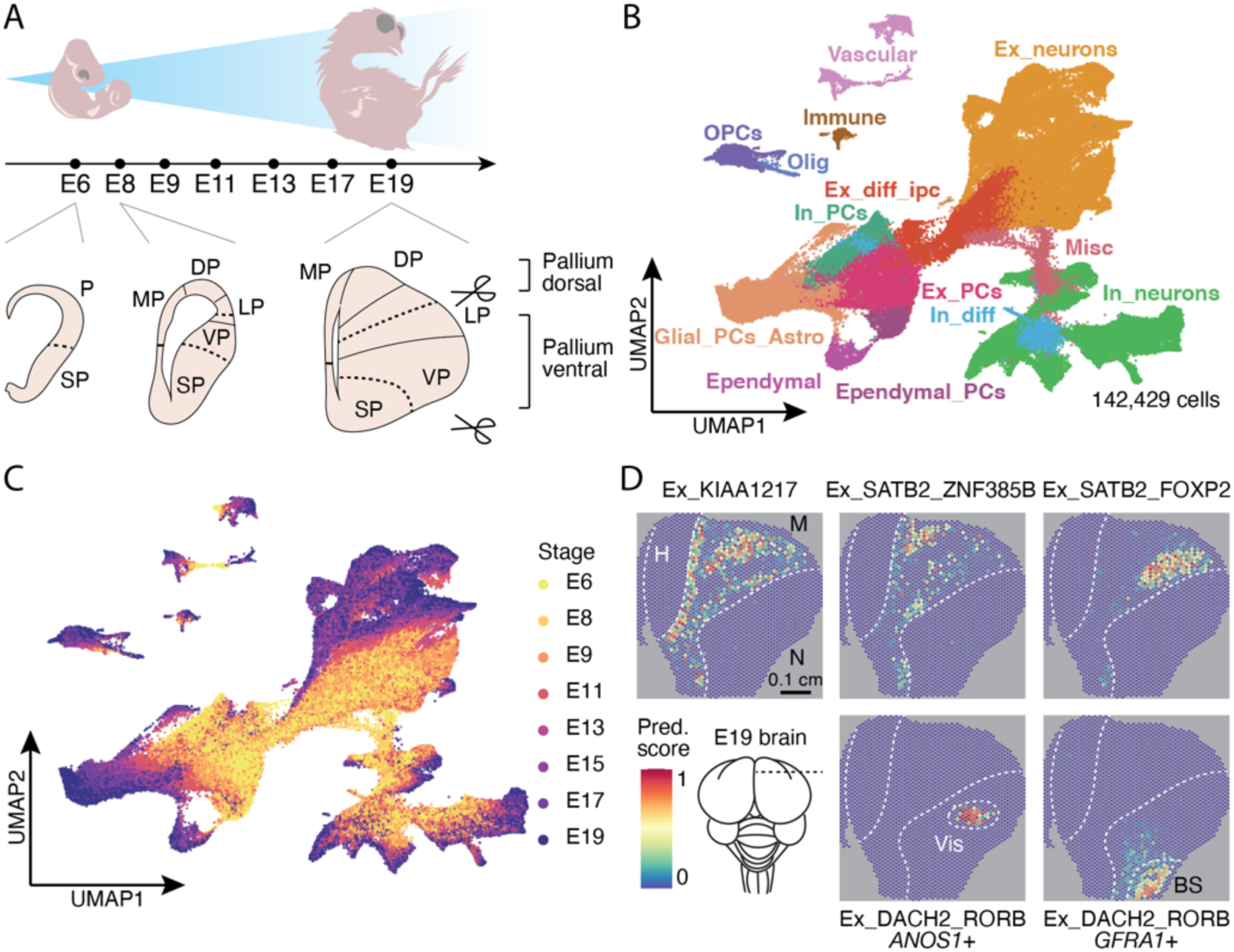
Cellular development of the chicken pallium. (**A**) Illustration of sampling strategy across pallial development in the chick. Samples were taken on seven different days of embryonic (E) development. At E6 the pallium (P) was collected as a whole, from E8 onwards the pallium was dissected into two halves for at least one individual per stage. SP, subpallium; MP, medial pallium; DP, dorsal pallium; LP, lateral pallium; VP, ventral pallium. (**B**) UMAP of pallial cells colored by and labelled with class annotation and (**C**) colored by developmental stage. Ex_neurons, excitatory neurons; OPCs, oligodendrocyte progenitor cells; Olig, oligodendrocytes; Ex_diff_ipc, excitatory intermediate progenitors and differentiating cells; In_PCs, inhibitory progenitors; In_diff, inhibitory differentiating neurons; In_neurons, inhibitory neurons; Glial_PCs_Astro, late radial glia (gliogenic) and astrocytes; Ex_PCs, radial glia (neurogenic); Ependymal_PCs, ependymal progenitor cells; Misc, miscellaneous. (**D**) Spatial location of selected excitatory supertypes in E19 Visium sections. E19 snRNA-seq data was integrated with adult data to predict supertypes labels. High prediction (Pred.) scores indicate high probability that cells with the respective identity were present within the spot’s area. Schematic of brain from top view (bottom left) illustrates position of the section.

We also profiled three sections of the E19 pallium using the Visium approach. Populations of mature neurons from this last profiled developmental stage were highly similar to supertypes defined in the adult (fig. S17), which indicates that most supertypes are already present at this stage. We therefore used these data to corroborate and refine our pallium cell type atlas. Overall, E19 mapping confirmed our observations in the adult but offered a clearer mapping in some cases (Fig. 5D; fig. S18). For instance, Ex_BCL6, which in the adult seemed to be located in the piriform and/or entorhinal area, clearly maps to an area separated from the nidopallium by the ventricle, arguing for its medial pallial origin and, therefore, rather for an entorhinal identity of this supertype (fig. S18F). Especially mesopallial supertypes exhibited a more refined spatial location (Fig. 5D; S19 B and C) compared to the adult pallium, indicating that the mesopallium is less homogeneous in its cell type composition than inferred from our adult spatial analyses (Fig. 4A). The E19 data also uncovered additional heterogeneity within the thalamic input receiving populations of the DVR (Fig. 5D).

To trace the developmental origins of the transcriptional similarity between glutamatergic cell populations in the adult nido- and hyperpallium (see above), we focused on the glutamatergic lineage, including radial glia, intermediate progenitors/differentiating cells, and excitatory neurons (Fig. 6A). We converted the developmental trajectory, clearly identifiable from the integration of sampled stages, into a continuous pseudotime progression (*44*) (Methods) (Fig. 6B). To assign early neurons to sublineages, we adapted a previous approach (*45*), which uncovered four major lineages of early neurons, corresponding to the medial/arco-, meso-, hyper- and nidopallial lineages, respectively (Fig. 6F; fig. S19).

**Fig. 6.**
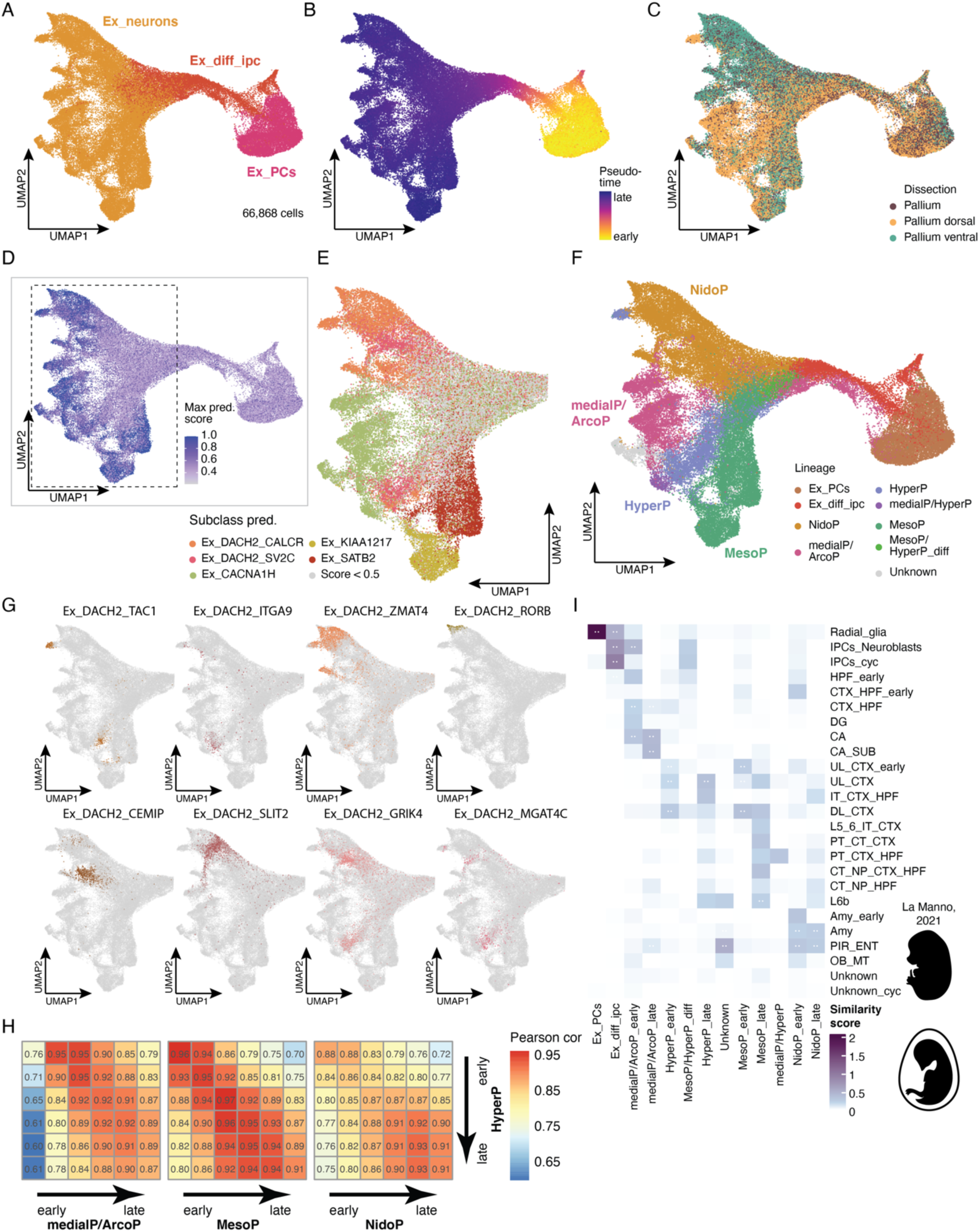
Developing excitatory neurons in the chicken pallium. UMAPs of excitatory neuron lineage colored by and labelled with (**A**) class annotation (Ex_PCs, radial glia (neurogenic); Ex_diff_ipc, excitatory intermediate progenitors and differentiating cells; Ex_neurons, excitatory neurons), (**B**) colored by diffusion pseudotime, (**C**) colored by dissection origin and (**D**) colored by maximal prediction (Max pred.) score for any adult excitatory subclass after CCA label transfer (**E**) Zoom into UMAP shown in (A – D) to show only early neurons colored by adult subclass prediction if prediction score (D) was above 0.5. (**F**) UMAP of excitatory neuron lineage colored by and labelled with sublineage annotation. NidoP, nidopallial lineage; medialP/ArcoP, medial and arcopallial lineage; HyperP, hyperpallial lineage; MesoP, mesopallial lineage; medialP/HyperP, between medial and hyperpallial lineage; MesoP/HyperP_diff, early differentiating cells of mesopallial or hyperpallial lineage. (**G**) UMAPs of early neurons as shown in (E), colored by most abundant supertypes in hyper- and nidopallial lineage as predicted by label transfer from adult data. (H) Heatmap of Pearson correlation between the hyperpallial and other early neuron sublineages. Sublineages were split into six bins according to pseudotime, each represented by one tile in the correlation heatmap. Values in tiles represent correlation values. (I) Comparison between chicken pallial lineages as shown in (F) and populations in the developing murine pallium (subset of (*47*), embryonic day 9 - 17). Lineages comprising many cells in chick were split into early and late (pseudotime (B) > 0.89). Comparison is based on two methods. Scores were scaled between 0 and 1 per method and summed across both methods to represent the similarity score. White dots in tiles are shown when populations are among the top reciprocal matches according to both methods. IPCs, intermediate progenitor cells; cyc, cycling; HPF, hippocampal formation; DG, dentate gyrus; CA, Cornu Ammonis; SUB, subiculum; UL, upper cortical layers; CTX, cortex; IT, intra-telencephalic; DL, deep cortical layers; PT, pyramidal-tract-projecting; NP, near-projecting; CT, corticothalamic-projecting; Amy, amygdala; PIR, piriform cortex; ENT, entorhinal cortex; OB_MT, olfactory bulb mitral tufted cells.

When transferring adult subclass labels, cells in the hyperpallial and nidopallial lineage were predominantly predicted to belong to the two subclasses, which comprise cell types from both regions in the adult (Ex_DACH2_CALCR, Ex_DACH2_SV2C) (Fig. 6, D and E; Fig. 4, D and E). However, contrary to the adult data, there is only minimal mixing of cells from dorsal and ventral dissections between these two lineages (Fig. 6C). Moreover, when we transferred adult supertype labels, cells in the nidopallial and hyperpallial lineage were mostly predicted to belong to supertypes that are also exclusive to the respective regions in the adult (Fig. 6G). By contrast, the most extensively mixed supertypes from the adult Ex_DACH2_SV2C subclass showed only limited presence in the developmental dataset (Fig. 6G). Together, our observations suggest that the hyperpallium and nidopallium have distinct developmental origins despite their extensive similarities in the adult; and, until E19, both structures comprise mostly transcriptomically different cell types.

Nevertheless, there are indications of the adult similarities in late developmental stages. A small population of mature hyperpallial *TAC1*+ nuclei clusters with mature neurons of the nidopallial lineage in low-resolution Louvain clustering (fig. S19C). Furthermore, a correlation analysis of the different lineages along pseudotime reveals that the correlation between the hyperpallial and nidopallial lineage increases during development (Fig. 6H). Together, these observations indicate that the similarity between the hyperpallial and nidopallial lineage increases progressively during development but is not yet extensive at E19. Notably, in two parallel studies, Rueda-Alaña et al. independently observe some cell type populations distributed across the hyper- and nidopallium at E15, and Hecker et al. (https://doi.org/10.1101/2024.04.17.589795) detect a strong similarity between the hyper- and nidopallium at post-hatch day 15, suggesting that the trend we see in late stages of *in ovo* development continues after hatching and is completed by day 15 of *ex ovo* development at the latest.

Given the clear differences between glutamatergic lineages originating from distinct territories in our developmental data, and considering the generally higher conservation of gene expression across species during development compared to adults (*46*), we reasoned that our data may facilitate the identification of broader regional homologies between avian and mammalian pallial structures. We thus compared our chicken developmental data to a subset of an available dataset of the developing mouse brain (*47*) in which we annotated the pallial glutamatergic lineage in more detail (Methods) (fig. S20).

We found that the two different types of progenitor cells (radial glia, RG; intermediate progenitor cells, IPCs) are the cell types with the highest similarity between the two species (Fig. 6I). It has previously been debated if IPCs exist in chicken (*48*), but we find a clear correspondence of potential IPCs in the chicken to mouse IPCs (Fig. 6I). Moreover, we also identified a sizable cluster of IPCs in the chicken that are likely to be cycling (fig. S19B), which indicates that IPCs do proliferate in birds. However, judging by the numbers of cycling IPCs relative to cycling RGs, IPCs in chicken are likely not proliferating to the same extent as in mammals, in accord with previous work (*48*, *49*) and findings in a related study by Rueda-Alaña et al.

The comparison of early neuronal lineages between both species largely corroborates our findings in the adult, while also unveiling relationships that were not discernible in the adult stage. The conservation of the hippocampus is evident during development as well, but in contrast to the adult we observe very little similarity to the mammalian amygdala (Fig. 6I). Interestingly, the hyperpallial lineage, that is the lineage likely forming the IHA and potentially HA, is most similar to upper and deep layer cortical neurons during development (Fig. 6I), in line with adults, where we observed low similarity to neurons in L4/5 (Fig. 4G). The mesopallial lineage matches best early and more mature deep layer cortical neurons. We observed a similar trend in the adult for most supertypes located in the mesopallium, except for one (Ex_SATB2_ZNF385D) that is homologous to the claustrum in the adult (Fig. 4G). Given the structure’s small size, we could not identify any claustral population in the developing mouse brain dataset to confirm this match in development. Interestingly, the nidopallial lineage exhibited the strongest similarity to the developing amygdala, piriform and entorhinal cortex, which was not evident in adults.

Taken together these results indicate that not only pallial radial glia, but also intermediate progenitor cells were already present in the last common ancestor of amniotes. Furthermore, early neurons in the avian pallium still exhibit clear transcriptomic signatures related to their origins in different developmental territories, which were only partially detectable in adults. These early regional identities allow the identification of broad homologies between mammals and birds.

## Discussion

In this study, we generated the first spatially resolved cell type atlas of the chicken pallium in adults and across development. Through comprehensive comparisons of this atlas to corresponding data from mouse and non-avian reptiles, we trace the origins and evolution of cell types in the amniote pallium. We confirm previous observations that inhibitory neurons are overall transcriptomically conserved across amniotes (*7*, *8*), although one LGE-derived population that resides in the small intercalated amygdala in mammals has become very abundant across the avian pallium. By contrast, excitatory neuron repertoires have diverged substantially during evolution, with some showing extraordinary convergences of gene expression programs during development in birds.

Our analyses revealed that homologs of excitatory neurons in the mammalian hippocampus are also present in corresponding regions of the chicken pallium. This finding, together with previous data for non-avian reptiles (*7*) and tetrapods (*50*), suggests that key hippocampal regions and constituent excitatory cell types were already present in the common tetrapod ancestor and have been preserved across amniotes during evolution. Notably, however, glutamatergic cell types of the hippocampal region in chicken seem to be closely related to the major glutamatergic cell type population (Ex_CACNA1H_LHX9) of the arcopallium, a region that has previously been suggested to be homologous to certain nuclei of the mammalian amygdala (*25*). In agreement with this suggestion, this arcopallial cell type exhibits strong similarities to cells from the mammalian amygdala in our comparisons. However, the avian arcopallium, as well as the mammalian amygdala, have been suggested to arise from the ventral pallium, whereas the hippocampal areas arise from the distinct medial pallial sector (*6*). By contrast, our findings indicate that the ventral and medial pallium do not represent strictly distinctive germinative zones in birds, but form one contiguous region around the caudal pole of the telencephalon. This notion is consistent with shared expression of selected genes in early chick development (*51*), as well as the close proximity of these two zones in early mammalian development (*52*) and alternative models of pallial evolution in amniotes (*53*).

We also uncover that neurons within distinct subregions of the avian hyperpallium, previously hypothesized to be homologous to the mammalian isocortex (*6*, *54*), do not exhibit a singular hyperpallial identity. Instead, populations in some subregions closely resemble cells in neighboring pallial structures. Specifically, our findings indicate that the densocellular hyperpallium forms one homogeneous territory together with the neighboring mesopallium in terms of its cell type composition, in accord with previous bulk transcriptome work (*34*), and arises from the same lineage as the rest of the mesopallium during development. In the apical hyperpallium, adjacent to the medial pallium, we identify a cell type (Ex_TSHZ2_NR4A2) characterized by the expression of marker genes of both, medial and hyperpallial lineages. In support of a rather medial identity, this cell type matches cell types of medial pallial structures in mammals and other reptiles; that is, the subiculum and the posterior dorsal cortex. The only hyperpallial subregion that clearly originates from a distinct lineage during development is the intercalated hyperpallium – the main input structure of the hyperpallium (*3*, *55*, *56*).

Intriguingly, we observe a remarkable transcriptomic and gene regulatory similarity between cell populations in the avian (dorsal) intercalated/apical hyperpallium and the (ventral) nidopallium. This similarity, which is also observed by Hecker et al. (https://doi.org/10.1101/2024.04.17.589795) in chicks at day 15, is so pronounced in the adult that several cell populations are essentially indistinguishable between the two regions. Gedman et al. (*34*), who inferred a similar pattern from bulk transcriptome analyses of the adult zebra finch pallium, suggested that the hyper- and nidopallium arise from one continuous embryonic region. However, in stark contrast to the patterns observed in all three studies in chicks/adults, we identify distinct cellular lineages for these two regions during *in ovo* development that exhibit only limited similarity at mid-neurogenesis stages. This is in line with the observation that some of the extensively shared cell type populations found in the adult seem to be mostly absent at embryonic day 19 (Fig. 6G; fig. S17). Tangential migration could explain our observations, as regions would contain similar cells due to a shared developmental origin and “new” populations could form from migrated cells being exposed to their new environment. However, previous lineage tracing studies covering development until E14 (i.e., several days after neurogenesis is complete) found no evidence for tangential migration between the hyperpallium and nidopallium (*19*). Our developmental dataset, which includes data for specific dissections of the dorsal and ventral pallium until late development (E19), does also not provide any indication of tangential cell migrations. Notably, however, the hyper- and nidopallial lineages exhibit progressively increasing similarities during development, although some of the populations that are shared between both structures in adults cannot be traced during *in ovo* development. We therefore hypothesize that the remarkable gene expression similarity between the hyper/nidopallial cell populations, which carry out analogous functions in different sensory circuits, is fully established soon after hatching due to the massively increased sensory input.

The evolutionary relationships of pallial regions across amniotes have long been a subject of debate (*32–34*). Leveraging our extensive adult and developmental datasets, we validated previously suspected relationships (as for the hippocampus, see above) and uncovered unexpected homologies. Specifically, our findings indicate a notable divergence between the avian nidopallium, constituting the ventral DVR and arising from the ventral pallium during development, and its mammalian counterpart. In adult comparisons, we predominantly observe low cell type similarities in the avian nidopallium to neurons in layers 4/5 of the mammalian isocortex, as well as cells in the entorhinal cortex, suggesting an overall diverged profile. However, a distinct cell type within the main thalamic input receiving areas of the nidopallium exhibits strong similarity to cells in the mammalian piriform cortex. In agreement with this correspondence, developmental data indicate clear cellular similarities between the avian nidopallial lineage and the mammalian amygdala, piriform cortex, and/or entorhinal cortex – structures largely derived from the putative ventral pallium in mammals as well. We thus conclude that the avian nidopallium is overall homologous to the aforementioned mammalian structures, but their constituent cell types have substantially diverged during evolution. Further developmental investigations are warranted to ascertain whether the nidopallium is homologous to all or only a subset of these structures or constituent cell types.

Despite the striking similarity between cell populations in the nidopallium and hyperpallium, our cross-species analyses suggest that only cells in the intercalated hyperpallium are homologous to cell type populations in layers 4/5 of the mammalian isocortex. This notion is in agreement with previous models indicating homology between the hyperpallium and mammalian isocortex (*6*), and with the cell types’ respective roles in the sensory circuit as input neurons (*55–57*).

Interestingly, we observe a strong similarity of the adult avian mesopallium – which seems to include an area previously defined as part of the hyperpallium (see above) – with the mammalian claustrum but also, unexpectedly, to neurons in deep layers (L6b, L6 CT) of the isocortex and retrohippocampal areas. One mesopallial population (Ex_SATB2_ZNF385B) exhibits clear similarity to the main murine claustral population and cells in the anterior medial DVR in lizards, a structure, which was in turn shown to correspond to the murine claustrum in terms of gene expression, connectivity, and function (*13*). Thus, this specific cell type has been strongly conserved across all major lineages of amniotes since their emergence. This finding is also in agreement with previous models based on developmental data, given that both the claustrum and the mesopallium are thought to emerge from the lateral pallium (*32*).

However, our findings regarding other mesopallial cell types challenge this model. First, we identify a population in the avian mesopallium and turtle anterior dorsal cortex strongly resembling neurons in mammalian L6b. Neurons in L6b are a remnant of the largely transient subplate, which is present across different pallial sectors in mammals (*58*). While a subplate structure likely does not form in the developing reptilian pallium (*39*), our observations therefore suggest that subplate-like cells were present in the last common ancestor of all amniotes. Second, other mesopallial cell types are most similar to mammalian cortico-thalamic (CT) projecting neurons in isocortical L6 and retrohippocampal areas, instead of any population derived from the mammalian lateral pallium. This similarity is particularly surprising, because cortico-thalamic neurons project to the thalamus, whereas the mesopallium predominantly projects within the telencephalon (*37*) and works as an integrating structure, rather akin to mammalian layer 2/3 intra-telencephalic neurons. However, in line with our findings in the adult, the overall similarity of the mesopallial lineage to deep layers of the isocortex and hippocampal formation is further supported by our developmental comparisons, and Hecker et al. (https://doi.org/10.1101/2024.04.17.589795) confirm this correspondence based on analyses of enhancer codes.

Despite the different projection patterns of mammalian cell types matching the avian mesopallium (claustral Car3, L6b and (L6) CT), all of these populations represent some of the earliest born neurons in the cortical neuroepithelium (*58*). The close association of these cell type populations is also reflected in their transcriptome profiles, as the claustrum and L6b share the expression of many marker genes during development (*59*), although the mammalian claustral population exhibits a quite distinct transcriptomic profile in the adult (*14*). Neurons in L6b, in turn, are closely related to L6 CT neurons based on single-cell RNA-seq data (*14*). These results suggest that the avian mesopallium might overall be homologous to early-born neurons of the mammalian lateral, dorsal, and medial pallium.

Altogether, our findings reveal a classification of glutamatergic cell types into four major developmental lineages in birds that have clearly distinct ontogenetic origins, partly consistent with the existence of a tetrapartite pallium in birds (*6*) (Fig. 1; fig. S21). However, our work also unveils a striking convergence of specific gene expression programs for the ventral and dorsal lineages that eventually completely override developmental signatures (fig. S21). This process leads to the emergence of major cell types in the nido- and hyperpallium that are essentially indistinguishable between these two spatially distant brain regions. Our cross-amniote analyses uncover general correspondences of cell populations in medial and ventral regions, of which the latter is only clearly evident during development (Fig. 6I). However, while certain cell populations are shared across amniotes within putative dorsal and lateral pallial regions, not all cell populations in these regions are homologous across amniotes (fig. S22). Thus, our work highlights the limitations of current models of pallial evolution in amniotes (*53*) and motivates further deep investigations of the evolutionary dynamics of neural development across species, including mammals.

## Acknowledgments

We thank members of the Kaessmann group for discussions during the project. We thank Dr. Monika Langlotz for her assistance with fluorescence activated cell sorting. We thank David Ibberson for his advice on Visium.

## Author contributions

BZ and HK conceived the study; BZ wrote the original manuscript with input from FG-M and HK; all authors reviewed the manuscript; FG-M and HK co-supervised the study; BZ performed most analyses and generated the data with support from CS and JS; AF collected samples; FL provided advice and code; IS performed analyses; MS provided advice; EL performed analyses; EVP collected samples; PJ collected samples, RS-G and FG-M collected samples, NT provided code and computational support; and FH-S provided advice and helped with sample collection.

## Data and materials availability

The raw and processed data generated in this study are deposited in heiDATA under https://doi.org/10.11588/data/BX6REK. All other data are provided in the manuscript. The code used to analyze the data is available at https://gitlab.com/kaessmannlab/amniote-pallium.

## Supplementary Materials

Materials and Methods

Figs. S1 to S22

Tables S1 to S6

References (*60–73*)

## References and Notes

1. O. Güntürkün, T. Bugnyar, Cognition without Cortex. Trends Cogn. Sci. 20, 291–303 (2016).

2. J. H. Kaas, Evolution of nervous systems. Evol. Nerv. Syst. 1–4, 1–2007 (2016).

3. M. Stacho, C. Herold, N. Rook, H. Wagner, M. Axer, K. Amunts, O. Güntürkün, A cortex-like canonical circuit in the avian forebrain. Science 369, eabc5534 (2020).

4. S. Kumar, M. Suleski, J. M. Craig, A. E. Kasprowicz, M. Sanderford, M. Li, G. Stecher, S. B. Hedges, TimeTree 5: An Expanded Resource for Species Divergence Times. Mol. Biol. Evol. 39, msac174 (2022).

5. S. D. Briscoe, C. W. Ragsdale, Homology, neocortex, and the evolution of developmental mechanisms. Science 362, 190–193 (2018).

6. L. Puelles, J. E. Sandoval, A. Ayad, R. del Corral, A. Alonso, J. L. Ferran, M. Martínez-de-la-Torre, The Pallium in Reptiles and Birds in the Light of the Updated Tetrapartite Pallium Model (2016)vols. 1–4.

7. M. A. Tosches, T. M. Yamawaki, R. K. Naumann, A. A. Jacobi, G. Tushev, G. Laurent, Evolution of pallium, hippocampus, and cortical cell types revealed by single-cell transcriptomics in reptiles. Science 360, 881–888 (2018).

8. B. M. Colquitt, D. P. Merullo, G. Konopka, T. F. Roberts, M. S. Brainard, Cellular transcriptomics reveals evolutionary identities of songbird vocal circuits. Science 371, eabd9704 (2021).

9. R. Ke, M. Mignardi, A. Pacureanu, J. Svedlund, J. Botling, C. Wählby, M. Nilsson, In situ sequencing for RNA analysis in preserved tissue and cells. Nat. Methods 10, 857–860 (2013).

10. T. Biancalani, G. Scalia, L. Buffoni, R. Avasthi, Z. Lu, A. Sanger, N. Tokcan, C. R. Vanderburg, Å. Segerstolpe, M. Zhang, I. Avraham-Davidi, S. Vickovic, M. Nitzan, S. Ma, A. Subramanian, M. Lipinski, J. Buenrostro, N. B. Brown, D. Fanelli, X. Zhuang, E. Z. Macosko, A. Regev, Deep learning and alignment of spatially resolved single-cell transcriptomes with Tangram. Nat. Methods 18, 1352–1362 (2021).

11. M. A. Lipiec, J. Bem, K. Kozinski, C. Chakraborty, J. Urban-Ciećko, T. Zajkowski, M. Dabrowski, Ł. M. Szewczyk, A. Toval, J. L. Ferran, A. Nagalski, M. B. Wisniewska, TCF7L2 regulates postmitotic differentiation programmes and excitability patterns in the thalamus. Dev. Camb. 147 (2020).

12. D. Hain, T. Gallego-Flores, M. Klinkmann, A. Macias, E. Ciirdaeva, A. Arends, C. Thum, G. Tushev, F. Kretschmer, M. A. Tosches, G. Laurent, Molecular diversity and evolution of neuron types in the amniote brain. Science 377, eabp8202 (2022).

13. H. Norimoto, L. A. Fenk, H. H. Li, M. A. Tosches, T. Gallego-Flores, D. Hain, S. Reiter, R. Kobayashi, A. Macias, A. Arends, M. Klinkmann, G. Laurent, A claustrum in reptiles and its role in slow-wave sleep. Nature 578, 413–418 (2020).

14. Z. Yao, C. T. J. van Velthoven, T. N. Nguyen, J. Goldy, A. E. Sedeno-Cortes, F. Baftizadeh, D. Bertagnolli, T. Casper, M. Chiang, K. Crichton, S.-L. Ding, O. Fong, E. Garren, A. Glandon, N. W. Gouwens, J. Gray, L. T. Graybuck, M. J. Hawrylycz, D. Hirschstein, M. Kroll, K. Lathia, C. Lee, B. Levi, D. McMillen, S. Mok, T. Pham, Q. Ren, C. Rimorin, N. Shapovalova, J. Sulc, S. M. Sunkin, M. Tieu, A. Torkelson, H. Tung, K. Ward, N. Dee, K. A. Smith, B. Tasic, H. Zeng, A taxonomy of transcriptomic cell types across the isocortex and hippocampal formation. Cell 184, 3222–3241.e26 (2021).

15. O. Gokce, G. M. Stanley, B. Treutlein, N. F. Neff, J. G. Camp, R. C. Malenka, P. E. Rothwell, M. V. Fuccillo, T. C. Südhof, S. R. Quake, Cellular Taxonomy of the Mouse Striatum as Revealed by Single-Cell RNA-Seq. Cell Rep. 16, 1126–1137 (2016).

16. H. Hochgerner, S. Singh, M. Tibi, Z. Lin, N. Skarbianskis, I. Admati, O. Ophir, N. Reinhardt, S. Netser, S. Wagner, A. Zeisel, Neuronal types in the mouse amygdala and their transcriptional response to fear conditioning. Nat. Neurosci. 2023, 1–13 (2023).

17. E. S. Lein, M. J. Hawrylycz, N. Ao, M. Ayres, A. Bensinger, A. Bernard, A. F. Boe, M. S. Boguski, K. S. Brockway, E. J. Byrnes, L. Chen, L. Chen, T. M. Chen, M. C. Chin, J. Chong, B. E. Crook, A. Czaplinska, C. N. Dang, S. Datta, N. R. Dee, A. L. Desaki, T. Desta, E. Diep, T. A. Dolbeare, M. J. Donelan, H. W. Dong, J. G. Dougherty, B. J. Duncan, A. J. Ebbert, G. Eichele, L. K. Estin, C. Faber, B. A. Facer, R. Fields, S. R. Fischer, T. P. Fliss, C. Frensley, S. N. Gates, K. J. Glattfelder, K. R. Halverson, M. R. Hart, J. G. Hohmann, M. P. Howell, D. P. Jeung, R. A. Johnson, P. T. Karr, R. Kawal, J. M. Kidney, R. H. Knapik, C. L. Kuan, J. H. Lake, A. R. Laramee, K. D. Larsen, C. Lau, T. A. Lemon, A. J. Liang, Y. Liu, L. T. Luong, J. Michaels, J. J. Morgan, R. J. Morgan, M. T. Mortrud, N. F. Mosqueda, L. L. Ng, R. Ng, G. J. Orta, C. C. Overly, T. H. Pak, S. E. Parry, S. D. Pathak, O. C. Pearson, R. B. Puchalski, Z. L. Riley, H. R. Rockett, S. A. Rowland, J. J. Royall, M. J. Ruiz, N. R. Sarno, K. Schaffnit, N. V. Shapovalova, T. Sivisay, C. R. Slaughterbeck, S. C. Smith, K. A. Smith, B. I. Smith, A. J. Sodt, N. N. Stewart, K. R. Stumpf, S. M. Sunkin, M. Sutram, A. Tam, C. D. Teemer, C. Thaller, C. L. Thompson, L. R. Varnam, A. Visel, R. M. Whitlock, P. E. Wohnoutka, C. K. Wolkey, V. Y. Wong, M. Wood, M. B. Yaylaoglu, R. C. Young, B. L. Youngstrom, X. F. Yuan, B. Zhang, T. A. Zwingman, A. R. Jones, Genome-wide atlas of gene expression in the adult mouse brain. Nature 445, 168–176 (2007).

18. L. Lim, D. Mi, A. Llorca, O. Marín, Development and Functional Diversification of Cortical Interneurons. Neuron 100, 294–313 (2018).

19. F. García-Moreno, E. Anderton, M. Jankowska, J. Begbie, J. M. Encinas, M. Irimia, Z. Molnár, Absence of Tangentially Migrating Glutamatergic Neurons in the Developing Avian Brain. Cell Rep. 22, 96–109 (2018).

20. B. Yu, Q. Zhang, L. Lin, X. Zhou, W. Ma, S. Wen, C. Li, W. Wang, Q. Wu, X. Wang, X.-M. Li, Molecular and cellular evolution of the amygdala across species analyzed by single-nucleus transcriptome profiling. Cell Discov. 2023 91 9, 1–22 (2023).

21. T. Stuart, A. Butler, P. Hoffman, C. Hafemeister, E. Papalexi, W. M. Mauck, Y. Hao, M. Stoeckius, P. Smibert, R. Satija, Comprehensive Integration of Single-Cell Data. Cell 177, 1888–1902.e21 (2019).

22. A. J. Tarashansky, J. M. Musser, M. Khariton, P. Li, D. Arendt, S. R. Quake, B. Wang, Mapping single-cell atlases throughout metazoa unravels cell type evolution. eLife 10 (2021).

23. A. Barnea, V. Pravosudov, Birds as a model to study adult neurogenesis: bridging evolutionary, comparative and neuroethological approaches. Eur. J. Neurosci. 34, 884–907 (2011).

24. S.-J. Chou, S. Tole, Lhx2, an evolutionarily conserved, multifunctional regulator of forebrain development. Brain Res. 1705, 1–14 (2019).

25. L. L. Bruce, “Evolution of the Nervous System in Reptiles” in Evolution of Nervous Systems (2007)vol. 2, pp. 125–156.

26. B. Murillo, N. Ruiz-Reig, M. Herrera, A. Fairén, E. Herrera, Zic2 Controls the Migration of Specific Neuronal Populations in the Developing Forebrain. J. Neurosci. 35, 11266–11280 (2015).

27. M. Shanahan, V. P. Bingman, T. Shimizu, M. Wild, O. Güntürkün, Large-scale network organization in the avian forebrain: a connectivity matrix and theoretical analysis. Front. Comput. Neurosci. 7 (2013).

28. M. A. Tosches, From Cell Types to an Integrated Understanding of Brain Evolution: The Case of the Cerebral Cortex. Httpsdoiorg101146annurev-Cellbio-120319-112654 37 (2021).

29. S. O’Mara, The subiculum: what it does, what it might do, and what neuroanatomy has yet to tell us. J. Anat. 207, 271–282 (2005).

30. J. m. Wild, M. n. Williams, Rostral Wulst in passerine birds. I. Origin, course, and terminations of an avian pyramidal tract. J. Comp. Neurol. 416, 429–450 (2000).

31. E. D. Jarvis, O. Güntürkün, L. Bruce, A. Csillag, H. Karten, W. Kuenzel, L. Medina, G. Paxinos, D. J. Perkel, T. Shimizu, G. Striedter, J. M. Wild, G. F. Ball, J. Dugas-Ford, S. E. Durand, G. E. Hough, S. Husband, L. Kubikova, D. W. Lee, C. V. Mello, A. Powers, C. Siang, T. V. Smulders, K. Wada, S. A. White, K. Yamamoto, J. Yu, A. Reiner, A. B. Butler, Avian brains and a new understanding of vertebrate brain evolution. Nat. Rev. Neurosci. 6, 151–159 (2005).

32. L. Puelles, Current status of the hypothesis of a claustro-insular homolog in sauropsids. Brain. Behav. Evol., doi: 10.1159/000520742 (2021).

33. S. D. Briscoe, C. B. Albertin, J. J. Rowell, C. W. Ragsdale, Neocortical Association Cell Types in the Forebrain of Birds and Alligators. Curr. Biol. 28, 686–696.e6 (2018).

34. G. Gedman, B. Haase, G. Durieux, M. T. Biegler, O. Fedrigo, E. D. Jarvis, As above, so below: Whole transcriptome profiling demonstrates strong molecular similarities between avian dorsal and ventral pallial subdivisions. J. Comp. Neurol. 529, 3222–3246 (2021).

35. L. Puelles, M. Martinez-de-la-Torre, S. Martinez, C. Watson, G. Paxinos, The Chick Brain in Stereotaxic Coordinates and Alternate Stains (Academic Press, 2019).

36. H. Peng, P. Xie, L. Liu, X. Kuang, Y. Wang, L. Qu, H. Gong, S. Jiang, A. Li, Z. Ruan, L. Ding, Z. Yao, C. Chen, M. Chen, T. L. Daigle, R. Dalley, Z. Ding, Y. Duan, A. Feiner, P. He, C. Hill, K. E. Hirokawa, G. Hong, L. Huang, S. Kebede, H.-C. Kuo, R. Larsen, P. Lesnar, L. Li, Q. Li, X. Li, Y. Li, Y. Li, A. Liu, D. Lu, S. Mok, L. Ng, T. N. Nguyen, Q. Ouyang, J. Pan, E. Shen, Y. Song, S. M. Sunkin, B. Tasic, M. B. Veldman, W. Wakeman, W. Wan, P. Wang, Q. Wang, T. Wang, Y. Wang, F. Xiong, W. Xiong, W. Xu, M. Ye, L. Yin, Y. Yu, J. Yuan, J. Yuan, Z. Yun, S. Zeng, S. Zhang, S. Zhao, Z. Zhao, Z. Zhou, Z. J. Huang, L. Esposito, M. J. Hawrylycz, S. A. Sorensen, X. W. Yang, Y. Zheng, Z. Gu, W. Xie, C. Koch, Q. Luo, J. A. Harris, Y. Wang, H. Zeng, Morphological diversity of single neurons in molecularly defined cell types. Nat. 2021 5987879 598, 174–181 (2021).

37. Y. Atoji, J. M. Wild, Afferent and efferent projections of the mesopallium in the pigeon (Columba livia). J. Comp. Neurol. 520, 717–741 (2012).

38. I. Bystron, C. Blakemore, P. Rakic, Development of the human cerebral cortex: Boulder Committee revisited. Nat. Rev. Neurosci. 9, 110–122 (2008).

39. W. Z. Wang, F. M. Oeschger, J. F. Montiel, F. García-Moreno, A. Hoerder-Suabedissen, L. Krubitzer, C. J. Ek, N. R. Saunders, K. Reim, A. Villalón, Z. Molnár, Comparative Aspects of Subplate Zone Studied with Gene Expression in Sauropsids and Mammals. Cereb. Cortex 21, 2187–2203 (2011).

40. H. J. Karten, The Organization of the Avian Telencephalon and Some Speculations on the Phylogeny of the Amniote Telencephalon. Ann. N. Y. Acad. Sci. 167, 164–179 (1969).

41. S. Nagappan, K. M. Franks, Parallel processing by distinct classes of principal neurons in the olfactory cortex. eLife 10, e73668 (2021).

42. T. Nomura, E. I. Izawa, Avian brains: Insights from development, behaviors and evolution. Dev. Growth Differ. 59, 244–257 (2017).

43. Q. Tong, C. E. Romanini, V. Exadaktylos, C. Bahr, D. Berckmans, H. Bergoug, N. Eterradossi, N. Roulston, R. Verhelst, I. M. McGonnell, T. Demmers, Embryonic development and the physiological factors that coordinate hatching in domestic chickens. Poult. Sci. 92, 620–628 (2013).

44. L. Haghverdi, M. Büttner, F. A. Wolf, F. Buettner, F. J. Theis, Diffusion pseudotime robustly reconstructs lineage branching. Nat. Methods 2016 1310 13, 845–848 (2016).

45. C. Qiu, J. Cao, B. K. Martin, T. Li, I. C. Welsh, S. Srivatsan, X. Huang, D. Calderon, W. S. Noble, C. M. Disteche, S. A. Murray, M. Spielmann, C. B. Moens, C. Trapnell, J. Shendure, Systematic reconstruction of cellular trajectories across mouse embryogenesis. Nat. Genet. 2022 543 54, 328–341 (2022).

46. M. Cardoso-Moreira, J. Halbert, D. Valloton, B. Velten, C. Chen, Y. Shao, A. Liechti, K. Ascenção, C. Rummel, S. Ovchinnikova, P. V. Mazin, I. Xenarios, K. Harshman, M. Mort, D. N. Cooper, C. Sandi, M. J. Soares, P. G. Ferreira, S. Afonso, M. Carneiro, J. M. A. Turner, J. L. VandeBerg, A. Fallahshahroudi, P. Jensen, R. Behr, S. Lisgo, S. Lindsay, P. Khaitovich, W. Huber, J. Baker, S. Anders, Y. E. Zhang, H. Kaessmann, Gene expression across mammalian organ development. Nature 571, 505–509 (2019).

47. G. La Manno, K. Siletti, A. Furlan, D. Gyllborg, E. Vinsland, A. Mossi Albiach, C. Mattsson Langseth, I. Khven, A. R. Lederer, L. M. Dratva, A. Johnsson, M. Nilsson, P. Lönnerberg, S. Linnarsson, Molecular architecture of the developing mouse brain. Nat. 2021, 1–5 (2021).

48. T. Nomura, C. Ohtaka-Maruyama, W. Yamashita, Y. Wakamatsu, Y. Murakami, F. Calegari, K. Suzuki, H. Gotoh, K. Ono, The evolution of basal progenitors in the developing non-mammalian brain. Development 143, 66–74 (2016).

49. A. Cárdenas, A. Villalba, C. de Juan Romero, E. Picó, C. Kyrousi, A. C. Tzika, M. Tessier-Lavigne, L. Ma, M. Drukker, S. Cappello, V. Borrell, Evolution of Cortical Neurogenesis in Amniotes Controlled by Robo Signaling Levels. Cell 174, 590–606.e21 (2018).

50. J. Woych, A. O. Gurrola, A. Deryckere, E. C. B. Jaeger, E. Gumnit, G. Merello, J. Gu, A. J. Araus, N. D. Leigh, M. Yun, A. Simon, M. A. Tosches, Cell-type profiling in salamanders identifies innovations in vertebrate forebrain evolution. Science 377 (2022).

51. C. C. Chen, C. M. Winkler, A. R. Pfenning, E. D. Jarvis, Molecular profiling of the developing avian telencephalon: Regional timing and brain subdivision continuities. J. Comp. Neurol. 521, 3666–3701 (2013).

52. R. Remedios, D. Huilgol, B. Saha, P. Hari, L. Bhatnagar, T. Kowalczyk, R. F. Hevner, Y. Suda, S. Aizawa, T. Ohshima, A. Stoykova, S. Tole, A stream of cells migrating from the caudal telencephalon reveals a link between the amygdala and neocortex. Nat. Neurosci. 10, 1141–1150 (2007).

53. L. Medina, A. Abellán, E. Desfilis, Evolving Views on the Pallium. Brain. Behav. Evol., 1–19 (2021).

54. F. García-Moreno, Z. Molnár, Variations of telencephalic development that paved the way for neocortical evolution. Prog. Neurobiol. 194, 101865 (2020).

55. H. J. Karten, W. Hodos, W. J. H. Nauta, A. M. Revzin, Neural connections of the “visual wulst” of the avian telencephalon. Experimental studies in the pigeon (Columba livia) and owl (Speotyto cunicularia). J. Comp. Neurol. 150, 253–277 (1973).

56. J. M. Wild, The avian somatosensory system. I. Primary spinal afferent input to the spinal cord and brainstem in the pigeon (Columba livia). J. Comp. Neurol. 240, 377–395 (1985).

57. J. Dugas-Ford, J. J. Rowell, C. W. Ragsdale, Cell-type homologies and the origins of the neocortex. Proc. Natl. Acad. Sci. U. S. A. 109, 16974–16979 (2012).

58. A. Hoerder-Suabedissen, Z. Molnár, Development, evolution and pathology of neocortical subplate neurons. Nat. Rev. Neurosci. 16, 133–146 (2015).

59. H. Bruguier, R. Suarez, P. Manger, A. Hoerder-Suabedissen, A. M. Shelton, D. K. Oliver, A. M. Packer, J. L. Ferran, F. García-Moreno, L. Puelles, Z. Molnár, In search of common developmental and evolutionary origin of the claustrum and subplate. J. Comp. Neurol. 528, 2956–2977 (2020).

