## Supplementary_material for "Developmental origins and evolution of pallial cell types and structures in birds"

Science
AAAS

### Supplementary Materials for

#### Developmental origins and evolution of pallial cell types and structures in birds

**Authors:** Bastienne Zaremba[1]*, Amir Fallahshahroudi[1,2], Céline Schneider[1], Julia Schmidt[1], Ioannis Sarropoulos[1,3], Evgeny Leushkin[1,4], Bianka Berki[5], Enya Van Poucke[6], Per Jensen[6], Rodrigo Senovilla-Ganzo[7], Francisca Hervas-Sotomayor[1,8], Nils Trost[1], Francesco Lamanna[1], Mari Sepp[1], Fernando García-Moreno[7,9,10]*†, Henrik Kaessmann[1]*†

**Affiliations:**

[1]Center for Molecular Biology (ZMBH), DKFZ-ZMBH Alliance, Heidelberg University, Heidelberg, Germany.

[2]Current affiliation: Department of Medical Biochemistry and Microbiology, Biomedical Center (BMC), Uppsala University, Uppsala, Sweden.

[3]Current affiliation: Wellcome Sanger Institute, Wellcome Genome Campus, Hinxton, Cambridge, UK.

[4]Current affiliation: LOEWE Centre for Translational Biodiversity Genomics, 60325 Frankfurt, Germany.

[5]Deep Sequencing Core Facility, CellNetworks Excellence Cluster, Heidelberg University, Im Neuenheimer Feld 267, D-69120, Heidelberg, Germany.

[6]IFM Biology, AVIAN Behavioural Genomics and Physiology group, Linköping University, Sweden.

[7]Achucarro Basque Center for Neuroscience, Scientific Park of the University of the Basque Country (UPV/EHU), Leioa, Spain.

[8]Current affiliation: INRAE, LPGP, Rennes, France.

[9]Department of Neuroscience, Faculty of Medicine and Odontology, UPV/EHU, Leioa, Spain.

[10]IKERBASQUE Foundation, Bilbao, Spain.

†These authors jointly supervised the work.

**The PDF file includes:**

Materials and Methods
Figs. S1 to S22
Tables S1 to S6
References

#### Materials and Methods

Sample collection

Adult chicken samples

Most samples from adult animals (*Gallus gallus*, aged 1-2 years, breed: red junglefowl) were received from Linköping university. The study was approved by the Linköping Council for Ethical Licensing of Animal Experiments, license number 288-2019. The animals were kept in pens under a 12:12 h dark:light schedule and provided with food and water ad libitum. In addition, we were provided with heads of healthy domestic chickens (*Gallus gallus*, aged ca. 2 years, breed: Lohmann white) by a local farm. Brains were extracted and either dissected in ice-cold PBS and snap frozen in liquid nitrogen, or embedded in OCT sectioning medium and frozen on dry ice to preserve for cryosectioning.

Developing chicken samples

All animal experiments were approved by a local ethical review committee and conducted in accordance with personal and project licenses in compliance with the current normative standards of the European Union (Directive 2010/63/EU) and the Spanish Government (Royal Decrees 1201/2005 and 53/2013, Law 32/107).

Fertilized chicken eggs (*Gallus gallus*) were purchased from Granja Santa Isabel and incubated at 37.5 °C in a humidified atmosphere until the required developmental stage. The day when eggs were incubated was considered embryonic day (E)0.

Mouse samples

All animal procedures were performed in compliance with national and international ethical guidelines for the care and use of laboratory animals, and were approved by the local animal welfare authorities: Heidelberg University Interfaculty Biomedical Research Facility (T-23/19, T-28/21). The animals (*Mus musculus*, RjOrl:SWISS) were housed under a 12h/12h dark/light cycle in a temperature (20-24 °C) and humidity (40-65%) controlled room with ad libitum access to food and water. Mice were sacrificed by cervical dislocation on post-natal day 56. Dissections of the frontal isocortex include the anterior cingulate area, the prelimbic area, the orbital area, the infralimbic area, primary and secondary motor areas and the agranular insular area up to the start of the corpus callosum. Dissections of the ventral and lateral pallial derived structures include the insular cortex, the claustrum, the endopiriform nucleus, the piriform cortex and the amygdala. The positions of these regions were determined according to the Allen Mouse Brain Reference Atlas (2004).

Green anole samples

All animal procedures were performed in compliance with national and international ethical guidelines for the use of laboratory animals, and were approved by the local animal welfare authorities: Heidelberg University Interfaculty Biomedical Research Facility (T-04/21).

Wild animals (*Anolis carolinensis*) were obtained from a commercial supplier (Interaquaristik.de) and were kept in temporary cages before they were anesthetized by intraperitoneal injection of Narcoren (16 g/100 ml Sodium – Pentobarbital, Boehringer Ingelheim; 200 mg/kg body weight) and subsequently decapitated. Brains were extracted and dissected in ice-cold PBS.

Nuclei preparation

Nuclei were isolated from fresh frozen tissue according to a protocol adapted from (*60*). Briefly, the tissue was homogenized by trituration on ice in 250 mM sucrose, 25 mM KCl, 5 mM $MgCl_2$, 10 mM Tris HCl (pH 8), 0.1 % IGEPAL, 1 µM DTT, 0.4 U/µl, Murine RNase Inhibitor (New England Biolabs), 0.2 U/µl SUPERas-In (Thermo Fisher) and Hoechst DNA dye. After 5 min of incubation, remaining unlysed tissue was pelleted and removed by centrifugation at 100 g for 1 min. Nuclei in the collected supernatant were pelleted at 400 g for 5 min. Then, nuclei were washed once in homogenization buffer before they were resuspended in 1x PBS and filtered using Flowmi cell filters (pore size 40 µm; Merck). For adult samples used in single nuclei RNA experiments, fluorescence activated cell sorting (BD FACSAria ii, 85 µm nozzle, BD Biosciences) was used to separate single nuclei from remaining debris and aggregates according to forward and sideward scatter properties, as well as DNA content based on Hoechst signal. Due to the limited volume allowed as input, samples used for single nuclei multiome experiments were not sorted to keep a high concentration of nuclei in the solution. Following sorting or final resuspension, nuclei were counted on Countess II FL Automated Cell Counter (Thermo Fisher). 17 000 nuclei were employed for single nuclei RNA and multiome sequencing experiments.

Library preparation and sequencing

Single nuclei RNA and multiome sequencing experiments were performed using the 10x Chromium Single Cell 3' v3 and v3.1 Gene Expression Kit (10x Genomics) and the 10x Chromium single cell multiome ATAC + gene expression kit (v1), respectively, following the manufacturer's instructions. Quantification and quality control of libraries was performed using a Qubit Fluorometer and the High Sensitivity NGS Fragment Kit for Agilent's Fragment Analyzer (Agilent, Santa Clara, CA, USA). V3 and v3.1 gene expression libraries were sequenced on an Illumina NextSeq 500/550 (Illumina) using the High Output Kit v2.5 (75 Cycles; Illumina) with paired end sequencing and 28 cycles for Read 1, 56 cycles for Read 2 and 8 cycles for i7 index to a depth of ca. 200 million reads.

Multiome libraries were sequenced on NextSeq 500/550 using the High Output Kit v2.5 (150 Cycles; Illumina) with paired end sequencing. A custom recipe, provided by Illumina, was used to sequence the multiome ATAC libraries to specify 8 dark cycles at i5 index. Multiome RNA libraries were sequenced with 28 cycles for Read 1, 90 cycles for Read 2 and 10 cycles for both indices to a depth of ca. 200 million reads. For Multiome ATAC libraries the read lengths were 50 cycles for Read 1 (DNA), 8 cycles for i7 index (sample index), 8 dark cycles followed by 16 cycles for i5 index (Barcode), and 50 cycles for Read 2 (DNA).

Genome annotation and alignment

For chicken samples, we used genome assembly galGal5 and a custom annotation. Our custom annotation is based on the reference genome annotation from Ensembl release 87, which was extended using chicken brain 3'-RNA-sequencing data as previously described in (*61*). Assembly AnoCar2.0 and the ensembl reference genome annotation (release 104) was used for green anole samples. For mouse samples we used the ensembl reference genome (release 91). We used STAR aligner (v 2.7.10a) to produce references for all species and used the STARsolo mode (--soloType CB_UMI_Simple) to align reads to references (--clipAdapterType CellRanger4; --outFilterScoreMin 20; --soloCBmatchWLtype 1MM_multi_Nbase_pseudocounts; --soloUMIfiltering MultiGeneUMI_CR; --soloUMIdedup 1MM_CR; --soloMultiMappers EM) (*62*).

Data quality control – snRNA-seq

Nuclei-containing barcodes were identified based on the number of UMIs and the fraction of intronic reads. Doublets were identified and removed in each library using ScDblFinder (v 1.12; settings: dbr = 0.1; dbr.sd = 1) (*63*). Subsequently, barcodes with a high fraction of mitochondrial reads (chicken adult > 0.03, developmental chicken > 0.1, mouse > 0.05, green anole > 0.2) were removed. Each library was then clustered and any low-quality clusters (lower average number of UMIs, higher average fraction of mitochondrial reads, lower average fraction of intronic reads, few differentially expressed genes and spread appearance on the UMAP) were removed.

Data integration snRNA-seq

Whole datasets, as well as subsets, were integrated as follows. Each library was individually normalized and highly variable genes were identified using the SCTransform function implemented in Seurat (v 4.3.0) (*64*) with residual variance cutoff (variable.features.rv.th) set to 1.4. The fraction of mitochondrial reads was regressed out during normalization. Subsequently, libraries were merged using Seurat's merge function. For chicken datasets, where we had profiled more than two individuals, we chose the union of variable genes, which were called variable in at least two individuals, as the set of variable genes for the merged dataset. Genes whose expression could not be detected in all individuals but were part of the set of variable genes, were set to 0 scaled expression in all libraries, in which expression could not be detected, so they were not lost in the merged object.

For mouse and green anole datasets we called variable genes independently for each dissection (isocortex or ventro-lateral pallium derivatives, and cortex or DVR, respectively), if they were variable in at least one individual and were expressed in the other individual (detected in at least 5 cells). We then used the union of these dissection specific variable genes as the set of variable genes for the whole dataset. As described above, genes, whose expression could not be detected in all individuals, i.e. across dissections, were set to 0 scaled expression.

Only for the glutamatergic lineage in the developing chick pallium we used batch integration to achieve a better integration across developmental stages. Specifically, we ran Harmony integration (v 0.1.1) (*65*) based on 40 principal components that were computed on the glutamatergic lineage dataset, which was merged as described above, before we generated the UMAP projections shown in Fig. 6 and ran diffusion pseudotime (see below).

Clustering snRNA- seq

In all merged datasets, we identified cell classes according to the expression of major known marker genes (Fig 1B). Cell populations in the non-neuronal class were only broadly annotated and not clustered further. In the adult chicken dataset, neuronal cell classes were subset and again split into broad groups, which roughly correspond to the annotated subclasses, although the exact subclass annotation was determined post-hoc. These groups were merged, renormalized and integrated as described above, and subsequently clustered using Louvain clustering at different resolutions. The highest resolution was chosen so the dataset was clearly overclustered. The cluster identity of cells at different resolutions was then used to construct a dendrogram using MRtree, which allows to build cluster hierarchy based on flat clustering obtained for multiple resolutions (*66*). Differentially expressed genes were called between clusters representing nodes of the dendrogram using Seurat's FindMarkers function (min.pct = 0.3; logfc.threshold = 0.2) to decide whether these clusters should be merged. Clusters were merged if we could not identify robust differential expression of any transcription factors or genes related to neuronal function. The green anole and adult mouse datasets were clustered similarly, only that neuronal cell classes were not further split into groups due to overall lower cell numbers. The developing chick dataset was only broadly clustered to define major cell populations shown in Fig. 5B.

Identification of supertypes and subclasses in the adult chicken pallium

Supertype and subclass identity of individual clusters was determined according to their position in a cluster dendrogram, which was constructed for the complete dataset (Fig. 1B) or for neuronal cell classes (Fig. 2A, Fig. 3A), and according to low resolution Louvain clustering. To construct cluster dendrograms, we calculated the average expression per cluster of the union of the top 5000 expressed genes per cluster in the dataset or cell class, respectively, using Seurat's AverageExpression function. Gene expression was correlated across clusters using Spearman correlation and the resulting correlation matrix was used as an input for hierarchical clustering using pvclust (method.hclust = "ward.D2", nboot = 1000) (*67*). For the dendrogram of excitatory neuron clusters we excluded clusters, which likely contained cells inadvertently dissected from the thalamus (TCF7L2+) (*11*).

Annotation of adult mouse data

Since we expected very little overlap between cell populations from different dissections (frontal isocortex or ventro-lateral pallial derivatives including insular cortex, claustrum, endopiriform nucleus, piriform cortex and amygdala) we analyzed dissections separately. Libraries from each dissection were integrated and clustered as detailed above. Subsequently we used CCA integration and label transfer implemented in Seurat (*21*) to transfer labels from different subsets of external datasets to our data to aid with annotation (see below). Specifically, we used subsets of (*14*) to annotate our data for the frontal isocortex, since dissections overlap between the two datasets. To aid with annotation of our data for ventro-lateral pallial derivatives, we used different datasets which partly covered the regions we dissected (*14*, *16*). Especially cell populations of the piriform cortex were annotated using available in situ hybridization data (*17*) of identified marker genes, since this region was not covered in any external dataset. We named overlapping populations according to the supertype and subclass nomenclature introduced in (*14*) and only introduced new names for non-overlapping populations (Fig. 1E; fig. S7). To create the final adult mouse pallium dataset, we subsampled 300 cells per neuronal supertype from all profiled regions in (*14*) and integrated this subset with our datasets from both dissections.

Annotation of lizard data

Libraries from all dissections (cortex and DVR) were integrated and clustered as detailed above. Clusters were assigned a probable regional identity according to the expression of marker genes, whose *in situ* expression is known in other reptiles, and according to comparisons to available scRNA-seq data from the pallium of one turtle and another lizard species (*7*, *12*, *13*). For details on comparisons across species see below.

Annotation of developing mouse pallium

Data from (*47*) was subset to include only pallial progenitors and excitatory neurons according to the expression of known marker genes (fig. SX) and only cells from embryonic days 9 to 17 to ensure a roughly equal representation of all pallial lineages. The subset was integrated as described under data integration, followed by batch integration using Harmony (*65*) to produce the UMAPs shown in fig. S21. In order to annotate early neurons in more detail, we subset late intermediate progenitor cells and neurons and ran the same integration procedure before clustering to a high resolution. Clusters of early neurons were annotated based on the expression of known marker genes and spatial expression of identified marker genes in available *in situ* hybridization data (*17*). Radial glia and intermediate progenitor cells (IPCs) were not annotated in more detail.

<u>In situ sequencing (ISS)</u>

Samples embedded in OCT mounting medium were cryosectioned into 10 µm coronal and sagittal sections and stored at −80 °C until further use. Sections were processed for ISS using the High Sensitivity Library Preparation Kit from CARTANA AB (10x Genomics) (*9*). After fixation in 3.7% (v/v) paraformaldehyde in UltraPure distilled water for 10 min, sections were processed in SecureSeal hybridization chambers (Grace Bio-Labs) following the manufacturer's protocol. The mounted sections were shipped to CARTANA's facility (Solna, Sweden) for ISS. The 50 chosen profiled genes (fig. S3) represent marker genes with medium expression levels identified from the chicken adult dataset, as well as known markers from mammals. In order to assign identities to cells in the sections, segmentation of cells is needed. Therefore, we first segmented cells based on the DAPI image only, using an approach provided by CARTANA which is based on intensity thresholding and a watershed segmentation (*68*). We then used this segmentation as a prior distribution for baysor (v 0.5.1) (*69*), which allows to incorporate information from gene molecule positions as well as the DAPI image. Baysor was run with the following specifications: scale = 12, scale_std = 3, prior-segmentation-confidence = 0.2, min_molecules_per_cell = 3.

Following segmentation, we used Tangram (v1.0.4) (*10*) to map snRNA-seq data to tissue sections. As spatial data input to Tangram, we subset baysor output to cells mostly located in the pallium in order to mimic the dissections used for snRNA-seq and make the spatial and snRNA-seq data most comparable, since Tangram also takes abundances of populations into account. We then integrated the chosen subsets of cells in each section with the complete snRNA-seq dataset from the adult chicken using the cluster mode and all 50 spatially profiled genes resulting in a matrix containing mapping probabilities for each cell in the section to each identity in the snRNA-seq dataset. We separately mapped class as well as supertype labels to all cells in the sections. Cells were confidently assigned a supertype identity if cell class as well as supertype label assignment were in agreement (e.g. a cell with supertype identity Ex_CACNA1H_PROX1 should also be identified as an excitatory neuron), and the cell belonged to the top 75% of cells with the highest probability for this supertype.

<u>Visium</u>

Samples embedded in OCT mounting medium were cryosectioned into 10 µm coronal and sagittal sections and collected on Visium Spatial Gene Expression slides (10x Genomics). Slides were fixed in methanol and stained using Hematoxylin & Eosin staining as suggested by 10x genomics. Images were taken with an Olympus VS200 Slide Scanner (20X magnification) before slides were processed using the Visium Spatial Gene Expression Reagent Kit (10x) according to the manufacturer's instructions. Sections from adult samples were permeabilized for 18 min, embryonic day 19 sections for 16 min. cDNA and library concentrations were quantified using the Qubit Fluorometer with the DNA High Sensitivity kit. cDNA quality was assessed on

Bioanalyzer High Sensitivity DNA chips. Library quality was assessed using Agilent's TapeStation with the D1000 kit. Visium libraries were sequenced on an Illumina NextSeq500/550 using the High Output Kit v2.5 (75 Cycles) with paired end sequencing and 28 cycles for Read 1, 56 cycles for Read 2 and 8 cycles for i7 index to a depth of ca. 200 million reads.

Analysis of multiome data

We used our custom chicken genome annotation to generate references for cellranger-arc (v2.0.1) and ArchR (v1.0.2). Single nucleus multiome libraries were demultiplexed and mapped to the genome using cellranger-arc (v2.0.1). Barcodes corresponding to cells were identified based on the following four metrics: number of UMIs (for full-length transcripts), fraction of intronic reads, number of ATAC fragments and transcription start site (TSS) enrichment scores (as estimated by ArchR, v1.0.2). We used Gaussian mixture models (v5.4.7) with two groups to identify the set of barcodes with the highest values for each of the metrics. Only barcodes corresponding to the group with higher values in all four metrics, as well as having at least 40% of the median number of UMIs across putative cells in that sample, were considered as high-quality cells. We used scrublet (v0.2.3) and ArchR (v1.0.2) to estimate doublet scores for the gene expression and chromatin accessibility modalities, respectively. To jointly consider both metrics, we standardized doublet scores within each modality as Z-scores and additionally estimated a consensus doublet score by taking the mean of the two scores for each barcode. We then removed barcodes that ranked in the top 10% when considering the consensus doublet score or in the top 5% when considering the doublet score in either modality.

Following the identification of high-quality cells, we recalculated gene expression counts using STARsolo as described above to be consistent with the rest of our dataset. We then integrated the multiome gene expression data with the full snRNA-seq dataset using Harmony. We used the integrated embedding to identify the 20 nearest annotated neighbors for each unannotated cell. The most frequent cell type label of these 20 neighbors was assigned to the unannotated cell, whereas the fraction of neighbors with that label was used to estimate the confidence score of the annotation. We only considered cells with at least 15 (75%) of neighbors having the same label. Additionally, to increase our confidence in the annotation of glutamatergic neuron subtypes, we repeated this procedure in a subsetted embedding that only included glutamatergic neurons, and only retained cells with the same label in both annotations for downstream analyses. In total, this stringent annotation strategy retained 8,119 cells.

We next used ArchR (v1.0.2) to quantify the accessibility of these 8,119 cells across 500 bp genomic tiles. We applied iterative latent semantic indexing (LSI) with 5 iterations with increasing resolution (0.1, 0.2, 0.4, 0.8) on the genomic tile matrix to project the data on 100 dimensions. To facilitate the detection of cell type-specific as well as brain region-specific open chromatin peaks, we subsetted fragment files by cell type and brain region and provided them as input to MACS (v2.1.2) to call peaks in each group with the following parameters: -f BEDPE --nomodel --nolambda --extsize 150 --shift -75 --keep-dup all -q 0.05 --gsize 1.2e9 --call-summits. To ensure that we make the most out of our dataset to identify peaks at the highest possible granularity, while retaining power to discover peaks shared across broader groups, we repeated the grouping at all three levels of our hierarchical annotation. We then used ArchR's peak merging procedure based on iterative overlaps to merge the group-specific peak sets into a consensus peak annotation, identifying a total of 367,896 peaks. We imported this peak set into ArchR using addPeakSet and addPeakMatrix. We repeated the iterative LSI projection described above using the peak matrix, obtaining similar results.

To assess the effect of cell type- and brain region-specificity on chromatin accessibility profiles, we aggregated peak counts across cells from the same cell type and brain region, only considering groups with at least 40 cells. We then computed Spearman's correlations across groups, using peaks reaching at least 10 CPM in at least one group. The resulting correlation matrix was used as input for hierarchical clustering.

To identify differentially accessible regions (DARs) between cell types and brain regions, we used ArchR's function getMarkerFeatures, with a scale factor of 10,000, accounting for TSS enrichment bias and the number of fragments detected per cell, and only considering groups with at least 40 cells. Statistically significant DARs were filtered for FDR < 5% and a log fold-change of 1.25. For plotting, we scaled the pseudobulk (cell type x brain region) peak count matrix by sequencing depth (counts per million, CPM) and additionally scaled each peak by its maximum accessibility across pseudobulks.

<u>Supertypes shared across regions – dendrogram and differential gene expression</u>
We used snRNA-seq data from two individuals (one profiled using 10x Chromium Single Cell 3' v3.1 Gene Expression Kit, the other using 10x Chromium single cell multiome ATAC + gene expression kit) for which the pallium was dissected into anatomical regions, to construct a dendrogram of supertypes split by dissection. For one individual we pooled the "anterior DVR" and "posterior DVR" samples *in silico* to represent the DVR. Shared excitatory supertypes, with at least 35 cells from both DVR and Wulst dissections in both individuals, were identified. The datasets were subset to shared supertypes and cells were subsampled for roughly equal contribution from both individuals, with a maximum of 300 cells per supertype and dissection. If one individual had fewer than 100 cells per supertype and dissection, the other individual was subsampled to contain 100 cells. Subsequently, data from both individuals was merged and we calculated average genes expression per supertype and dissection across all expressed genes using Seurat's AvergeExpression function. Supertypes per dissection were correlated using Spearman correlation, and the resulting correlation matrix was hierarchically clustered using pvclust (method.hclust = "ward.D2", nboot = 1000) (*67*).
To identify differentially expressed genes between supertypes shared across regions, we used the same dataset as described above, but did not subsample cells before merging data from both individuals. Counts were aggregated per dissection and supertype using Seurat's AggregateExpression function and significantly (adjusted p-value <= 0.05) differentially expressed genes were identified using DESeq2 (v 1.38.3) (*70*) according to the standard workflow.

<u>Comparison across species</u>
We used three different methods to compare adult cell type populations across species, namely gene specificity index (GSI) correlation as described in (*7*), Seurat's CCA integration with label transfer and SAMap (v1.0.7) (*21*, *22*). To calculate the similarity score, scores resulting from each method were scaled from 0 to 1 and added, meaning the similarity score can range from 0 to 3. Since we did not identify any immature neuron populations in the adult mouse dataset, we

excluded these in other species in order to make datasets as comparable as possible. For each method and comparison, we determined the top reciprocal matches per cell type population by finding the overlap between the top five most similar populations for each cell type population as viewed from one species and the top five most similar populations as viewed from the other species.

Orthologous genes: For GSI correlation and label transfer, 1:1 orthologous genes were identified using OrthoFinder (v 2.5.4) (*71*). The genomes and annotations used for OrthoFinder are listed in Table S5. The standard SAMap workflow includes a different approach to identify orthologous genes.

Subsampling: To mitigate any differences in power to identify shared gene expression across species between different cell type populations within one species, we subsampled cells and UMIs within each species. Specifically, for label transfer and SAMap, cell populations containing more than the median cell number were subsampled to the median cell number. For GSI correlation, we additionally subsampled UMI counts per cell in large cell type populations (cell number > median cell number) because we saw that this approach is especially sensitive to differences in power, as large populations always showed higher correlations on average. Mouse datasets were subsampled so that cells from our own dataset were preferred over cells from (*14*) in shared populations.

Seurat CCA label transfer: For label transfer we identified all 1:1 orthologs, which could be detected in both species. We then subset each dataset to the chosen set of genes and ran the Seurat pipeline to map and annotate query datasets, using canonical correlation analysis (CCA) as the reduction method. To calculate one third of the overall similarity score (see above), we first calculated the fraction of cell type populations in one species predicted to belong to cell type populations in the other species. We then weighted this fraction by the average prediction score for each prediction. Values from one comparison (two species, glutamatergic and/or GABAergic neurons) were then scaled between 0 and 1.

GSI correlation: For GSI correlation, we first calculated the average expression of all genes per cell type population in the subsampled datasets of each species. We then subset the expression matrix to genes, which were robustly expressed in both species i.e. genes which showed an average expression of at least 5 UMIs summed across all cell type populations, and which were expressed by at least 2% of cells of at least one cell type population in both species in the complete dataset (not subsampled). We then calculated the gene specificity indices according to (*7*) and correlated cell type populations based on the calculated indices using spearman correlation. We scaled the resulting correlation between 0 and 1.

SAMap: We used datasets subsampled as detailed above to run the standard SAMap workflow. When constructing the SAMap object, we set keys to the highest resolution defined in the dataset, which was usually clusters. When running the SAMap algorithm we set neigh_from_keys to the levels of annotation we wanted to compare. Resulting cell type mapping scores were scaled from 0 to 1.

Comparison of developmental datasets: Due to the continuous nature of the datasets for the developing chicken and murine pallium we did not subsample cell type populations for the comparison. Since GSI correlation is particularly sensitive to differences in cell number and number of detected genes, we decided to only use SAMap and Seurat CCA label transfer for this comparison, meaning the maximum similarity score is 2.

<u>Identification of excitatory neuron lineages in the developing chicken pallium</u>

To identify different lineages of early excitatory neurons in the developing chicken pallium, we adapted an approach previously established to identify cellular lineages in mouse embryogenesis (*45*). Therefore, cells of the excitatory lineage were subset and split by developmental stage. Data for each stage was normalized and integrated separately as described under Data integration snRNA-seq with minor adaptations. The residual variance cutoff was set to 1.3 and variable genes of the merged object encompassed all variable genes which were called variable in at least one individual and detected in at least two individuals. Integrated datasets per stage were clustered to high resolutions and subsequently integrated with the previous and following stages using Harmony integration (*65*) to calculate a shared embedding. We then used a k-nearest-neighbor (k-NN) heuristic on the shared embeddings, as described in (*45*), to connect "pseudoancestor" and "pseudodescendant" populations across stages with weighted edges resulting in a weighted graph. Because the weighted graph was too complex to examine visually, we filtered out edges with a weight below 0.1 and applied leiden clustering implemented in igraph (v 1.4.2) with different resolution parameters (n_iterations = 4, objective_function = 'modularity') (*72*) to identify potential lineages of early neurons. We identified four major lineages which encompassed most neurons in the dataset. Some populations of the earliest and most mature neurons formed separate leiden communities at higher resolutions likely because these are highly similar across few stages and are connected to later or previous stages, respectively, by low weight edges only. To assign these largely disconnected populations to lineages, we identified marker genes of the four major identified lineages using Seurat's FindMarkers function and different subsets of cells per lineage with high or low pseudotime values and evaluated the expression of these genes in populations with unidentified lineage identity. We could assign a clear lineage identity to most cell populations, although some could not be unambiguously assigned and remained "unknown" or were named after their two most closely related lineages.

Pseudotime

We calculated diffusion-based pseudotime (*44*) implemented in scanpy (v 1.9.3) for the excitatory neuron lineage in the developing chicken pallium as described in (*73*). Briefly, we used Harmony-corrected components to construct a graph based on the 20 nearest neighbors of each cell. A diffusion map was computed based on the neighborhood graph and was used as input for pseudotime estimated with zero branchings. The root of the pseudotime was specified as a random cell belonging to the earliest developmental stage (E6) and to a cluster of non-cycling radial glia (Ex_PCs).

Correlation of early neuron lineages

The four major lineages of early neurons identified in the developing chicken pallium (HyperP, medial/ArcoP, MesoP and NidoP) were split into 6 bins according to pseudotime. Pseudotime - lineage bins were correlated using Pearson correlation of average expression across variable genes identified in the whole excitatory lineage as described under Data integration snRNA-seq.

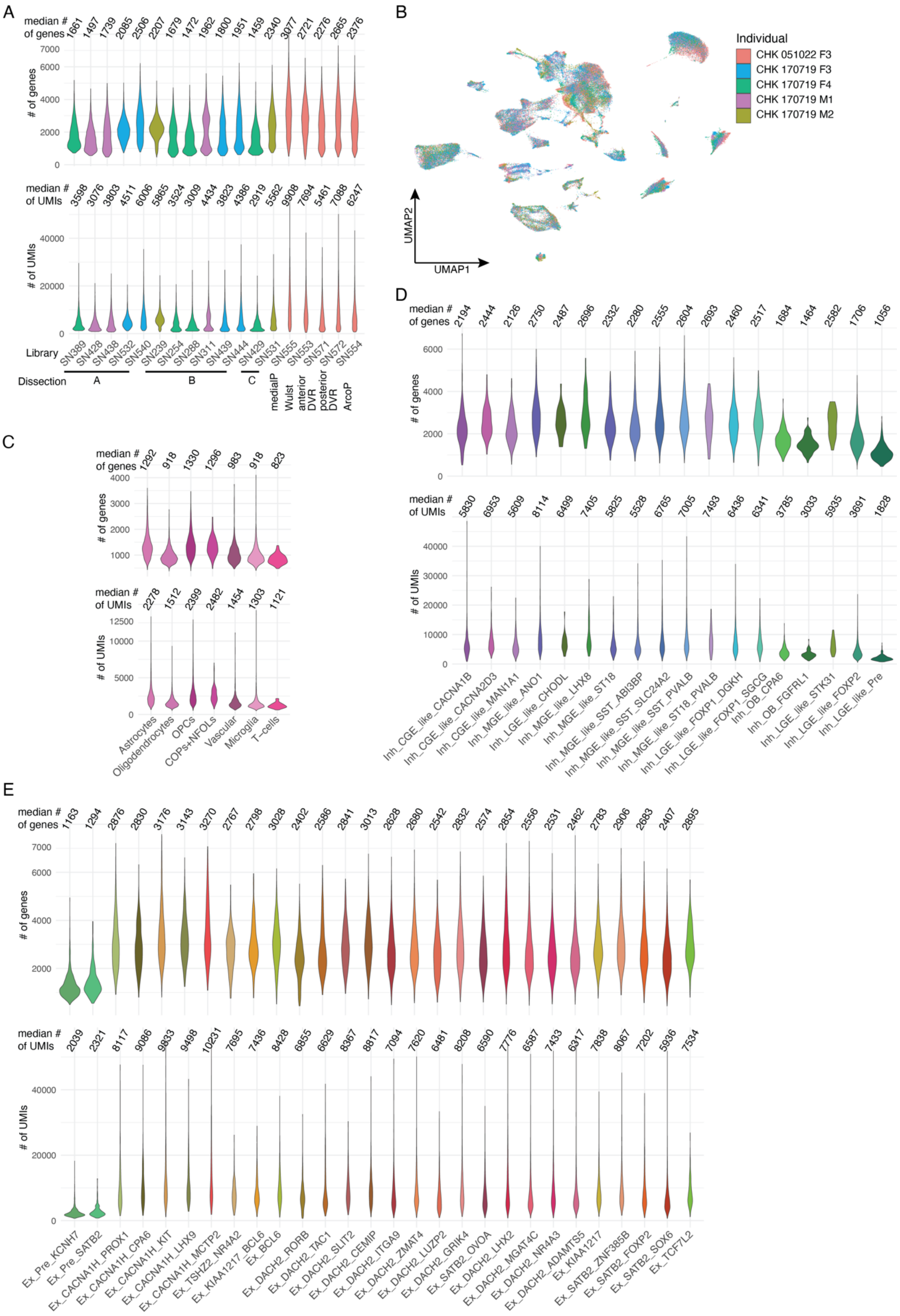
A
median # of genes
1661 1497 1739 2085 2506 2207 1679 1472 1962 1800 1951 1459 2340 3077 2721 2276 2665 2376
### of genes
median # of UMIs
3598 3076 3803 4511 6006 5865 3524 3009 4434 3823 4386 2919 5562 9908 7694 5461 7088 6247
### of UMIs
Library
SN389 SN428 SN438 SN532 SN540 SN239 SN254 SN288 SN311 SN439 SN444 SN429 SN531 SN555 SN553 SN571 SN572 SN554
Dissection
A
B
C
medialP
Wulst
anterior DVR
posterior DVR
ArcoP
B
Individual
CHK 051022 F3
CHK 170719 F3
CHK 170719 F4
CHK 170719 M1
CHK 170719 M2
UMAP2
UMAP1
C
median # of genes
1292 918 1330 1296 983 918 823
### of genes
median # of UMIs
2278 1512 2399 2482 1454 1303 1121
### of UMIs
Astrocytes
Oligodendrocytes
OPCs
COPs+NFOLs
Vascular
Microglia
T-cells
D
median # of genes
2194 2444 2126 2750 2487 2696 2332 2280 2555 2604 2693 2460 2517 1684 1464 2582 1706 1056
### of genes
median # of UMIs
5830 6953 5609 8114 6499 7405 5825 5528 6765 7005 7493 6436 6341 3785 3033 5935 3691 1828
### of UMIs
Inh_CGE_like_CACNA1B
Inh_CGE_like_CACNA2D3
Inh_CGE_like_MAN1A1
Inh_MGE_like_ANO1
Inh_LGE_like_CHODL
Inh_MGE_like_LHX8
Inh_MGE_like_ST18
Inh_MGE_like_SST_ABI3BP
Inh_MGE_like_SST_SLC24A2
Inh_MGE_like_SST_PVALB
Inh_MGE_like_ST18_PVALB
Inh_LGE_like_FOXP1_DGKH
Inh_LGE_like_FOXP1_SGCG
Inh_OB_CPA6
Inh_OB_FGFRL1
Inh_LGE_like_STK31
Inh_LGE_like_FOXP2
Inh_LGE_like_Pre
E
median # of genes
1163 1294 2876 2830 3176 3143 3270 2767 2798 3028 2402 2586 2841 3013 2628 2680 2542 2832 2574 2854 2556 2531 2462 2783 2906 2683 2407 2895
### of genes
median # of UMIs
2039 2321 8117 9086 9833 9498 10231 7695 7436 8428 6855 6629 8367 8817 7094 7620 6481 8208 6590 7776 6587 7433 6317 7838 8067 7202 5936 7534
### of UMIs
Ex_Pre_KCNH7
Ex_Pre_SATB2
Ex_CACNA1H_PROX1
Ex_CACNA1H_CPA6
Ex_CACNA1H_KIT
Ex_CACNA1H_LHX9
Ex_CACNA1H_MCTP2
Ex_TSHZ2_NR4A2
Ex_KIAA1217_BCL6
Ex_BCL6
Ex_DACH2_RORB
Ex_DACH2_TAC1
Ex_DACH2_SLIT2
Ex_DACH2_CEMIP
Ex_DACH2_ITGA9
Ex_DACH2_ZMAT4
Ex_DACH2_LUZP2
Ex_DACH2_GRIK4
Ex_SATB2_OVOA
Ex_DACH2_LHX2
Ex_DACH2_MGAT4C
Ex_DACH2_NR4A3
Ex_DACH2_ADAMTS5
Ex_KIAA1217
Ex_SATB2_ZNF385B
Ex_SATB2_FOXP2
Ex_SATB2_SOX6
Ex_TCF7L2

**Fig. S1. Adult chicken pallium single nucleus RNA-sequencing statistics**
(A) Number of UMI counts (bottom) and detected genes (top) per snRNA-seq library after selection of high-quality cells colored by sampled individuals. (B) UMAP of chicken pallium dataset colored by sampled individuals. (C) Number of UMI counts (bottom) and detected genes (top) per cell population of non-neuronal class (C), per inhibitory supertype (D) or excitatory supertype (E) colored cell population/supertype.

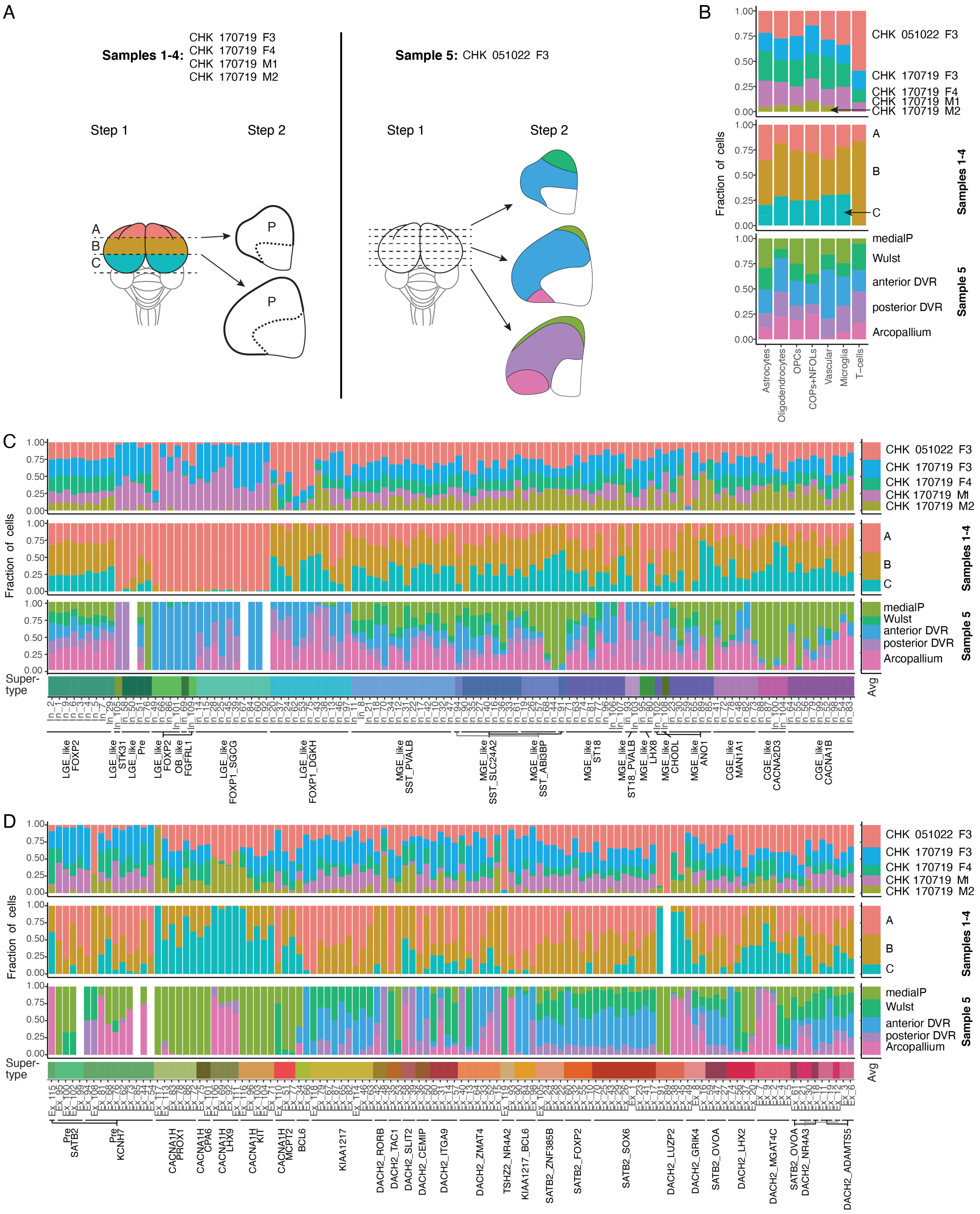


**Fig. S2. Adult chicken dissection statistics**

(A) Dissection scheme of adult chicken individuals. (B) Fraction of cells per cell non-neuronal cell population (B), inhibitory supertype (C) or excitatory supertype (D) stemming from sampled individuals (top), broad pallial dissections (individuals 1-4) (mid) or pallial dissections of anatomical regions (individual 5) (bottom). medialP, medial Pallium.

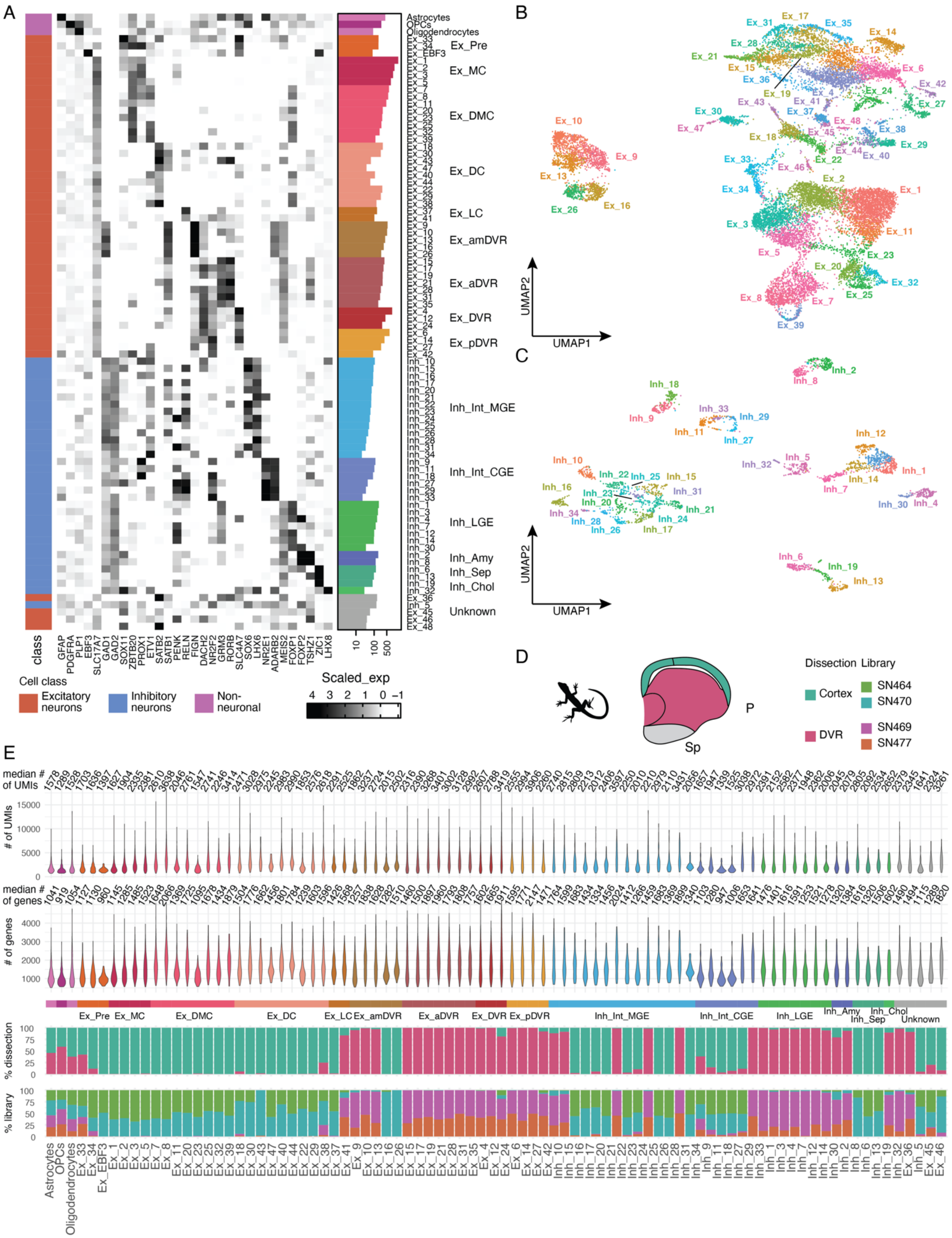

**Fig. S3. Lizard pallium single nucleus RNA-sequencing dataset**
(A) Heatmap of selected marker gene expression across 86 clusters in snRNA-seq-based lizard pallium atlas. Barplots of cell numbers per cluster on the right are coloured by regional annotation as indicated by text labels. UMAP of excitatory (B) and

inhibitory (C) neurons colored by and labeled with cluster annotation. (D) Illustration of dissections and snRNA-seq libraries per dissection. (E) Number of UMI counts and detected genes (top) per cluster and fraction of cells per cluster stemming from different dissections and snRNA-seq libraries (bottom).

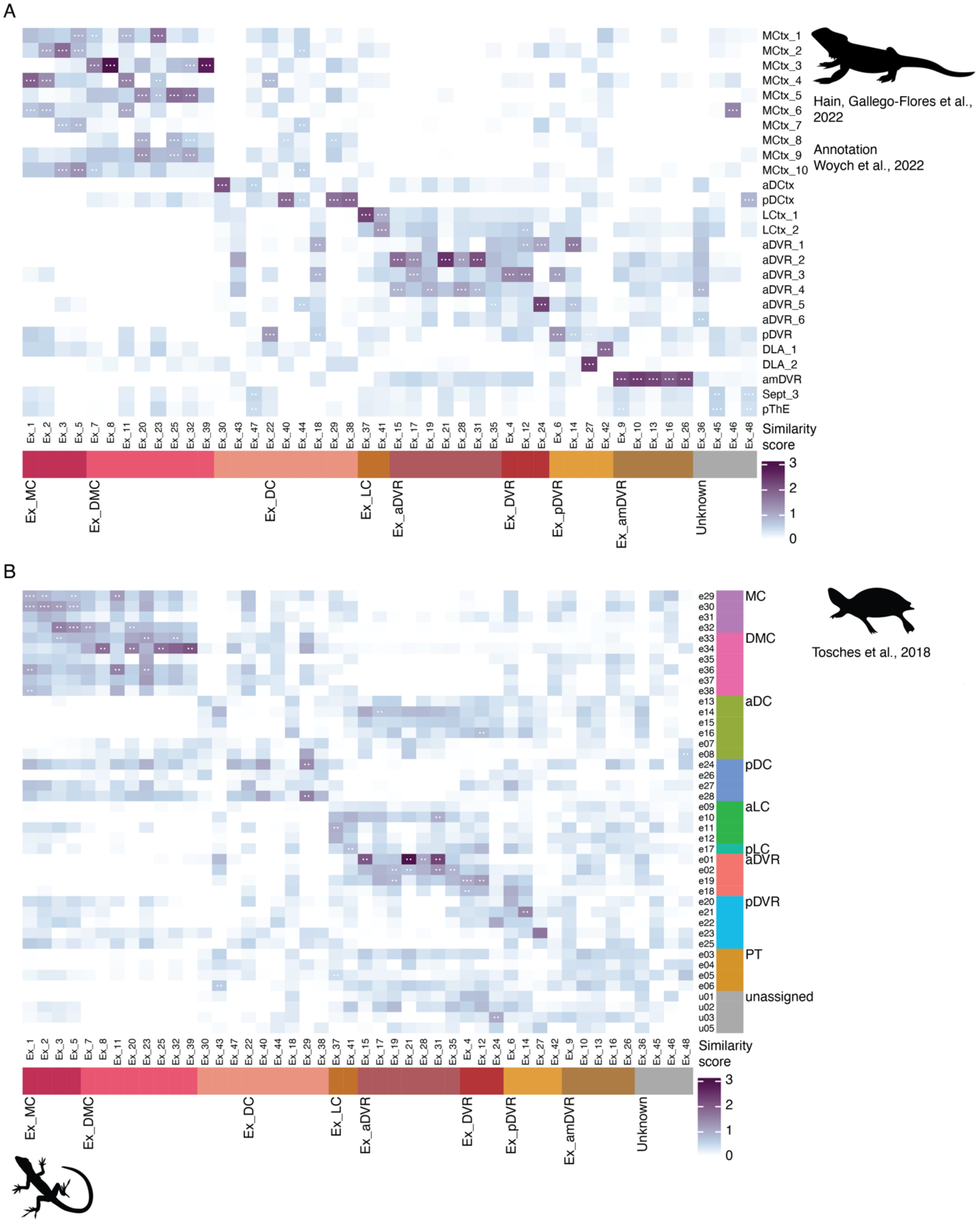


**Fig. S4. Comparison of excitatory neurons between lizard and available reptilian datasets**
Comparison between *Anolis carolinensis* excitatory clusters and *Pogona vitticeps* single cell data from (*12*), annotated by (*50*) (A) and turtle data from (*7*) (B) using three methods. Scores were scaled between 0 and 1 per method and summed across all methods to represent the similarity score. White dots in tiles are shown when populations are among the top reciprocal matches according to two or all three methods. Abbreviations Pogona: MCtx, medial cortex; aDCtx, anterior dorsal cortex; pDCtx,

posterior dorsal cortex; LCtx, lateral cortex; aDVR, anterior DVR; pDVR, posterior DVR; DLA, dorsal lateral amygdala; amDVR, anterior medial DVR; Sept, septum; pThE, pre-thalamic eminence. Abbreviations Anolis: MC, medial cortex; DMC, dorsal medial cortex; DC, dorsal cortex; LC, lateral cortex. Abbreviations turtle: PT, pallial thickening.

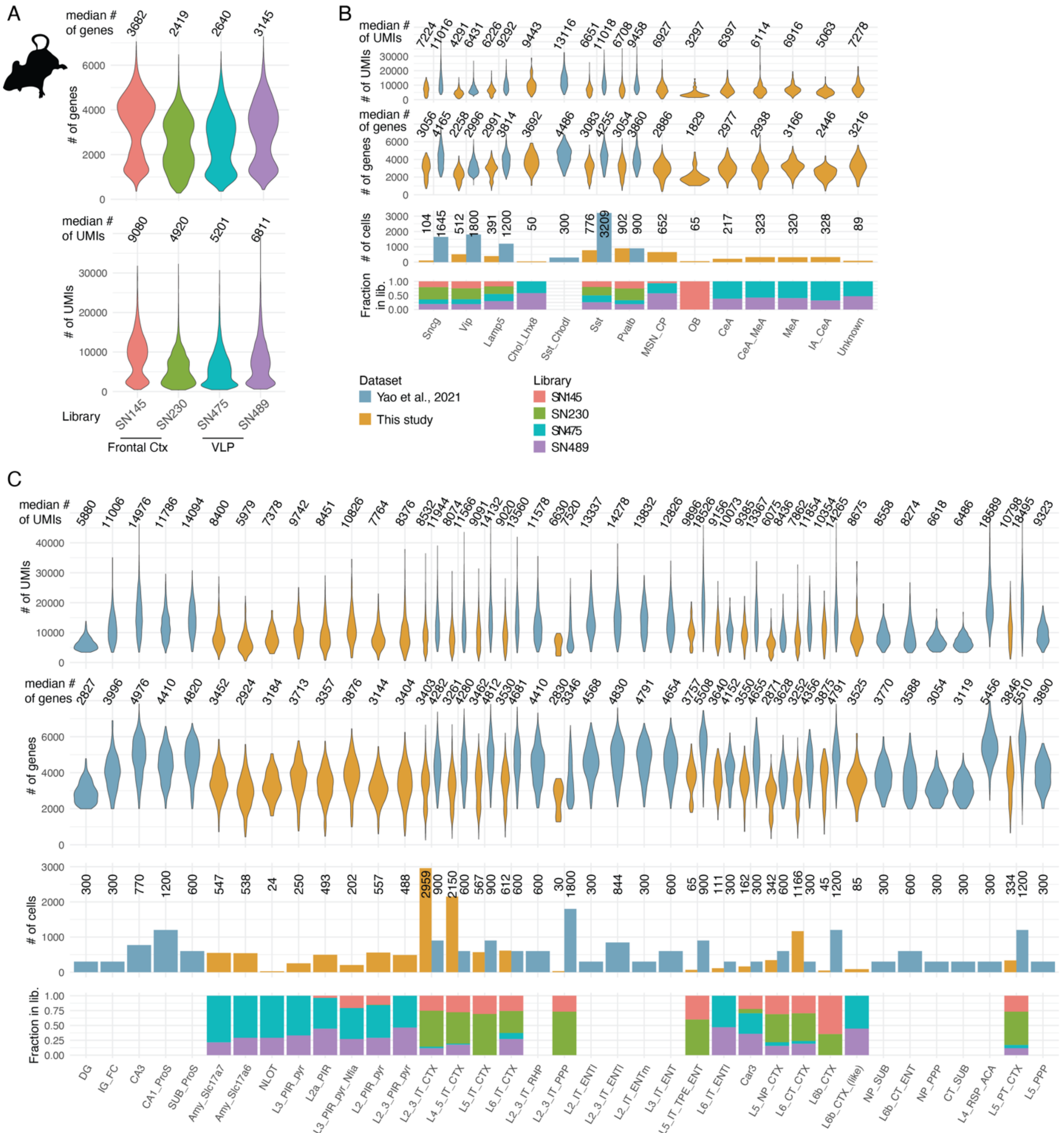


**Fig. S5. Adult mouse pallium dataset**

(A) Number of UMI counts (bottom) and detected genes (top) per snRNA-seq library in our dataset after selection of high quality cells colored by library (each library sampled from a different individual). Ctx, cortex; VLP, ventral and lateral pallia derived structures. Number of UMI counts and detected genes (top), number of cells (mid) per datasets and fraction of cells in our dataset stemming from different snRNA-seq libraries/individuals per inhibitory subclass (B) and excitatory subclass (C). For abbreviations see Table S3.

A

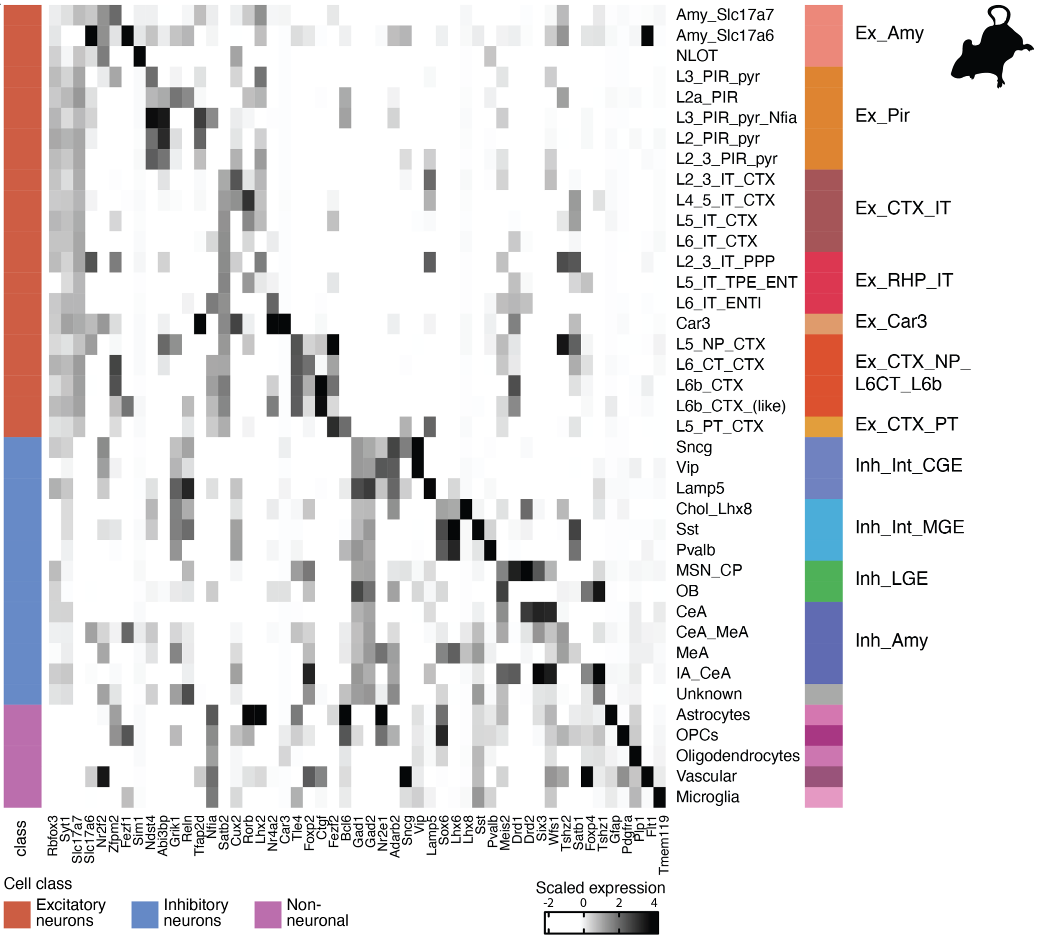


**Fig. S6. Adult mouse pallium marker gene expression**

(A) Heatmap of selected marker gene expression across subclasses in the mouse pallium atlas. Coloured bar and text on the right represents broad neighborhood labels adapted from (*14*). For abbreviations see Table S3.

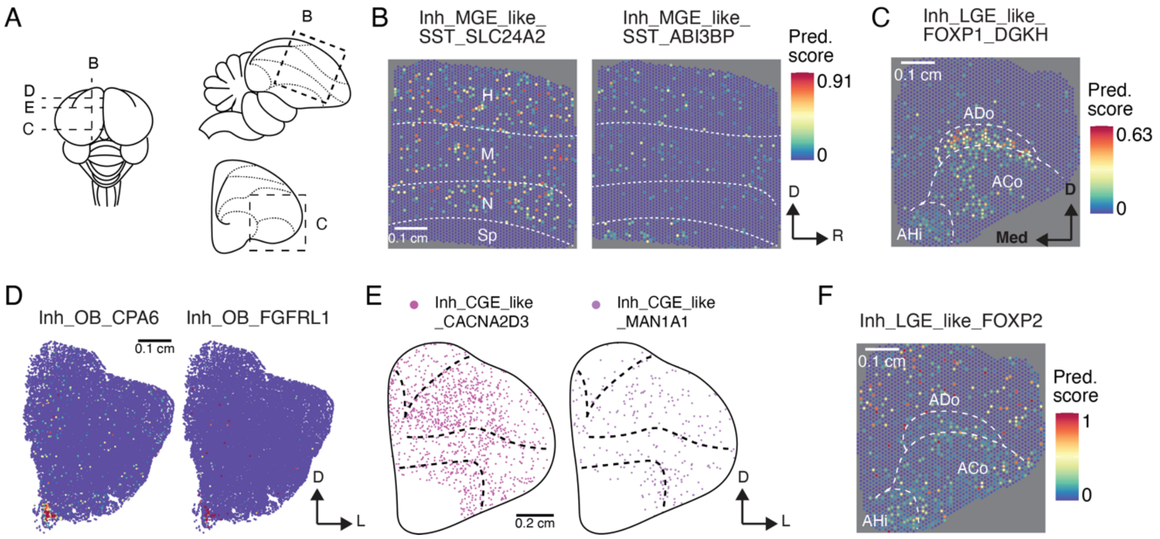


**Fig. S7. Spatial distribution of inhibitory neurons in the adult chicken pallium**
(A) Schematics of chicken brain from the top (left), in a sagittal section (top right) and posterior coronal section (bottom right) to illustrate positions of tissue sections shown in (B-E). Dotted lines represent borders between pallial brain regions. (B) Spatial location of MGE-derived supertypes according to Visium. High prediction (Pred.) scores indicate high probability that cells with the respective identity were present within the spot's area. H, Hyperpallium; M, Mesopallium; N, Nidopallium; Sp, subpallium. (C) Spatial location of LGE-derived supertype. ADo, amygdala dorsal region; ACo, arcopallial core nuclei; AHi, amygdalo-hippocampal region. (D) Spatial location probabilities of two LGE-derived supertypes according to *in situ* sequencing (ISS). (E) Spatial location of CGE-derived supertypes according to ISS. Only segmented cells with confidently assigned identity are shown. (F) Spatial location of LGE-derived supertype Inh_LGE_like FOXP2 according to Visium.

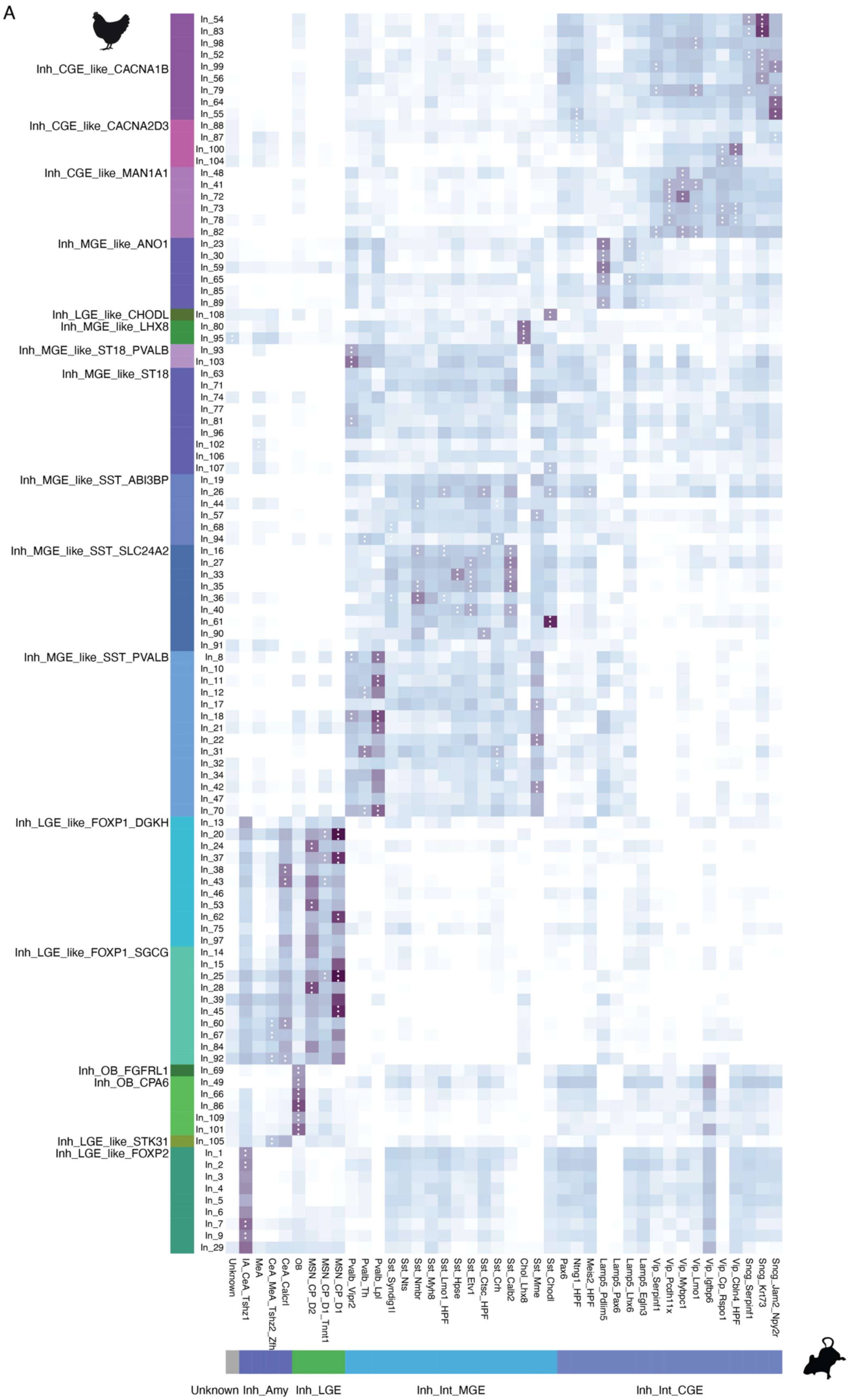

A
Inh_CGE_like_CACNA1B
Inh_CGE_like_CACNA2D3
Inh_CGE_like_MAN1A1
Inh_MGE_like_ANO1
Inh_LGE_like_CHODL
Inh_MGE_like_LHX8
Inh_MGE_like_ST18_PVALB
Inh_MGE_like_ST18
Inh_MGE_like_SST_ABI3BP
Inh_MGE_like_SST_SLC24A2
Inh_MGE_like_SST_PVALB
Inh_LGE_like_FOXP1_DGKH
Inh_LGE_like_FOXP1_SGCG
Inh_OB_FGFRL1
Inh_OB_CPA6
Inh_LGE_like_STK31
Inh_LGE_like_FOXP2
Unknown
Inh_Amy
Inh_LGE
Inh_Int_MGE
Inh_Int_CGE

**Fig. S8. Cluster level comparison of inhibitory neurons between chicken and mouse pallium**

(A) Comparison between chicken clusters and mouse inhibitory supertypes based on three different methods. Scores were scaled between 0 and 1 per method and summed across all methods to represent the similarity score. White dots in tiles are shown when populations are among the top reciprocal matches according to two or all three methods. Amy, amygdala; MSN; medium spiny neuron; CP, caudate putamen; OB, olfactory bulb; CeA, central amygdala; MeA, medial amygdala; IA, intercalated amygdala.

A

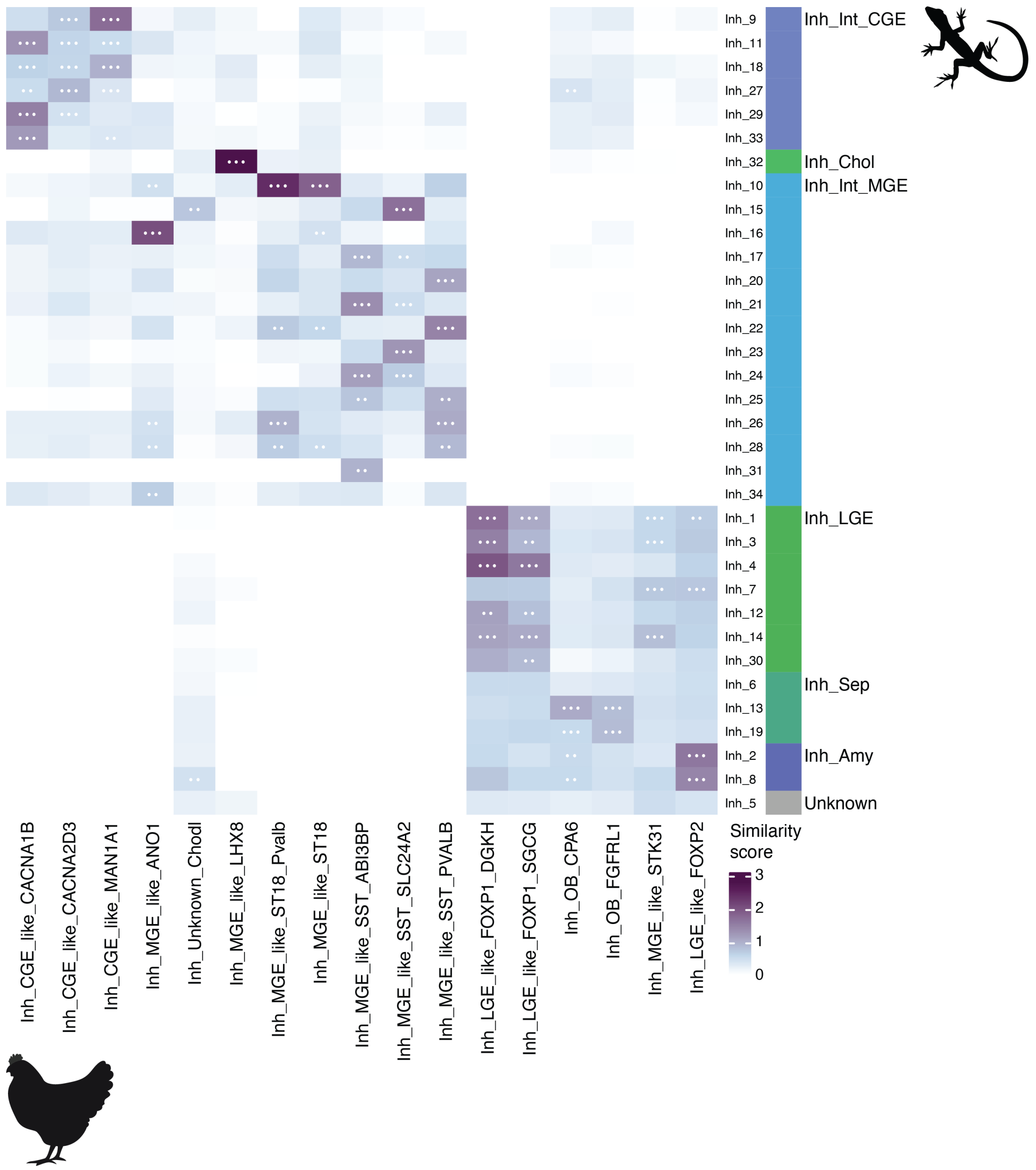


**Fig. S9. Comparison of inhibitory neurons between chicken and lizard pallium**

(A) Comparison between inhibitory supertypes in the chicken pallium to clusters of inhibitory neurons in the lizard pallium based on three methods. Scores were scaled between 0 and 1 per method and summed across all methods to represent the similarity score. White dots in tiles are shown when populations are among the top reciprocal matches according to two or all three methods. CGE, caudal ganglionic eminence; MGE, medial ganglionic eminence; LGE, lateral ganglionic eminence;Chol, cholinergic; Sep, septum; Amy, amygdala.

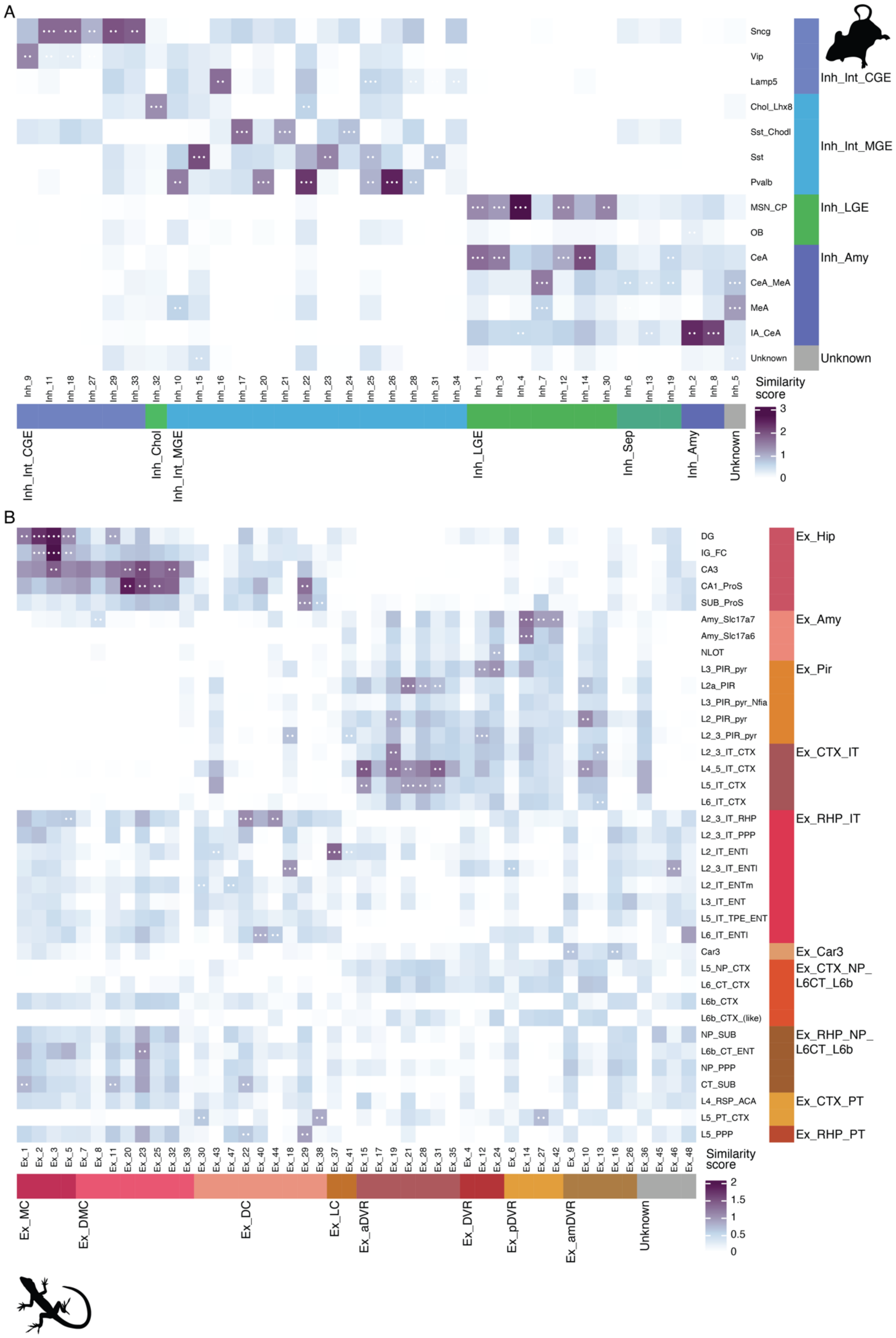
A
Sncg
Vip
Lamp5
Chol_Lhx8
Sst_Chodl
Sst
Pvalb
MSN_CP
OB
CeA
CeA_MeA
MeA
IA_CeA
Unknown
Inh_Int_CGE
Inh_Int_MGE
Inh_LGE
Inh_Amy
Unknown
Inh_9
Inh_11
Inh_18
Inh_27
Inh_29
Inh_33
Inh_32
Inh_10
Inh_15
Inh_16
Inh_17
Inh_20
Inh_21
Inh_22
Inh_23
Inh_24
Inh_25
Inh_26
Inh_28
Inh_31
Inh_34
Inh_1
Inh_3
Inh_4
Inh_7
Inh_12
Inh_14
Inh_30
Inh_6
Inh_13
Inh_19
Inh_2
Inh_8
Inh_5
Similarity score
3
2
1
0
Inh_Int_CGE
Inh_Chol
Inh_Int_MGE
Inh_LGE
Inh_Sep
Inh_Amy
Unknown
B
DG
IG_FC
CA3
CA1_ProS
SUB_ProS
Amy_Slc17a7
Amy_Slc17a6
NLOT
L3_PIR_pyr
L2a_PIR
L3_PIR_pyr_Nfia
L2_PIR_pyr
L2_3_PIR_pyr
L2_3_IT_CTX
L4_5_IT_CTX
L5_IT_CTX
L6_IT_CTX
L2_3_IT_RHP
L2_3_IT_PPP
L2_IT_ENTl
L2_3_IT_ENTl
L2_IT_ENTm
L3_IT_ENT
L5_IT_TPE_ENT
L6_IT_ENTl
Car3
L5_NP_CTX
L6_CT_CTX
L6b_CTX
L6b_CTX_(like)
NP_SUB
L6b_CT_ENT
NP_PPP
CT_SUB
L4_RSP_ACA
L5_PT_CTX
L5_PPP
Ex_Hip
Ex_Amy
Ex_Pir
Ex_CTX_IT
Ex_RHP_IT
Ex_Car3
Ex_CTX_NP_
L6CT_L6b
Ex_RHP_NP_
L6CT_L6b
Ex_CTX_PT
Ex_RHP_PT
Ex_1
Ex_2
Ex_3
Ex_5
Ex_7
Ex_8
Ex_11
Ex_20
Ex_23
Ex_25
Ex_32
Ex_39
Ex_30
Ex_43
Ex_47
Ex_22
Ex_40
Ex_44
Ex_18
Ex_29
Ex_38
Ex_37
Ex_41
Ex_15
Ex_17
Ex_19
Ex_21
Ex_28
Ex_31
Ex_35
Ex_4
Ex_12
Ex_24
Ex_6
Ex_14
Ex_27
Ex_42
Ex_9
Ex_10
Ex_13
Ex_16
Ex_26
Ex_36
Ex_45
Ex_46
Ex_48
Similarity score
2
1.5
1
0.5
0
Ex_MC
Ex_DMC
Ex_DC
Ex_LC
Ex_aDVR
Ex_DVR
Ex_pDVR
Ex_amDVR
Unknown

**Fig. S10. Comparison of neurons between mouse and lizard pallium**

Comparison between inhibitory (A) and excitatory (B) mouse subclasses and lizard clusters based on three methods. Scores were scaled between 0 and 1 per method and summed across all methods to represent the similarity score. White dots in tiles are shown when populations are among the top reciprocal matches according to two or all three methods. CGE, caudal ganglionic eminence; MGE, medial ganglionic eminence; LGE, lateral ganglionic eminence;Chol, cholinergic; Sep, septum; Amy, amygdala; MC, medial cortex; DMC, dorsal medial cortex; DC, dorsal cortex; LC, lateral cortex; aDVR, anterior DVR; pDVR, posterior DVR; amDVR, anterior medial DVR. For mouse abbreviations see Table S3.

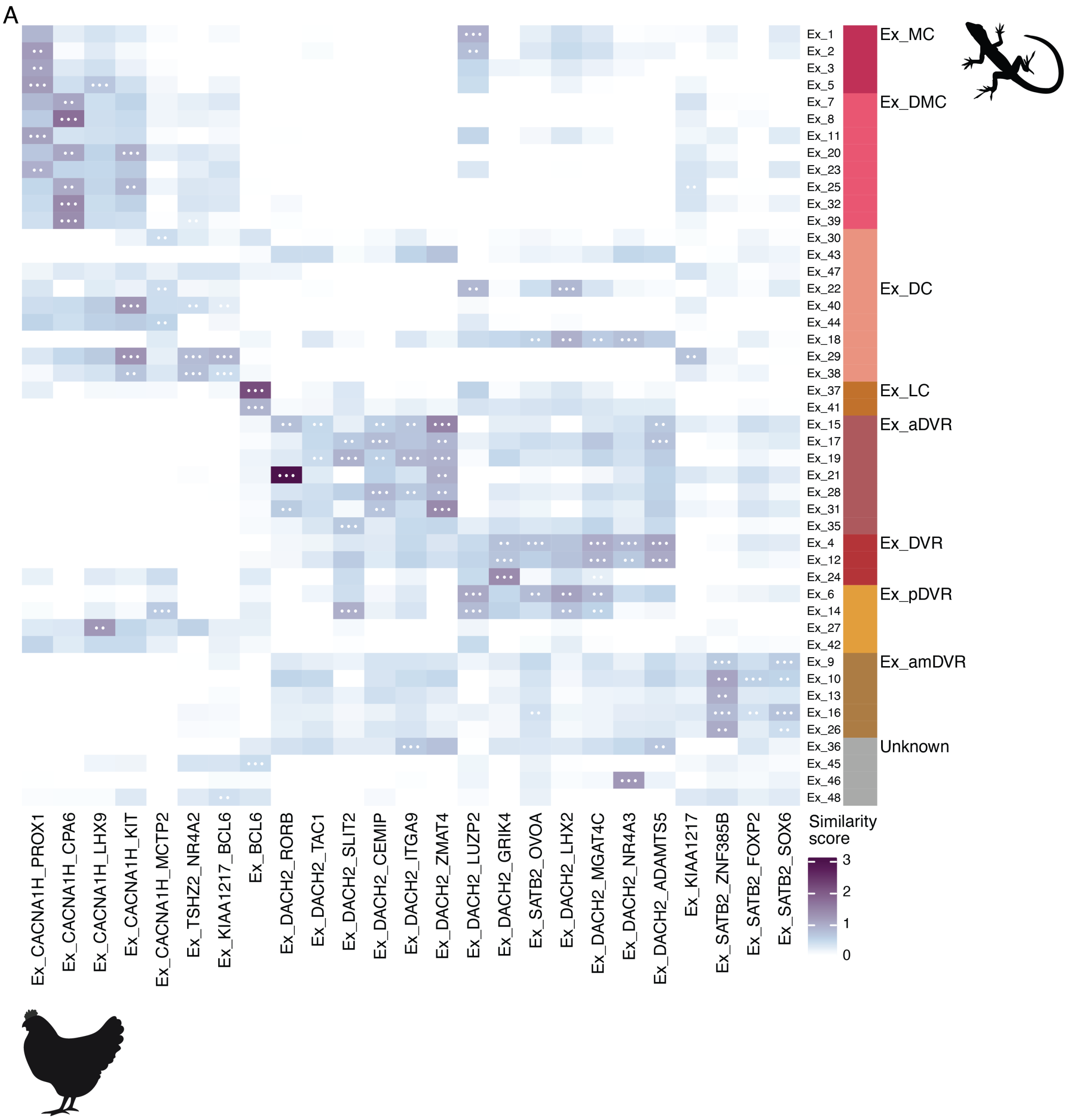


**Fig. S11. Comparison of excitatory neurons between chicken and lizard pallium**

(A) Comparison between excitatory supertypes in the chicken pallium to clusters of excitatory neurons in the lizard pallium based on three methods. Scores were scaled between 0 and 1 per method and summed across all methods to represent the similarity score. White dots in tiles are shown when populations are among the top reciprocal matches according to two or all three methods. MC, medial cortex; DMC, dorsal medial cortex; DC, dorsal cortex; LC, lateral cortex; aDVR, anterior DVR; pDVR, posterior DVR; amDVR, anterior medial DVR.

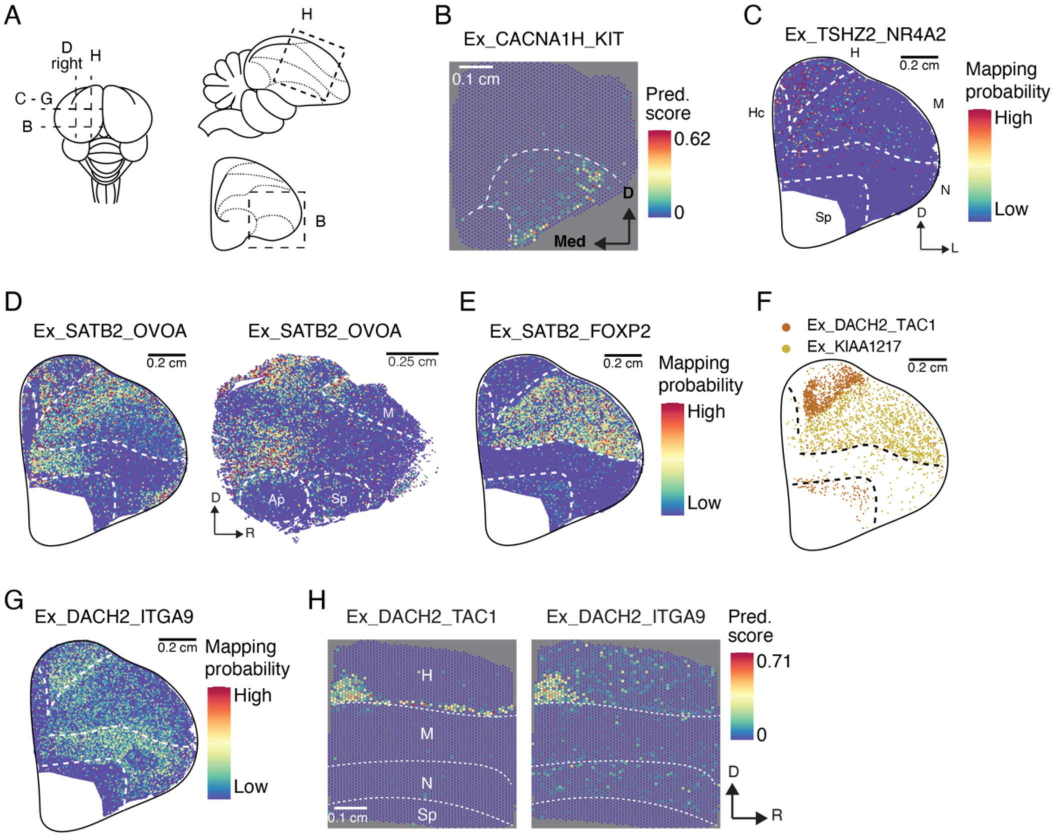


**Fig. S12. Spatial distribution of excitatory neurons in the adult chicken pallium**
(A) Schematics of chicken brain from the top (left), in a sagittal section (top right) and posterior coronal section (bottom right) to illustrate positions of tissue sections shown in (B-H). Dotted lines represent borders between pallial brain regions. (B) Spatial location of medial pallial supertype according to Visium. High prediction (Pred.) scores indicate high probability that cells with the respective identity were present within the spot's area. Med, medial; D, dorsal. (C) Spatial location probabilities of hyperpallial supertype according to *in situ* sequencing (ISS). Hc, hippocampal areas; H, Hyperpallium; M, Mesopallium; N, Nidopallium; Sp, subpallium. (D) Spatial location probabilities of supertype Ex_SATB2_OVOA according to ISS. M (E) Spatial location of a mesopallial supertype according to Visium. R, rostral. (F) Spatial location of a hyperpallial and mesopallial supertype according to ISS. Only segmented cells with confidently assigned identity are shown. (G) Spatial location probabilities of supertype Ex_DACH2_ITGA9 according to ISS. (H) Spatial location of hyperpallial supertype Ex_DACH2_TAC1 and supertypes Ex_DACH2_ITGA9 according to Visium.

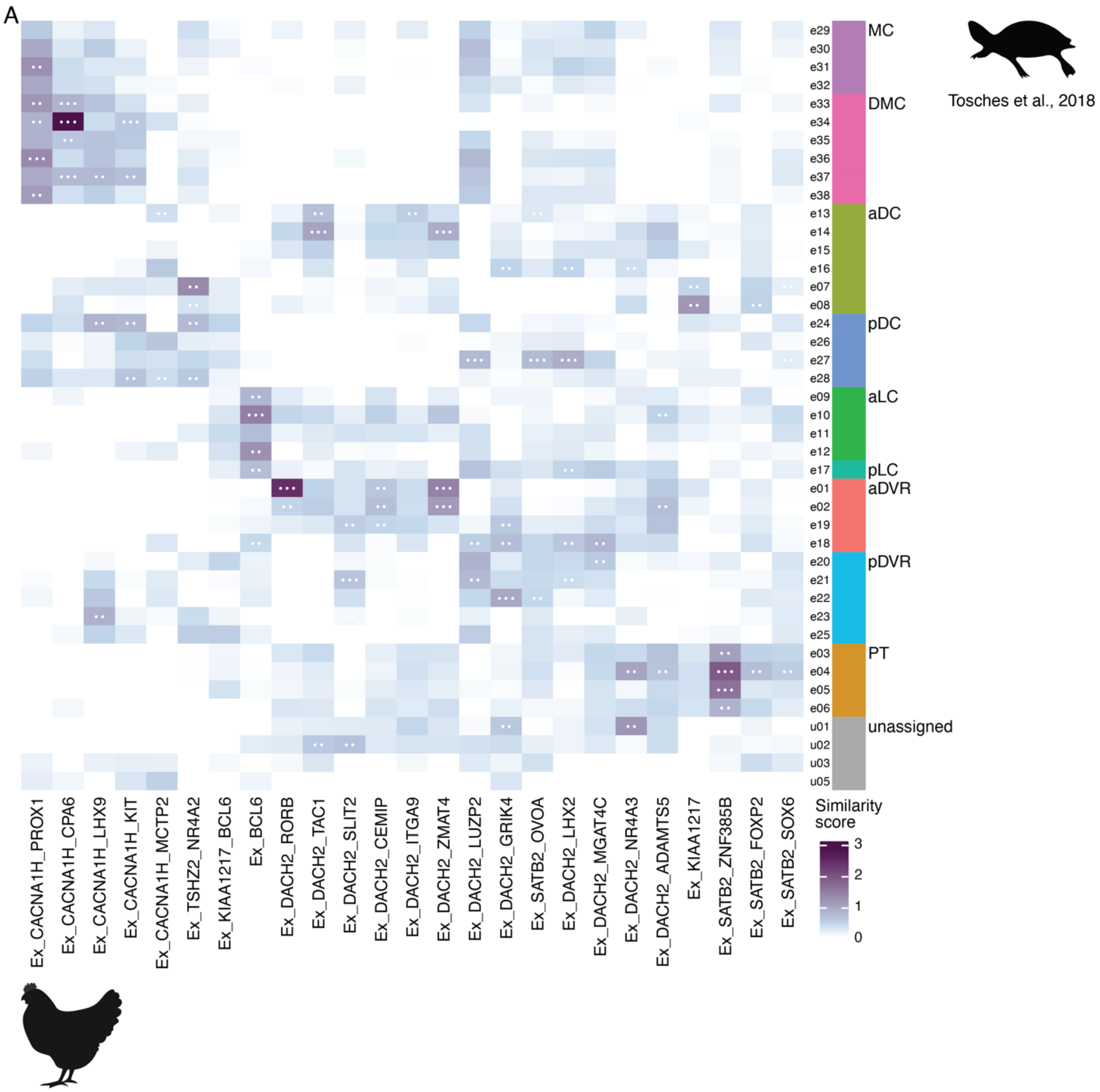


**Fig. S13. Comparison of excitatory neurons between chicken and turtle pallium**

(A) Comparison between excitatory supertypes in the chicken pallium to clusters of excitatory neurons in the turtle pallium from (*7*) based on three methods. Scores were scaled between 0 and 1 per method and summed across all methods to represent the similarity score. White dots in tiles are shown when populations are among the top reciprocal matches according to two or all three methods. MC, medial cortex; DMC, dorsal medial cortex; aDC, anterior dorsal cortex; pDC, posterior dorsal cortex; aLC, anterior lateral cortex; pLC, posterior lateral cortex; aDVR, anterior DVR; pDVR, posterior DVR; PT, pallial thickening.

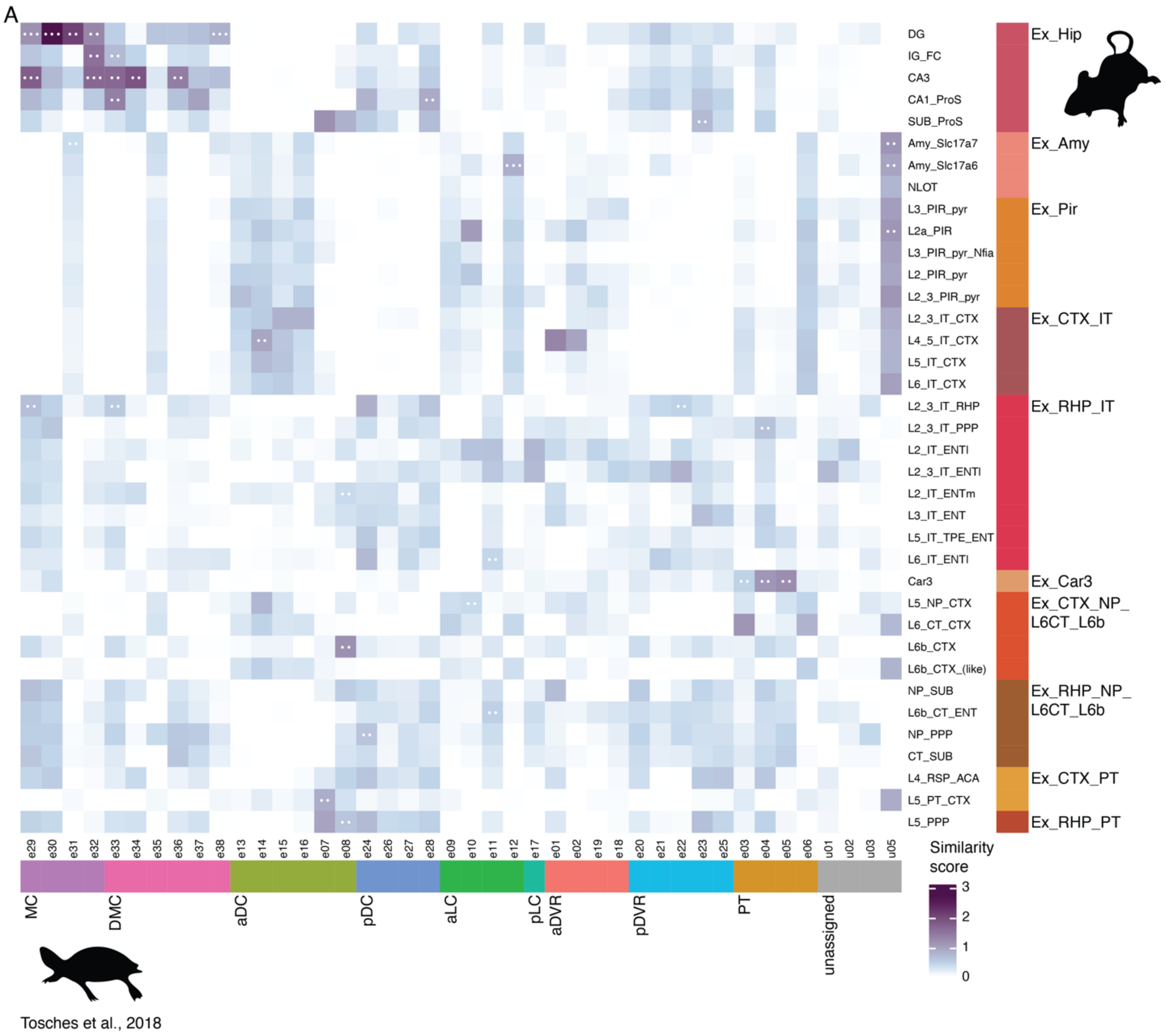


**Fig. S14. Comparison of excitatory neurons between mouse and turtle pallium**

(A) Comparison between excitatory subclasses in the mouse pallium to clusters of excitatory neurons in the turtle pallium from (*7*) based on three methods. Scores were scaled between 0 and 1 per method and summed across all methods to represent the similarity score. White dots in tiles are shown when populations are among the top reciprocal matches according to two or all three methods. MC, medial cortex; DMC, dorsal medial cortex; aDC, anterior dorsal cortex; pDC, posterior dorsal cortex; aLC, anterior lateral cortex; pLC, posterior lateral cortex; aDVR, anterior DVR; pDVR, posterior DVR; PT, pallial thickening. For mouse abbreviations see Table S3.

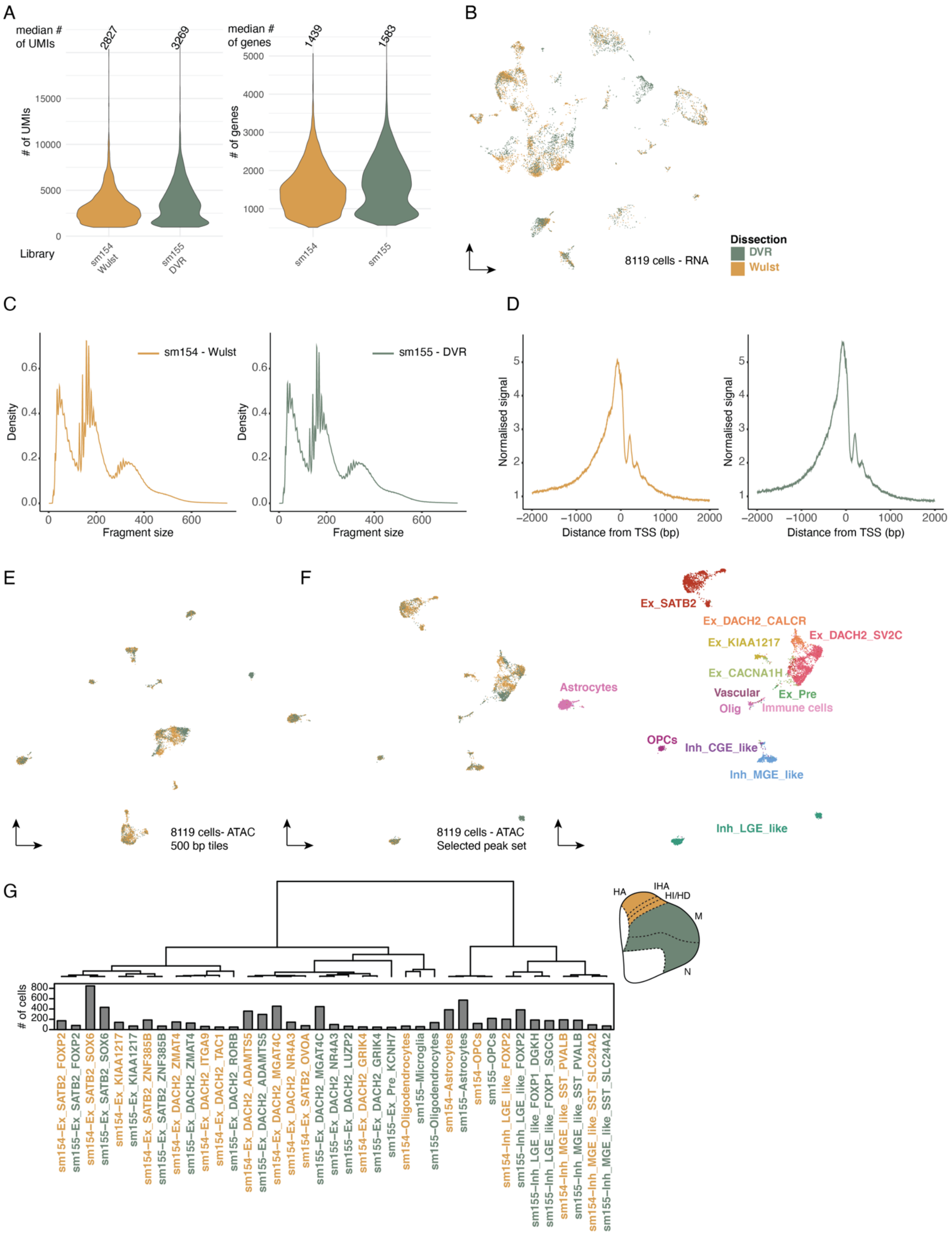


**Fig. S15. snRNA and ATAC multiome of the chicken pallium**
(A) Number of UMI counts (left) and detected genes (right) per snRNA-seq library after selection of high quality cells colored by library/sampled region. (B) UMAP of snRNA-seq data colored by library/dissection. (C) Fragment size distribution of ATAC

libraries. (D) Transcription start site (TSS) enrichment scores of the snATAC-seq libraries. (E) UMAP of snATAC data, based on accessibility across 500bp tiles colored by library/sampled region. (F) UMAP of snATAC data, based on accessibility in selected peak set colored by library/sampled region (left) and assigned subclass identity (right). (G) Correlation dendrogram of pseudobulk accessibility profiles per supertype and library/region (at least 40 cells). Schematic top right: illustration of borders between dissections. HA, apical hyperpallium; IHA, intercalated hyperpallium; HI/HD, intermediate/densocellular hyperpallium; M, mesopallium; N, nidopallium.

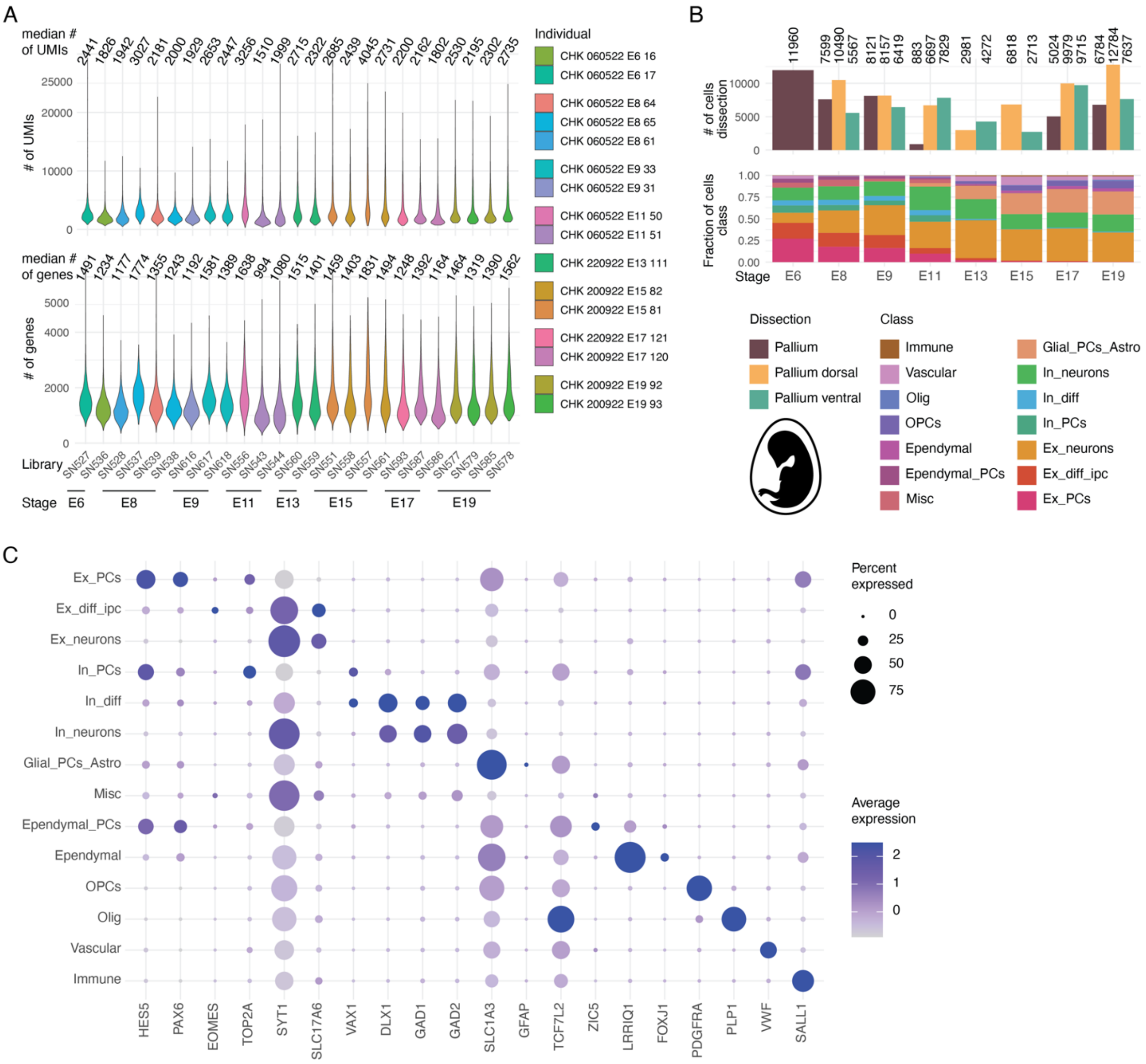

**Fig. S16. Developing chicken pallium single nucleus RNA-sequencing dataset**
(A) Number of UMI counts (top) and detected genes (bottom) per snRNA-seq library after selection of high quality cells colored by sampled individuals. (B) Number of cells per developmental stage and dissected region (top) and fraction of cells per developmental stage belonging to different cell populations (bottom). E, embryonic day. (C) Gene expression dotplot of selected marker genes across cell populations. PCs, progenitor cells; diff, differentiating; ipc, intermediate progenitor cell; Misc, miscellaneous; Olig, Oligodendrocytes.

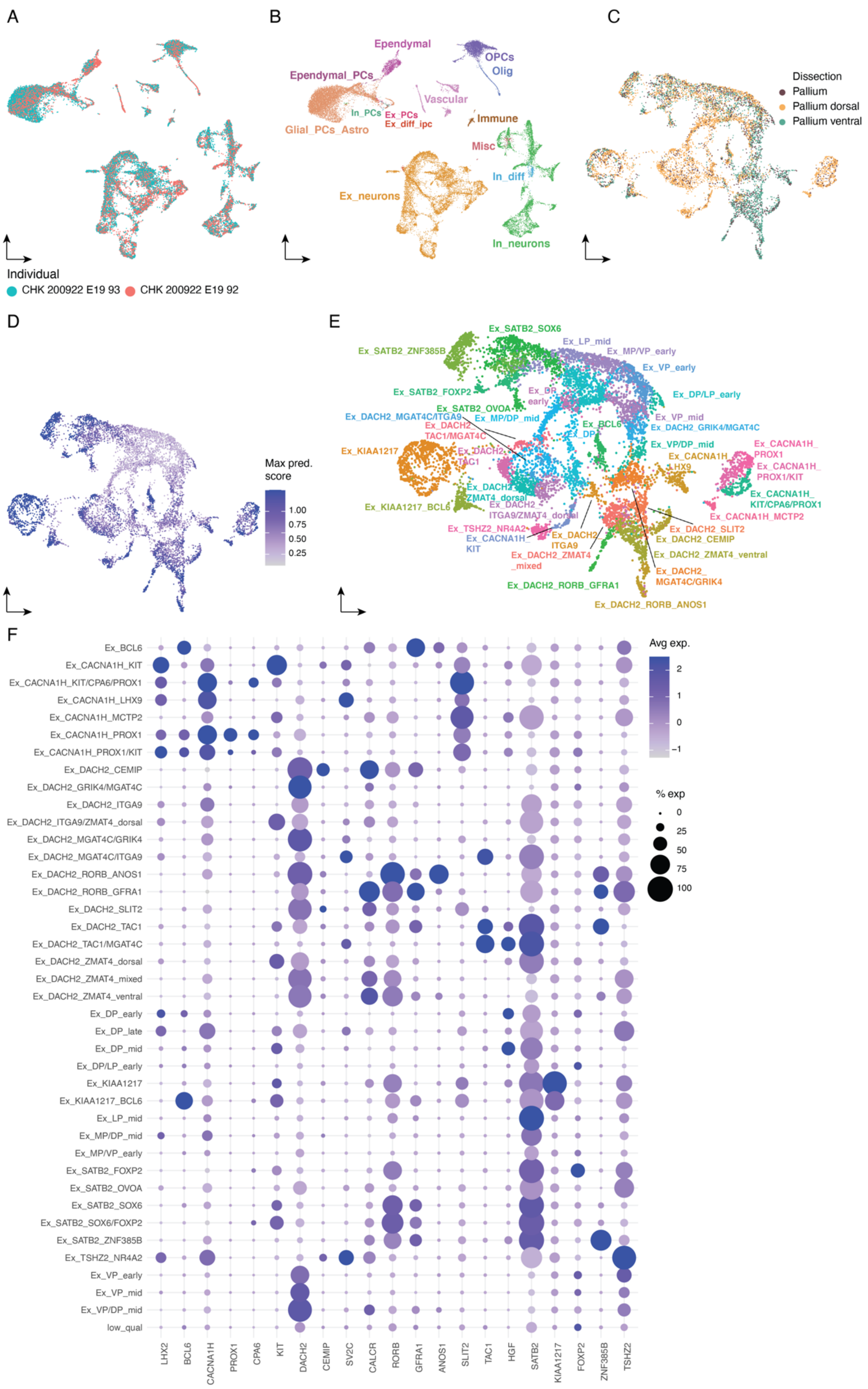

A
B
C
D
E
F
Individual
CHK 200922 E19 93
CHK 200922 E19 92
Ependymal
OPCs
Olig
Ependymal_PCs
Vascular
In_PCs
Ex_PCs
Ex_diff_ipc
Immune
Glial_PCs_Astro
Misc
In_diff
Ex_neurons
In_neurons
Dissection
Pallium
Pallium dorsal
Pallium ventral
Max pred. score
Avg exp.
% exp

**Fig. S17. Embryonic day 19 chicken pallium dataset**

UMAP of E19 snRNA-seq data colored by sampled individual (A), colored by cell class (B) or colored by dissected region (C). UMAP of excitatory neurons (Ex_neurons) colored by maximum prediction score for any adult excitatory supertype after CCA label transfer (D) and colored by and labeled with an annotation, which was used to map cell populations to the tissue in E19 Visium (Fig. 5D, fig. S18). (F) Gene expression dotplot of selected marker genes across annotated cell populations.

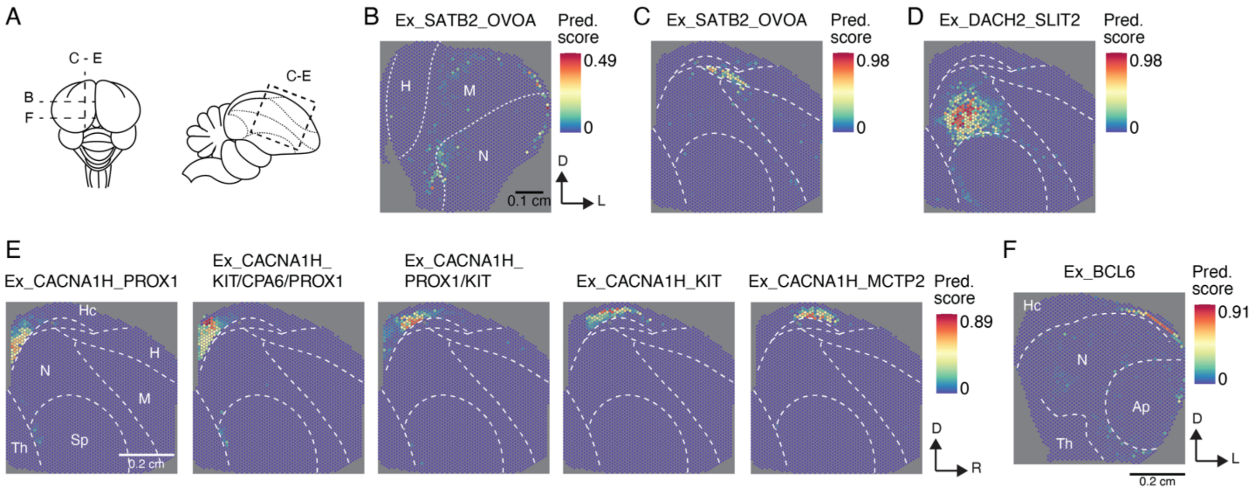


**Fig. S18. Spatial distribution of excitatory neurons in the developing (E19) chicken pallium**
(A) Schematics of E19 chicken brain from the top (left) or in a sagittal section (right) to illustrate positions of tissue sections shown in (B-F). Dotted lines represent borders between pallial brain regions. (B) Spatial location of supertype Ex_SATB2_OVOA in a coronal (B) and sagittal (C) section according to Visium. High prediction (Pred.) scores indicate high probability that cells with the respective identity were present within the spot's area. D, dorsal; L, lateral; H, hyperpallium; M, mesopallium; N, Nidopallium. (D) Spatial location of supertype Ex_DACH2_SLIT2 according to Visium. (E) Spatial location of hippocampal/medial pallial cell populations according to Visium. Hc, hippocampal areas; Sp subpallium; Th, Thalamus; R, rostral. (F) Spatial location of Ex_BCL6 population above the ventricle according to Visium. Ap, Arcopallium.

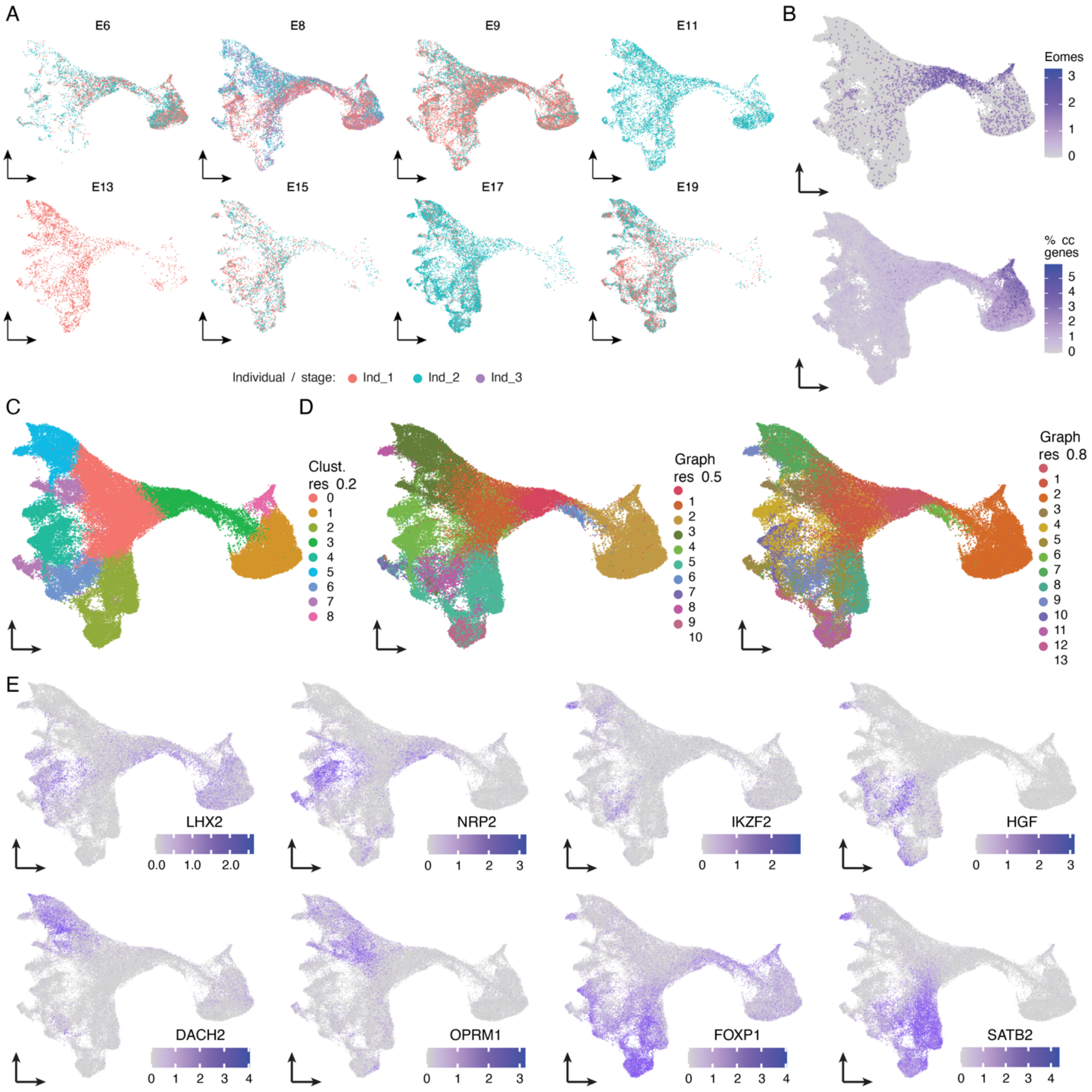


**Fig. S19. Lineages of excitatory neurons in the developing chicken pallium**
(A) UMAP of excitatory neuron lineage in the developing chicken pallium split by developmental stage and colored by sampled individual per stage. (B) UMAP of excitatory neuron lineage colored by expression of *Eomes* (top) and percentage of UMIs stemming from cell cycle (cc) related genes (bottom). UMAP of excitatory neuron lineage colored by low resolution clustering according to standard clustering procedure (C) or by clusters resulting from clustering of a weighted graph at two different resolutions, which was constructed based on the clustering and integration of individual developmental stages (D). (E) UMAP colored by expression of selected marker genes of excitatory neuron sublineages.

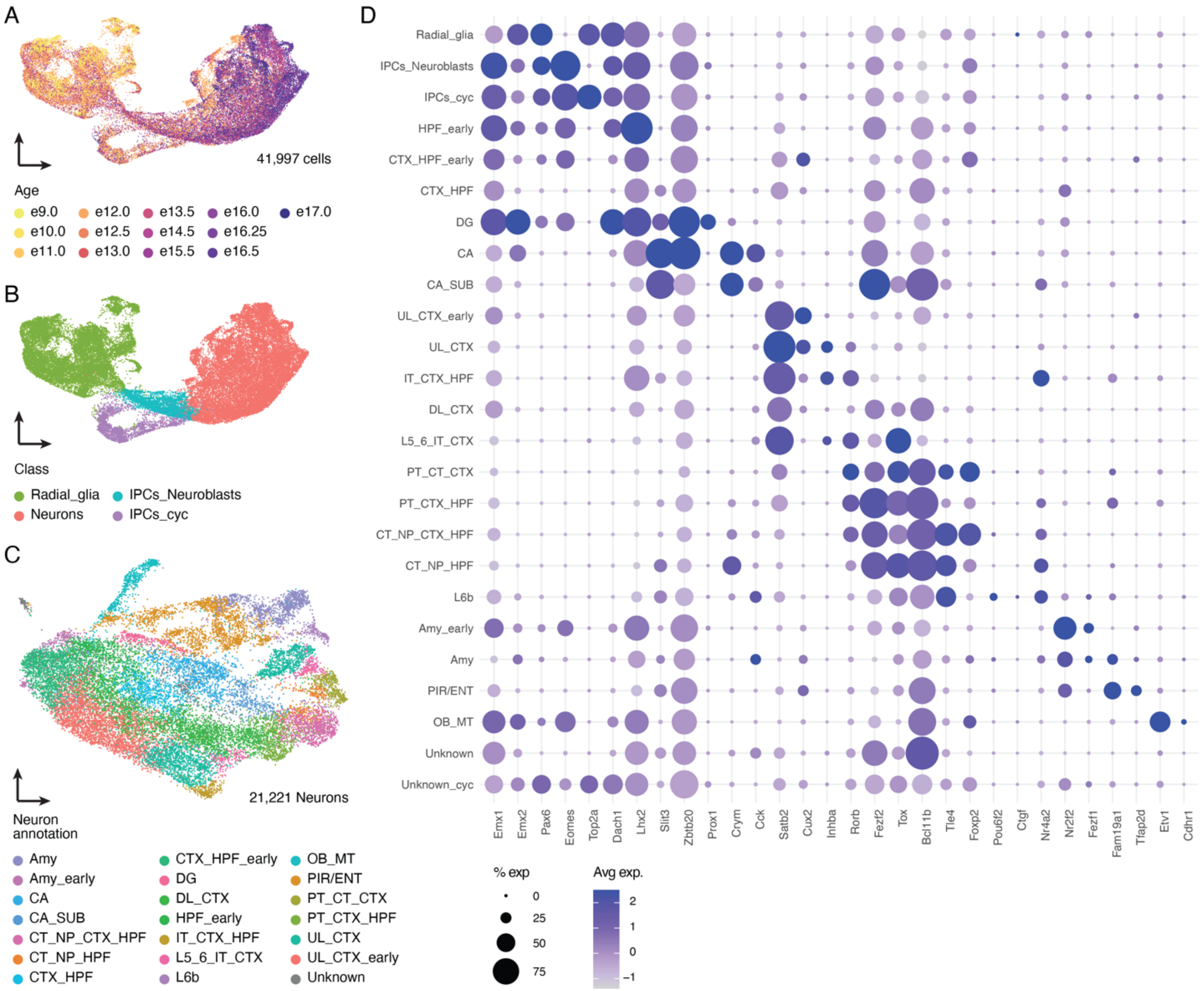


**Fig. S20. Excitatory neurons in the developing mouse pallium**
UMAP of the pallial excitatory neuron lineage subset from (*47*) colored by Age (A) and cell class (B). IPCs, intermediate progenitor cells; cyc, cycling. (C) UMAP of excitatory neurons subset from whole excitatory lineage colored by annotation, which was used to compare cell populations between the embryonic mouse and chicken pallium in Fig. 6H. Amy, amygdala; CA, cornu ammonis; SUB, subiculum; CT, cortico-thalamic; NP, near-projecting; HPF, hippocampal formation; CTX, cortex; DG, dentate gyrus; DL, deep layer; IT, intra-telencephalic; OB-MT, olfactory bulb mitral tufted; PIR/ENT, piriform/entorhinal cortex; PT, pyramidal tract projecting; UL, upper layer.

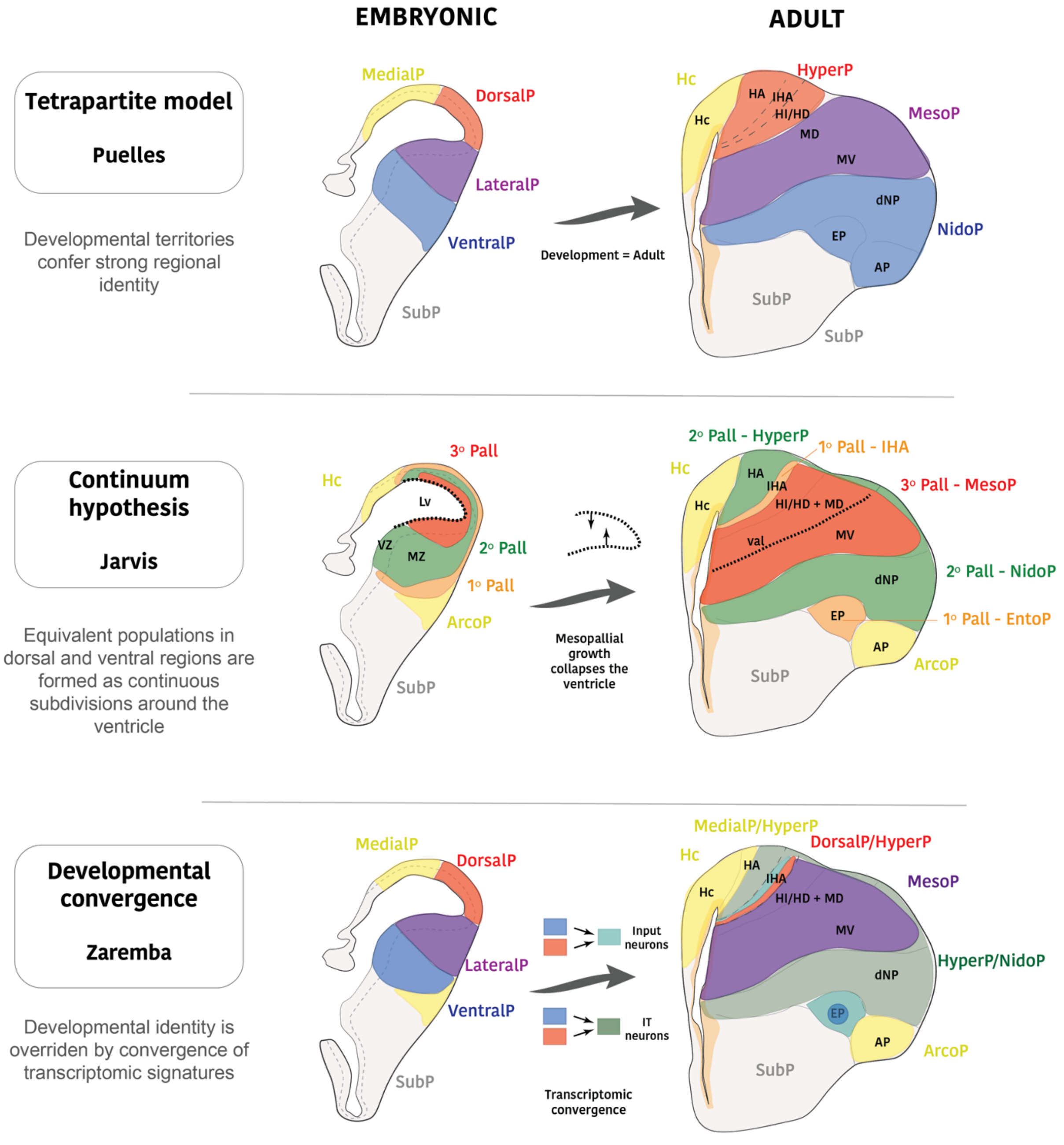


**Fig. S21. An updated model of the chicken pallium**

Illustration of our updated model of chicken pallium development (bottom) in comparison to previous models according to (*6*) (top) and (*34*) (mid).P, pallium; Hc, hippocampal areas; IHA, intercalated hyperpallium; HI/HD, intermediate/densocellular hyperpallium; MD, dorsal mesopallium; MV, ventral mesopallium; dNP, dorsal nidopallium; Ep, entopallium (sensory input area of the DVR); AP, Arcopallium; IT, intra-telencephalic.

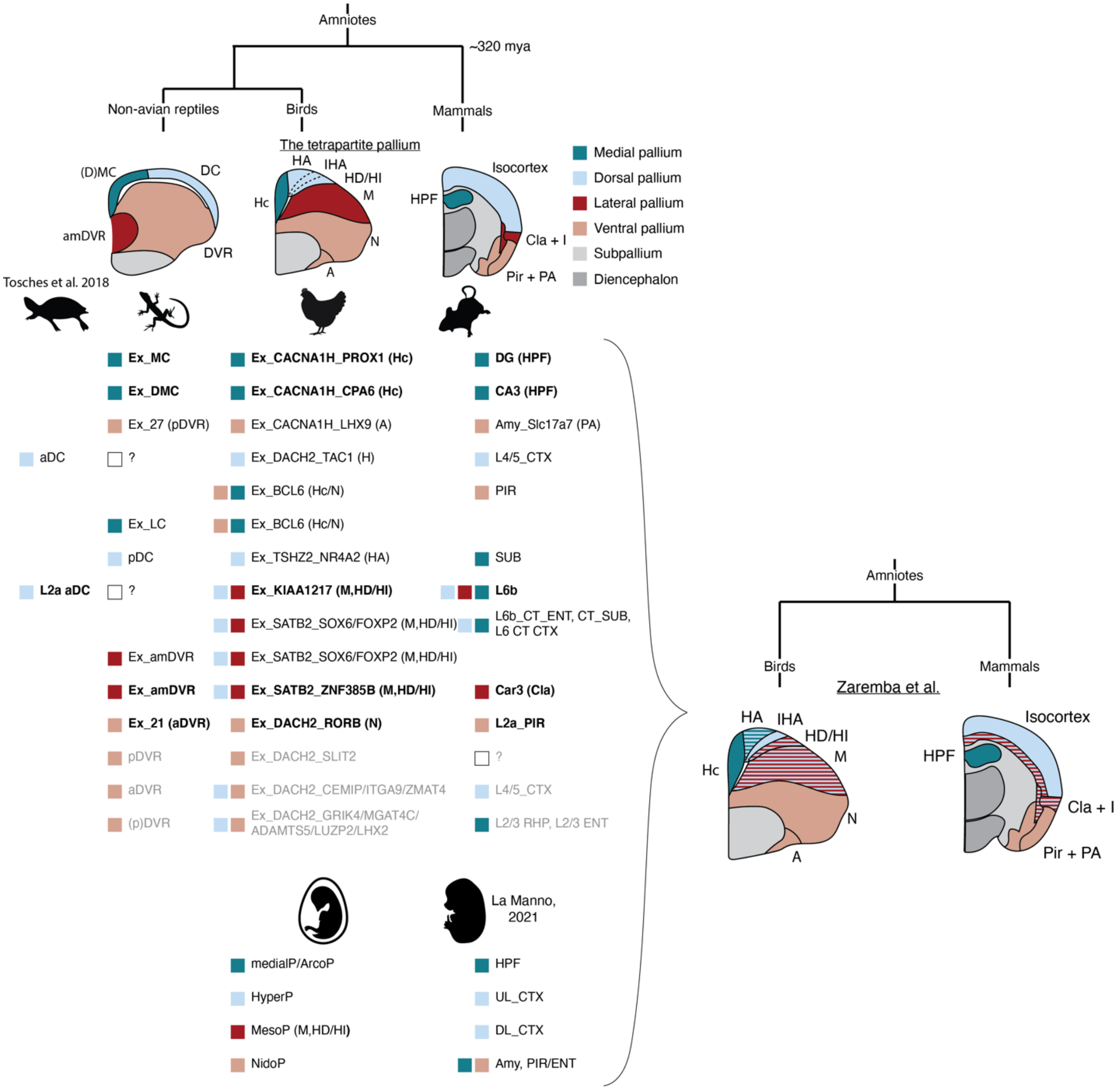


**Fig. S22. Homology relationships of cell types and regions**

(Top) Schematic representation of coronal sections of the telencephalon in lizard (left), chicken (middle), and mouse (right). Brightly colored areas represent the pallium divided into developmental, homologous sectors according to the tetrapartite pallium model (*6*). Mya, million years ago; DVR, dorsal ventricular ridge; Hc, hippocampus; Cla, claustrum; I, insular cortex; Pir, piriform cortex; PA, pallial amygdala. (Bottom) Identified homologous cell types. Coloured squares indicate the cell type's spatial location in the framework of the tetrapartite pallium model. Bold text indicates homologies assigned with high confidence, grey text indicates low confidence. Cell types in the chicken pallium are listed twice, if the corresponding reptilian or mammalian populations do not correspond based on the separate comparison of lizard/turtle and mammalian cell types. Based on our identified cell type homologies we suggest a new model of evolutionary relationships between different regions of the amniote pallium.

**Table S1.**
SnRNA-seq samples, libraries and dissections.

**Table S2.**
Adult chicken pallium snRNA-seq atlas annotation

**Table S3.**
Abbreviations of pallial structures in the mouse

**Table S4.**
Differentially expressed genes between hyper- and nidopallium

**Table S5.**
List of genes profiled by in situ sequencing

**Table S6.**
Genome assemblies used for OrthoFinder

**Supplementary References**


60. S. R. Krishnaswami, R. V. Grindberg, M. Novotny, P. Venepally, B. Lacar, K. Bhutani, S. B. Linker, S. Pham, J. A. Erwin, J. A. Miller, R. Hodge, J. K. McCarthy, M. Kelder, J. McCorrison, B. D. Aevermann, F. D. Fuertes, R. H. Scheuermann, J. Lee, E. S. Lein, N. Schork, M. J. McConnell, F. H. Gage, R. S. Lasken, Using single nuclei for RNA-seq to capture the transcriptome of postmortem neurons. *Nat. Protoc.* **11**, 499–524 (2016).
61. Z. Y. Wang, E. Leushkin, A. Liechti, S. Ovchinnikova, K. Mößinger, T. Brüning, C. Rummel, F. Grützner, M. Cardoso-Moreira, P. Janich, D. Gatfield, B. Diagouraga, B. de Massy, M. E. Gill, A. H. F. M. Peters, S. Anders, H. Kaessmann, Transcriptome and translatome co-evolution in mammals. *Nature*, doi: 10.1038/s41586-020-2899-z (2020).
62. A. Dobin, C. A. Davis, F. Schlesinger, J. Drenkow, C. Zaleski, S. Jha, P. Batut, M. Chaisson, T. R. Gingeras, STAR: ultrafast universal RNA-seq aligner. *Bioinformatics* **29**, 15–21 (2013).
63. P.-L. Germain, A. Lun, C. G. Meixide, W. Macnair, M. D. Robinson, Doublet identification in single-cell sequencing data using *scDblFinder*. F1000Research 10:979 [Preprint] (2022). https://doi.org/10.12688/f1000research.73600.2.
64. C. Hafemeister, R. Satija, Normalization and variance stabilization of single-cell RNA-seq data using regularized negative binomial regression. *Genome Biol.* **20**, 296 (2019).
65. I. Korsunsky, N. Millard, J. Fan, K. Slowikowski, F. Zhang, K. Wei, Y. Baglaenko, M. Brenner, P. Loh, S. Raychaudhuri, Fast, sensitive and accurate integration of single-cell data with Harmony. *Nat. Methods* **16**, 1289–1296 (2019).
66. M. Peng, B. Wamsley, A. G. Elkins, D. H. Geschwind, Y. Wei, K. Roeder, Cell type hierarchy reconstruction via reconciliation of multi-resolution cluster tree. *Nucleic Acids Res.* **49**, e91–e91 (2021).
67. R. Suzuki, H. Shimodaira, Pvclust: an R package for assessing the uncertainty in hierarchical clustering. *Bioinformatics* **22**, 1540–1542 (2006).
68. X. Qian, K. D. Harris, T. Hauling, D. Nicoloutsopoulos, A. B. Muñoz-Manchado, N. Skene, J. Hjerling-Leffler, M. Nilsson, Probabilistic cell typing enables fine mapping of closely related cell types in situ. *Nat. Methods* **17**, 101–106 (2020).
69. V. Petukhov, R. J. Xu, R. A. Soldatov, P. Cadinu, K. Khodosevich, J. R. Moffitt, P. V. Kharchenko, Cell segmentation in imaging-based spatial transcriptomics. *Nat. Biotechnol.* **40**, 345–354 (2022).
70. M. I. Love, W. Huber, S. Anders, Moderated estimation of fold change and dispersion for RNA-seq data with DESeq2. *Genome Biol.* **15**, 550 (2014).
71. D. M. Emms, S. Kelly, OrthoFinder: phylogenetic orthology inference for comparative genomics. *Genome Biol.* **20**, 238 (2019).
72. G. Csardi, T. Nepusz, The igraph software package for complex network research. *InterJournal* **Complex Systems**, 1695 (2006).
73. I. Sarropoulos, M. Sepp, R. Frömel, K. Leiss, N. Trost, E. Leushkin, K. Okonechnikov, P. Joshi, P. Giere, L. M. Kutscher, M. Cardoso-Moreira, S. M. Pfister, H. Kaessmann, Developmental and evolutionary dynamics of cis-regulatory elements in mouse cerebellar cells. *Science* **373** (2021).
